# A new mechanism of fibronectin fibril assembly revealed by live imaging and super-resolution microscopy

**DOI:** 10.1101/2020.09.09.290130

**Authors:** Darshika Tomer, Cecilia Arriagada, Sudipto Munshi, Brianna E. Alexander, Brenda French, Pavan Vedula, Valentina Caorsi, Andrew House, Murat Guvendiren, Anna Kashina, Jean E. Schwarzbauer, Sophie Astrof

**Affiliations:** Department of Cell Biology and Molecular Medicine, Cardiovascular Research Institute, Rutgers Biomedical and Health Sciences, 185 South Orange Ave, Newark, NJ, 07103, USA; Sidney Kimmel Medical College of Thomas Jefferson University, Philadelphia, PA, 19107, USA; Multidisciplinary Ph.D. Program in Biomedical Sciences. Cell Biology, Neuroscience and Physiology track, Rutgers Biomedical and Health Sciences, Newark, NJ, 07103, USA; Department of Biomedical Sciences, University of Pennsylvania, Philadelphia, PA 19104, USA; Abbelight, 191 Avenue Aristide Briand, 94230 Cachan, France; Otto H. York Chemical and Materials Engineering, Department of Biomedical Engineering, New Jersey Institute of Technology, Newark, NJ 07102, USA; Department of Molecular Biology, Princeton University, Princeton, New Jersey, USA

**Author notes:** Department of Chemical and Structural Biology, Weizmann Institute of Science, Rehovot, Israel (Sudipto Munshi). Author for correspondence: Sophie Astrof, Ph.D.

## Abstract

Fn1 fibrils have long been viewed as continuous fibers composed of extended, periodically aligned Fn1 molecules. However, our live imaging and single-molecule localization microscopy (SMLM) are inconsistent with this traditional view and show that Fn1 fibrils are composed of roughly spherical nanodomains containing 6-11 Fn1 dimers. As they move toward the cell center, Fn1 nanodomains become organized into linear arrays, wherein nanodomains are spaced at the average periodicity of 105±17 nm. Periodical Fn1 nanodomain arrays are bona fide fibrils: they are resistant to deoxycholate treatment and retain nanodomain periodicity in the absence of cells. The nanodomain periodicity in fibrils remained constant when probed with antibodies recognizing distinct Fn1 epitopes or combinations of antibodies recognizing epitopes spanning the length of Fn1. FUD, a bacterial peptide that binds Fn1 N-terminus and disrupts Fn1 fibrillogenesis does not disrupt the formation of Fn1 nanodomains, instead, it blocks the organization of Fn1 nanodomains into periodical arrays. These studies establish a new paradigm of Fn1 fibrillogenesis.

## Introduction

Fibronectin (Fn1) is a requisite component of the extracellular matrix (ECM) necessary for embryogenesis and homeostasis (Schwarzbauer and DeSimone, 2011). In the absence of Fn1 fibrillogenesis, the binding of Fn1 to cells is not sufficient to regulate key biological processes such as embryonic development, angiogenesis, vascular remodeling, or cartilage condensation (Chiang et al., 2009; Rozario et al., 2009; Singh and Schwarzbauer, 2014; Zhou et al., 2008). Therefore, understanding the mechanisms by which Fn1 proteins assemble into macromolecular fibrils is essential to gain insights into the various functions of Fn1. Fn1 is secreted as a dimer wherein two Fn1 molecules are held in an anti-parallel orientation by two disulfide bonds close to their C-termini (Skorstengaard et al., 1986; Wagner and Hynes, 1979). Fn1 fibrillogenesis is a cell-dependent process (McKeown-Longo and Mosher, 1983), occurs following the binding of Fn1 dimers to cell-surface integrins (Schwarzbauer and DeSimone, 2011). Following integrin binding, intracellular cytoskeletal forces such as actomyosin contractility acting through integrins generate pulling forces on Fn1 dimers, exposing epitopes that promote Fn1 fibrillogenesis (Chernousov et al., 1987; Hocking et al., 1994; Smith et al., 2007; Zhang et al., 1994; Zhang et al., 1997; Zhong et al., 1998). At a cellular level, the process of Fn1 fibrillogenesis is correlated with the formation of fibrillar adhesions, whereby small mobile adhesions containing Fn1 and integrin *α*5*β*1 somehow elongate while translocating toward the nucleus, first giving rise to focal adhesions (< 1 μm), and then as the translocation continues, to longer filaments, termed fibrillar adhesions (≥1 μm) containing Fn1 and intracellular cytoplasmic effectors linking Fn1 and actin cytoskeleton (Geiger et al., 2001; Geiger and Yamada, 2011; Lu et al., 2020; Pankov et al., 2000; Zamir et al., 1999; Zamir et al., 2000).

It has been thought that Fn1 fibrils resemble ropes, in which extended Fn1 dimers align such that regions containing overlapping N-termini alternate with regions containing C-termini (Chen et al., 1997; Dzamba and Peters, 1991; Fruh et al., 2015), also illustrated in **Sup. Fig. 8A**. To understand how the process of fibrillogenesis occurs in real-time and at a nanoscale level, we adopted a CRISPR/Cas9-mediated mutagenesis approach to generate fluorescently-labeled Fn1, which was subject to the physiological regulation of expression and splicing. This approach has enabled visualization of Fn1 fibrillogenesis over an extended time. Using live imaging and super-resolution microscopy, we uncovered an unexpected mechanism of Fn1 fibrillogenesis.

Our data demonstrate that Fn1 fibrils form as a result of centripetally-moving Fn1 nanodomains originating at the cell periphery. As Fn1 nanodomains move toward the cell center, they assemble into arrays of periodically-spaced nanodomains. The arrays become longer as the movement towards the cell center continues and as more nanodomains are added. We show that these nanodomain arrays are in bona fide fibrils and that each Fn1 nanodomain contains multiple Fn1 dimers. Our live imaging and SMLM reveal a new role by which the N-terminal region of Fn1 protein regulates fibril assembly, and show that interactions mediated by the N-terminal Fn1 assembly region are not required for the formation of Fn1 nanodomains or their centripetal translocation; Instead, the N-terminus of Fn1 regulates the organization of Fn1 nanodomains into nanodomain arrays. Our model integrates the process of fibrillogenesis with the process of adhesion elongation and provides significant new insights into the mechanisms of Fn1 ECM formation, remodeling, and signaling.

## Results

### Beaded structure of Fn1 fibrils revealed by diffraction-limited microscopy

While examining Fn1^+^ ECM in mid-gestation mouse embryos by confocal immunofluorescence microscopy using an Abcam monoclonal anti-Fn1 antibody which binds an epitope within a central region of Fn1 (**Table M2** in Methods), we observed that Fn1 fibrils in the pharyngeal arches and the heart appeared dotted, with regularly-spaced regions of high and low fluorescence intensity (**Fig. 1, Movie 1**). The dotted appearance of embryonic Fn1 suggested that the distribution of Fn1 molecules in Fn1 fibrils is not homogenous. To understand how Fn1 fibrils form, we employed a CRISPR/Cas9 knock-in strategy to modify the endogenous Fn1 locus by replacing the termination codon of Fn1 with a sequence encoding a fluorescent protein. We used this strategy to generate cell lines and Fn1^mEGFP/mEGFP^ homozygous knock-in mice expressing Fn1 fused to monomeric enhanced green fluorescent protein (mEGFP) or other monomeric fluorescent proteins (**Sup. Figs. 1-2**). Homozygous Fn1^mEGFP/mEGFP^ mice were obtained at the correct Mendelian ratio (**Sup. Fig. 1B,** panels 4 and 5), and are viable and fertile, indicating that Fn1-mEGFP supports all functions of Fn1 necessary for embryonic development, fetal viability, and adult homeostasis. Examination of Fn1 expression patterns in knock-in mice showed that Fn1-mEGFP is expressed in the same pattern as the unmodified Fn1 in embryos (Peters and Hynes, 1996), i.e., there were no regions that were GFP+ but Fn1-negative and vice versa in Fn1^mEGFP/+^ embryos expressing one wild-type allele of Fn1 (**Sup. Fig. 2D**). In addition, we used CRISPR/Cas9 mutagenesis to generate five independent lines of mouse embryo fibroblasts (MEFs) expressing Fn1-mEGFP, Fn1-mScarlet-I, Fn1-Neon Green, or Fn1-tdTomato fluorescent proteins (FP). Western blots showed that FP fusions to Fn1 were specific: FPs were only fused to Fn1 as no other FP fusions were detected either by western blotting or immunofluorescence (IF) (**Sup. Fig. 2A-B**).

**Figure 1.**
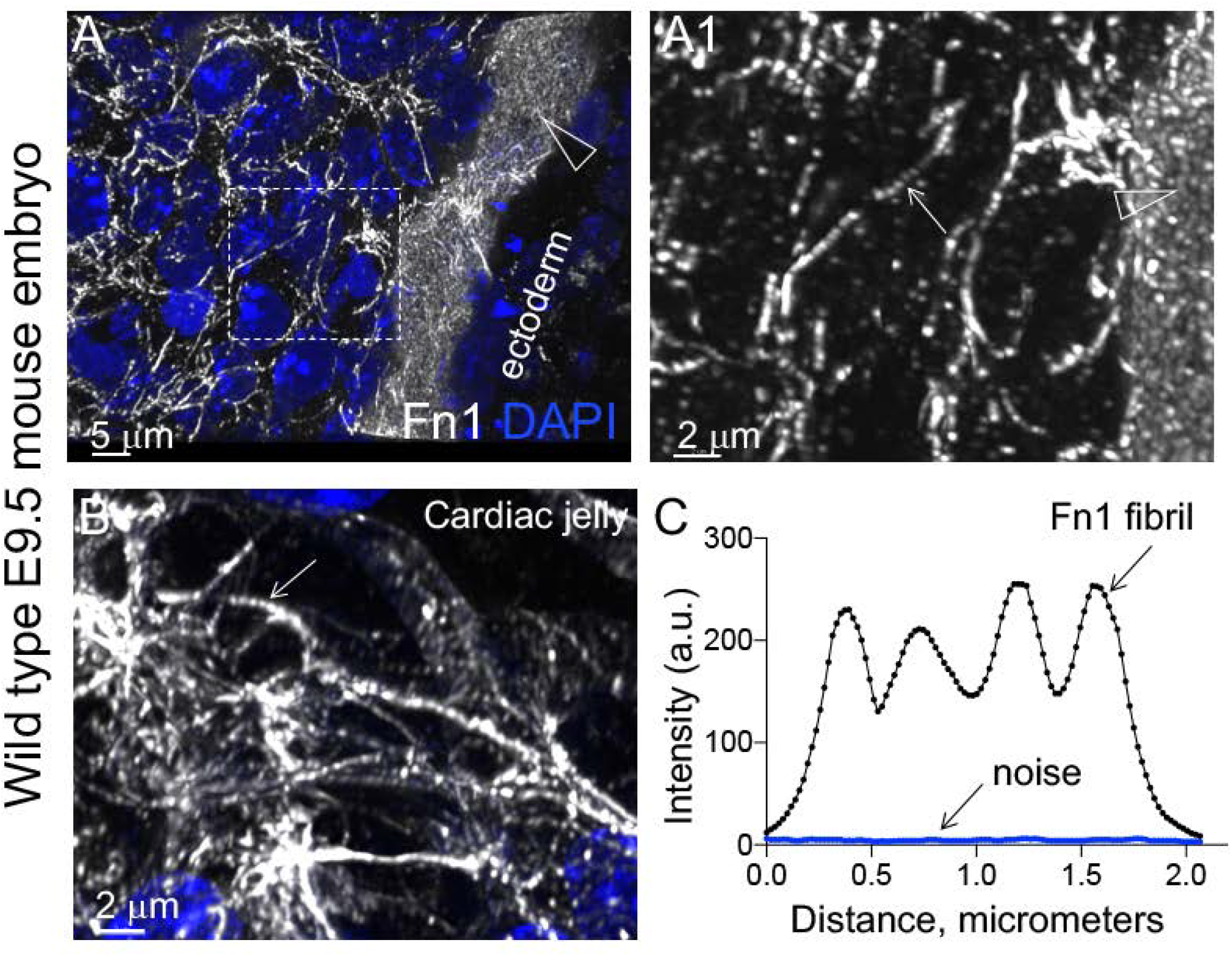
Beaded architecture of Fn1 fibrils in embryonic ECM. Wild-type E9.5 mouse embryos were fixed and stained with the Abeam monoclonal antibody to Fn1 (white) and DAPI (blue), and imaged using 100x oil objective, N.A. 1.49, pinhole 0.8, and sampling rate of 40 nm/pixel. **A–A1.** Sagittal optical section through the first pharyngeal arch and B. the cardiac jelly, an ECM-rich region between the myocardial and endocardial layers of the cardiac outflow tract. Large arrowheads in **A–A1** point to the ECM at the ectoderm-mesenchyme boundary of the 1^st^ pharyngeal arch. The box in **(A)** is expanded in **A1** to show Fn1 microarchitecture. Arrow in **A1** points to a beaded Fn1 fibril within the 1^st^ arch mesenchyme; **B.** Beaded architecture of Fn1 fibrils in cardiac jelly, e.g., arrow. **C.** Intensity profile plot of a Fn1 fibril shows a regularly-undulating profile, with troughs well above the background; a.u. arbitrary units.

Measuring deoxycholate (DOC) insolubility of Fn1 ECM is a classical biochemical assay to assay stable incorporation Fn1 proteins into the ECM (Choi and Hynes, 1979; McKeown-Longo and Mosher, 1983; Schwarzbauer, 1991; Singh et al., 2010; Wierzbicka-Patynowski et al., 2004). For these assays, we carefully controlled the number of cells plated, as cell density affects the extent of Fn1 fibrillogenesis (Hynes, 1990). We performed DOC insolubility assays using 5 independently-generated cells lines expressing Fn1-FP proteins. These experiments demonstrated that the incorporation of Fn1-FPs into the ECM was indistinguishable from wild-type, untagged Fn1 (**Sup. Fig. 2C**). In addition, DOC insolubility assays showed that Fn1-mEGFP proteins isolated from mouse embryos are incorporated into a deoxycholate-insoluble matrix similar to unmodified Fn1 (**Sup. Fig. 2D-E**). Together with the viability of Fn1^mEGFP/mEGFP^ homozygous knock-in mice, these data demonstrate that Fn1-FP fusion proteins are suitable reagents for investigating mechanisms of Fn1 fibrillogenesis.

To visualize the process of fibrillogenesis in real-time, we plated Fn1^mEGFP/+^ MEFs on the gelatin-coated cover glass and imaged cells 16 hours after plating using total internal reflection (TIRF) microscopy at the critical angle of incidence. These experiments showed that Fn1 fibrillogenesis initiated at cell periphery as distinct, brightly-fluorescent Fn1 densities that moved centripetally in parallel with F-actin and became aligned into linear arrays of “beads” (arrows in **Movie 2**). TIRF imaging also showed that domains of higher fluorescence intensity of Fn1 co-localized with integrin *α*5*β*1 both in non-fibrillar (arrows in **Fig. 2A-A2**) and in fibrillar adhesions (arrows in **Fig. 2B-B2**), and that both Fn1 and *α*5*β*1 fibrillar adhesions appeared beaded (**Fig. 2B-B2**, arrows). Beaded architecture of Fn1 fibrils was also observed by an independent imaging method using Zeiss Airyscan technology (**Sup. Fig. 3**). To test whether the beaded appearance of Fn1 fibrils depended on antibody epitope availability, we stained wild-type MEFs or mouse endothelial cells using three different antibodies 1) 297.1 polyclonal antibody raised against the rat plasma Fn1 protein and recognizing multiple Fn1 epitopes (**Sup. Fig. 4**); 2) 3E2 monoclonal antibody recognizing the alternatively spliced EIIIA exon of Fn1, and 3) an Abcam monoclonal antibody recognizing an epitope within the central region of Fn1 (see **Table M2**, in Methods). The beaded appearance of Fn1 fibrils did not depend on the type of antibody used (**Sup. Fig. 5**). Moreover, staining with a polyclonal antibody to Fn1, 297.1, which recognizes multiple epitopes along the Fn1 molecule resulted in the beaded appearance of Fn1 fibrils (**Sup. Fig. 5A-A1**). Fn1 fibrils formed by cells plated on uncoated glass, as well as on glass coated with gelatin, laminin 111, or vitronectin, also appeared beaded **Sup. Figs. 5-6**). Similarly, Fn1 fibrils in cell-free areas (**Sup. Fig. 5A, A1**) and between cells were beaded (**Sup Fig. 5B, B1, C, C1**). Fn1 fibrils produced by cells plated on soft substrata such as hydrogels of variable stiffness also appeared beaded (**Sup. Fig. 6C-D, 6C1-D1**). In this latter experiment, Fn1 was detected by imaging the fluorescence of Fn1-mEGFP protein, indicating that the beaded appearance of Fn1 fibrils is independent of antibody staining. To determine whether Fn1 fibrils in cell-free, fibrillar ECM were beaded, cultures were treated with 2% DOC (Lu et al., 2020). 2% DOC treatment dissolves cell membranes and cytoplasmic components, leaving behind ECM devoid of cell contact (**Movie 3** for time-lapse of dissolution of cellular components, F-actin and DNA). This experiment showed that Fn1 fibrils retained their beaded architecture in the absence of cell contact (**Sup. Fig. 5D**). Taken together, these studies indicated that the beaded appearance of Fn1 fibrils is a general feature of Fn1 ECM that can be seen in multiple contexts: in DOC-resistant cell-free fibrils, in different embryonic tissues *in vivo*, and when cells are cultured on various substrata *in vitro*. Moreover, the beaded appearance of Fn1 fibrils is independent of the type of antibody and a method of detection: It is seen by indirect detection methods such as immunofluorescence and by direct detection of fluorescent Fn1-mEGFP protein.

**Figure 2.**
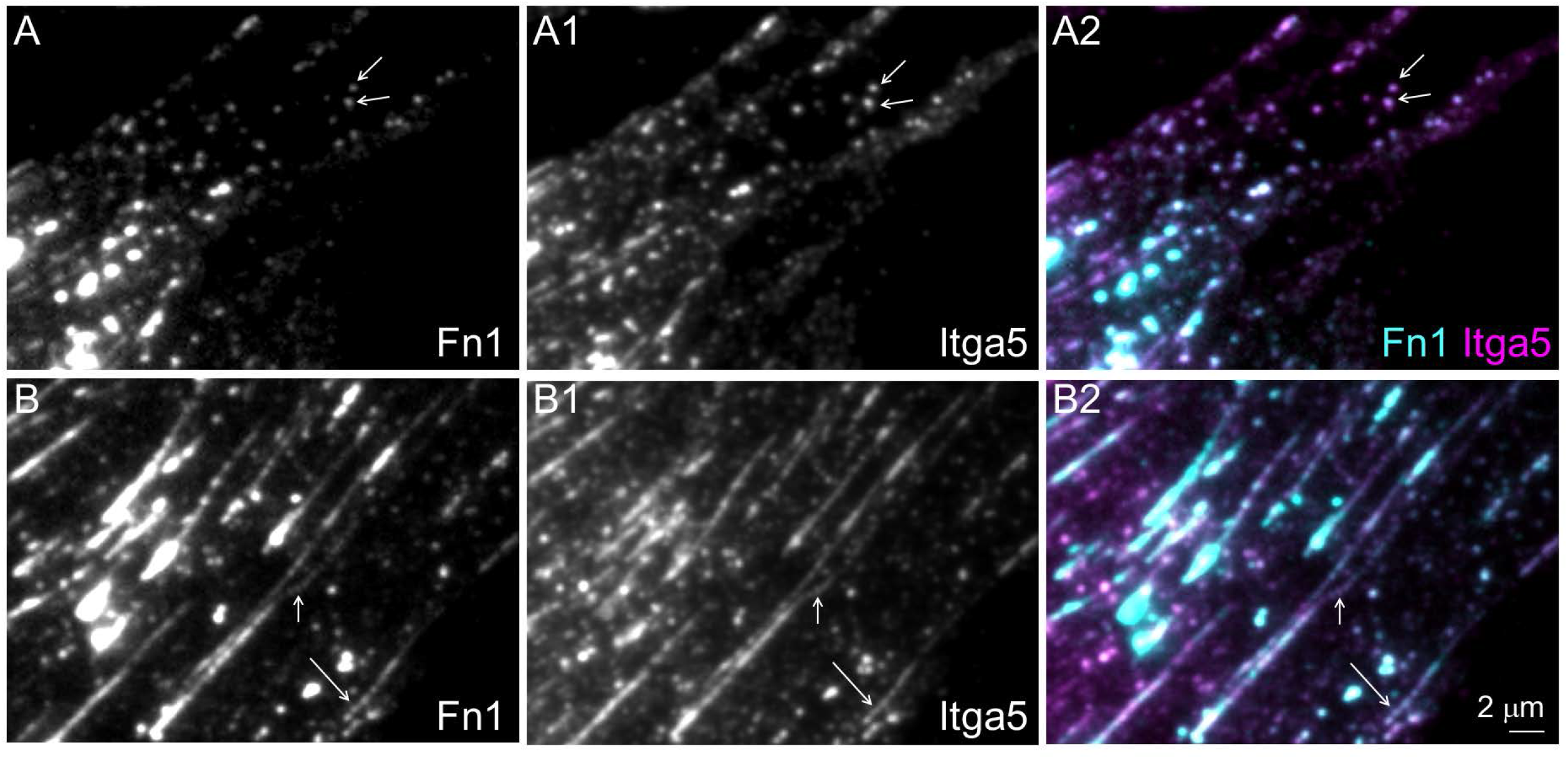
Integrin α5 and Fn1 co-localize in beaded adhesions. Wild-type MEFs were plated for 16 hours on glass coverslips without coating, then fixed, stained with Abeam monoclonal antibodies to Fn1 and anti-integrin α5 (Itga5) antibodies. Cells were imaged at the critical angle of incidence using 100x oil objective, NA 1.49. **A–A2** cell periphery. Arrows in **A–A2** point to examples of non-fibrillar Fn1 adhesions. **B–B2** medial portion of a cell containing beaded fibrillar adhesions (arrows). Magnifications in all panels are the same.

### Nanoarchitecture of Fn1 fibrils revealed by standardized single-molecule localization microscopy

To examine the structure of Fn1 fibrils at a nanoscale level, we adopted single-molecule localization microscopy (SMLM) using the direct stochastic optical reconstruction microscopy (dSTORM) method (Heilemann et al., 2008; Rust et al., 2006). To optimize SMLM imaging conditions and determine the effective labeling efficiency of our reagents, we used the methodology and the reference cell line NUP96-mEGFP, as gold-standard tools to optimize image quality, measure effective labeling efficiency, and to quantify the number of GFP molecules in our structure(s) of interest (Lelek et al., 2021; Thevathasan et al., 2019). In this reference cell line, both copies of the *NUP96* gene are modified by CRISPR/Cas9 mutagenesis to generate NUP96-mEGFP protein, a component of the nuclear pore complex, NPC (Thevathasan et al., 2019). Thirty-two copies of NUP96 protein are evenly distributed among the eight corners of the NPC at known distances (see schematic in **Sup. Fig. 7**) (Bui et al., 2013; Thevathasan et al., 2019; von Appen et al., 2015). To optimize our SMLM imaging, we were guided by the current best SMLM practices (Lelek et al., 2021; Mund and Ries, 2020), the methodology, and the SMAP software developed by (Diekmann et al., 2020; Ries, 2020; Thevathasan et al., 2019). Measurements using the Fourier ring correlation method (Nieuwenhuizen et al., 2013) implemented in the SMAP software indicated that the resolution of our images ranged between 14 and 28 nm. Localization precision was 9.1±1.9 nm, on average. Using antibodies to detect NUP96-mEGFP and the SMAP software, we analyzed images from 3 independent experiments, 8 nuclei and 4571 NPCs, and determined the NPC radius to be 63.6±0.86 nm (**Fig. 3A-C** and **Table 1**). This radius is consistent with the reported NPC radius of 64.3+/-2.6 nm, measured using this cell line and a combination of commercial anti-GFP primary and Alexa-647-conjugated secondary antibodies (Thevathasan et al., 2019). We then used NUP96-mEGFP cells to optimize the effective labeling efficiency (ELE) of our reagents. ELE summarily measures how well one’s reagents and methods, including the SMLM imaging protocol, generate images that reflect the actual structure under the study (Thevathasan et al., 2019). SMLM imaging of NUP96 in the x-y plane should produce 8-fold symmetrical ring-like structures wherein 4 copies of the NUP96 protein are positioned at each of the NPC’s 8 vertices at known intervals and with known dimensions (Thevathasan et al., 2019). Using NUP96-mEGFP as a reference cell line, anti-GFP primary antibodies, commercially-labeled Alexa-647-conjugated secondary antibodies, and the SMAP software (Thevathasan et al., 2019), we analyzed 4,571 NPCs from 8 cells and 3 independent experiments, and determined the ELE to be 79.6±5.4% (**Table 1**), reflecting that the majority of NPCs have eight NUP96^+^ corners in our images (**Fig. 3A-C**). This ELE and the standard deviation of our measurements (<10%) are comparable with the best ELE measured for this system ∼74% (Thevathasan et al., 2019). Together, these experiments indicate that our reagents and SMLM imaging conditions are within the accepted SMLM standards (Lelek et al., 2021).

**Figure 3.**
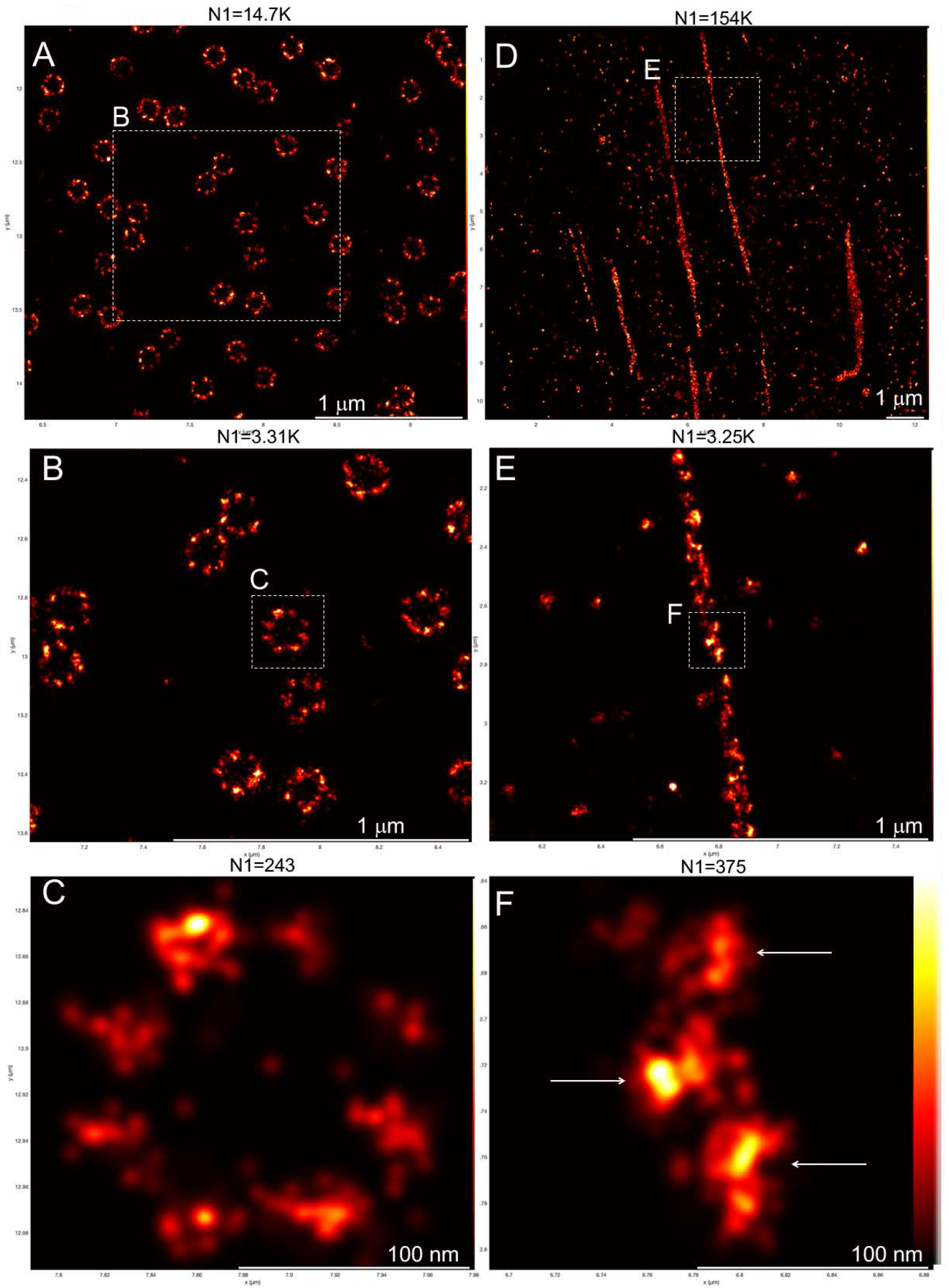
Standardized SMLM imaging results in high resolution and high effective labeling efficiency (ELE). NUP96^mEGFP/mEGFP^ homozygous knock-in cells (**A-C**) and FnimEGFP/mEGFP homozygous knock-in cells (**D-F**) were imaged using the same staining and imaging solutions containing anti-GFP 1° at 1:100 dilution and Alexa-647-conjugated 2° antibodies, the same GLOX/BME buffer, and the same Imaging protocol (*see Methods*). **A-C.** NUP96^mEGFP/mEGFP^ cells. The distribution of Nup96-mEGFP proteins in the majority of nuclear pore complexes (NPCs) exhibits an expected 8-fold symmetry. See *Table 1* for quantification of NPC dimensions and the ELE. The resolution in (**C**) measured by the Fourier ring curve method is 18.3+/-6.7 nm. **D-F.** Fn1^mEGFP/mEGFP^ cells. Nanodomain architecture of Fn1-mEGFP fibrils is clearly visible in subsequent magnifications of the fibril boxed in (**D**). The boxed region in **D** is shown in **E**. Boxed region in **E** is shown in **F**. Arrows in **F** point to Fn1 nanodomains. The resolution in (**F**) measured by Fourier ring curve method is 14.5+/-1.1 nm. Note that nanodomain size is similar to the size of an NUP96+ NPC corner, which Is ∼12 nm in actual diameter. N1 is the number of grouped localizations in each panel. See *Table 2* for quantifications of the numbers of mEGFP molecules per nanodomain. Vertical bar in **F** shows color-coding according to localization density for all panels in this figure.

**Table 1.** Effective labeling efficiency and standardized SMLM measurements. NUP96^mEGFP/mEGFP^ cells were imaged using *α*GFP 1° antibody and AF647-conjugated 2° antibody detecting NUP96-mEGFP; SMLM imaging protocol I was used to acquire data. Data is from 3 independent experiments, eight cells, and 4571 NPCs

| **Sample** | **Buffer** | **ELE (%)** | **Average # Localizations per NPC** | **# Localizations per protein** | **Radius of NPC, nm** |
| --- | --- | --- | --- | --- | --- |
| U2OS-Nup96 | GLOX/BME | 79.6+/- 5.4 | 280 +/- 49.2 | 8.75 +/- 1.54 | 63.6 +/- 0.86 |

To attain a comparable ELE to that measured with NUP96-mEGFP, we stained homozygous Fn1^mEGFP/mEGFP^ MEFs at the same time as NUP96-mEGFP cells, using aliquots of the same mixtures of reagents. Furthermore, pairs of stained Fn1^mEGFP/mEGFP^ MEFs and NUP96-mEGFP cells were imaged on the same day, using aliquots of the same imaging buffer, and the same off-switching and imaging protocols (see Methods). Similar to (Fruh et al., 2015), we saw that Fn1 fibrils appeared as arrays of nanodomains situated along Fn1 fibrils with a regular periodicity (**Figs. 3D-F, Fig. 5D-D2, Fig. 6D-D”**). As in (Fruh et al., 2015), nanodomain periodicity was ascertained using autocorrelation, wherein the position of the first autocorrelated peak marks the nanodomain periodicity (see discussion and simulations in (Fruh et al., 2015)). Autocorrelation analysis of 14 fibrils from 7 cells and 3 independent experiments indicated that the nanodomain periodicity of the GFP epitopes in Fn1-mEGFP fibrils was 102±29 nm (**Figs. 6D-D”, 6F, Table 2**). This periodicity is comparable to the nanodomain periodicity measured by (Fruh et al., 2015), which was 99±17 nm, and the periodicity determined by immuno-electron microscopy using anti-EIIIA antibody, which was on average 84 nm and ranged between 40 and 280 nm (Dzamba and Peters, 1991).

**Figure 4.**
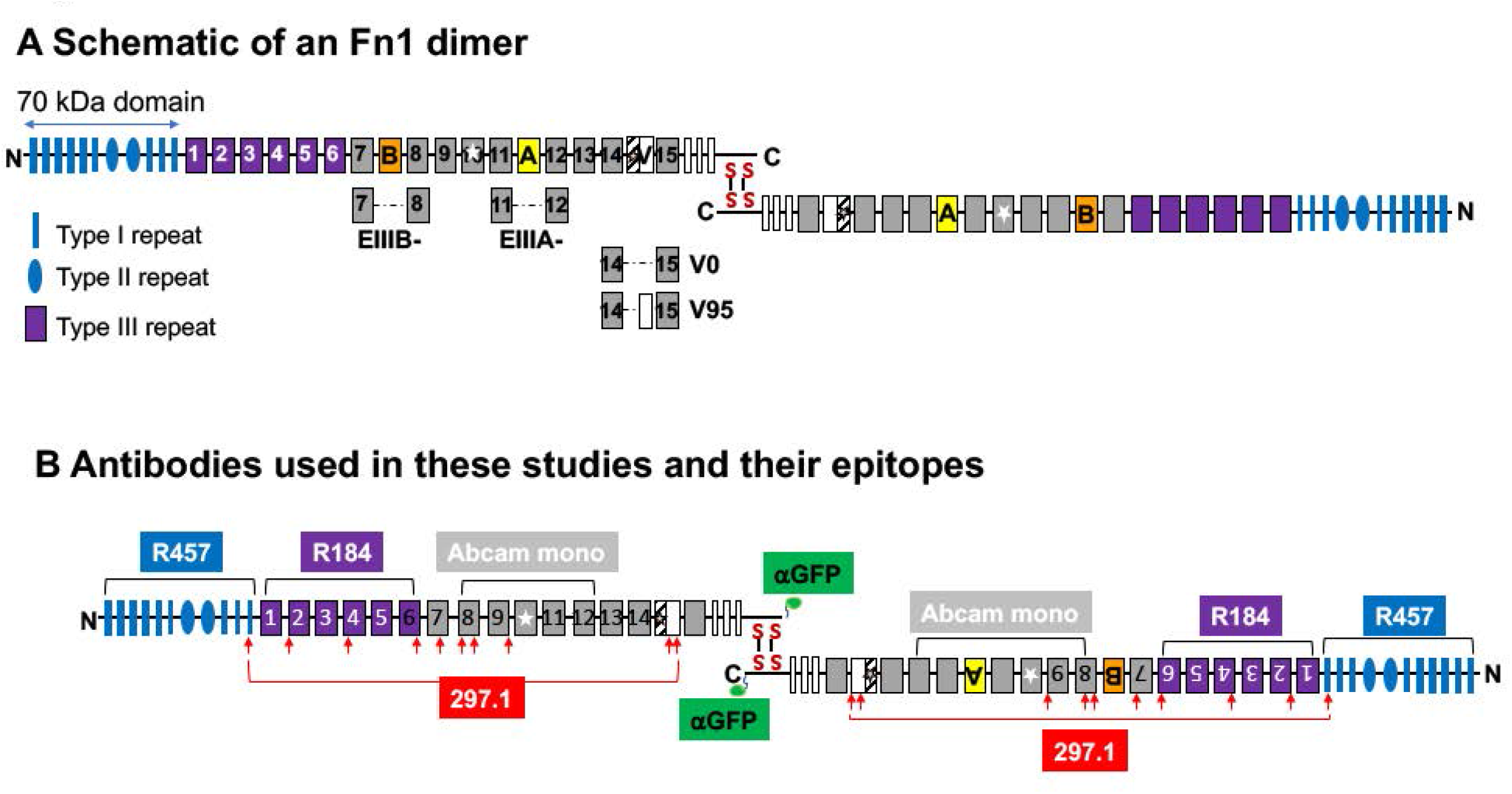
Schematic representations of Fn1 dimers and antibody binding sites. **A.** Fn1 is secreted as a dimer. Two disulfide bonds at the C-terminus hold Fn1 molecules in an anti-parallel orientation. 70 kDA Fn1 assembly domain is marked in blue and by a double-headed arrow. Fn1 type III repeats are numbered. Splice variants characterized in mice are shown. **B.** Schematic of binding sites for the antibodies used in this manuscript. All antibodies except one are polyclonal, see *Table M2* in the Methods section. Arrows mark mapped epitopes. Brackets mark regions of Fn1 recognized by the following antibodies: R457 is a rabbit polyclonal antibody raised to the 70 kDA domain of Fn1; R184 is a rabbit polyclonal raised against the first six type III repeats of Fn1 (purple); 297.1 is a rabbit polyclonal antibody raised to the full length rat plasma Fn1; The binding sites of 297.1 (red arrows) were mapped in this study (see Sup. Fig. 4); Abeam mono is a monoclonal antibody recognizing an epitope within the marked region. αGFP is a polyclonal antibody raised to GFP

**Figure 5.**
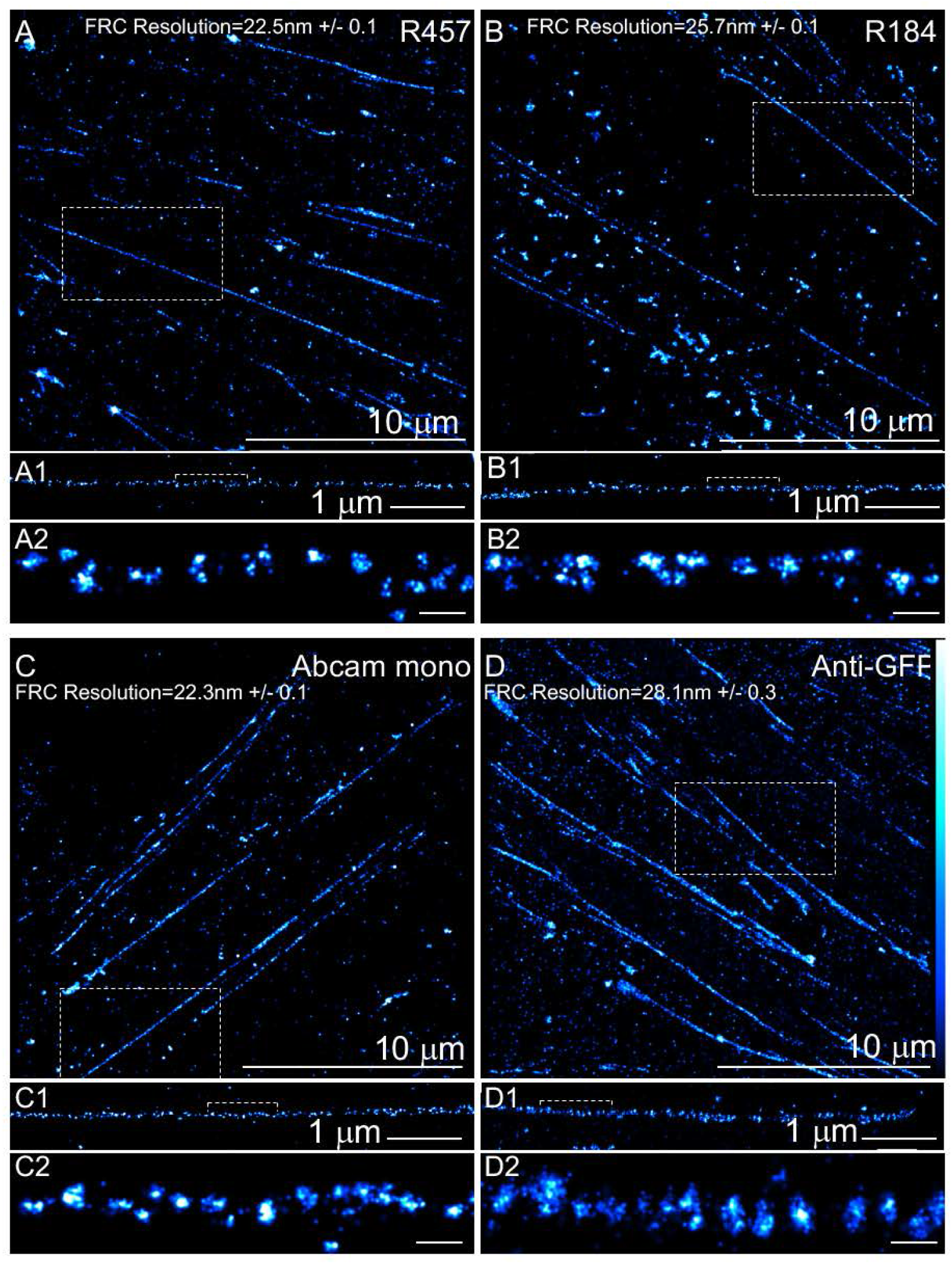
Nanodomain architecture of Fn1 fibrils detected with four different antibodies. Fn1^mEGFP/mEGFP^ MEFs were plated on glass cover slips without coating, cultured over night, and stained with the following antibodies: **A.** R457 (1:200 dilution). **B.** R184 (1:100 dilution). **C.** Abeam monoclonal antibody (1:200 dilution). **D.** anti-GFP antibody (1:100 dilution). Boxes in **A, B, C,** and **D** are expanded to show fibrils in **A1, B1, C1,** and **D1,** respectively. Brackets in **A1, B1, C1,** and **D1** are expanded in **A2, B2, C2,** and **D2,** respectively. Scale bars in **A2, B2, C2,** and **D2** are 100 nm. Fourier ring correlation (FRC) was used to determine image resolution. Vertical bar in **D** shows color-coding according to localization density for all panels in this figure.

**Figure 6.**
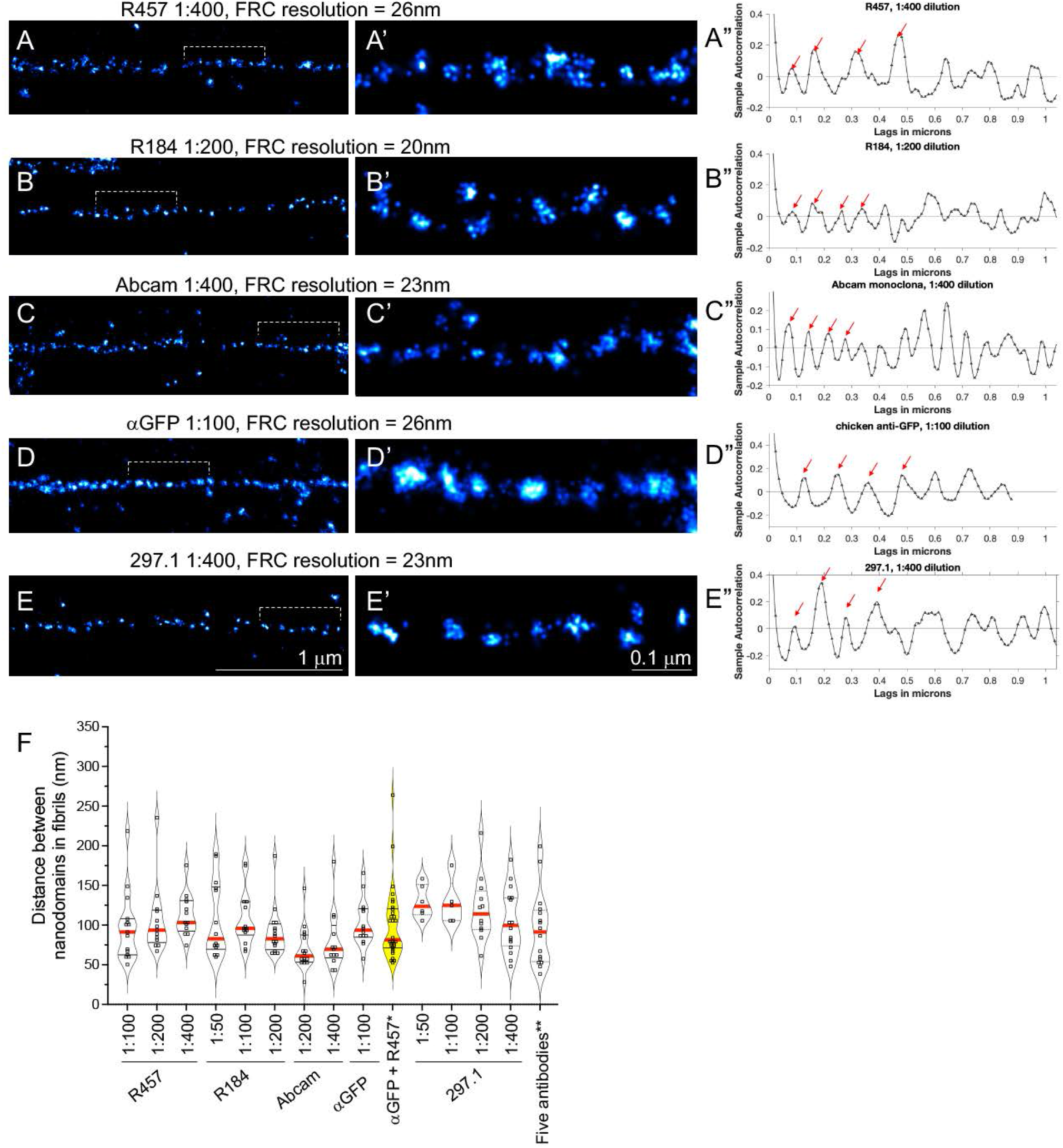
Periodical labeling of Fn1 fibrils by multiple antibodies, their dilutions and combinations. Fn1^mEGFP/mEGFP^ MEFs were plated on glass cover slips without coating, cultured over night, and stained with antibodies Indicated. **A-E.** SMLM images resulting from the highest dilutions of antibodies tested. Brackets in **A–E** are expanded in **A’-E’. A”-E”** Autocorrelation traces for fibrils shown in **A-E.** Ljung-Box Q-test for autocorrelation at the peak positions marked by the red arrows resulted in *h*=1, p=0 indicating strong evidence of autocorrelation at least up to the 4^th^ peak; *h* (the null hypothesis for this test) is that the first 4 autocorrelations are jointly zero. *h*=1 rejects this hypothesis. **F.** Summary of autocorrelation analyses of multiple fibrils (each fibrils analyzed is marked by a square) stained with different antibody dilutions, and combinations of antibodies. Experiments for each antibody dilution were repeated three independent times, and each time, at least three cells were imaged. One-way ANOVA test with Tukey’s correction for multiple testing showed no significant differences among the means. FRC-Fourier ring correlation was used to determine image resolution. * αGFP and R457 were both used at 1:100 dilution. ** The 5 antibodies used and their dilutions were the following: R457(1:400), R184(1:200), Abeam (1:400), αGFP (1:100), 297.1(1:400).

**Table 2.** Characterization of Fn1-mEGFP nanodomains in Fn1 fibrils using standardized SMLM protocol. *α*GFP 1° antibody; AF647-conjugated 2° antibody were used to detect Fn1-mEGFP. SMLM imaging protocol I was used to acquire data. Data is from 3 independent experiments, 6 cells, 52 fibrils and 833 nanodomains were used to calculate the number of GFP molecules per nanodomain; To quantify nanodomain dimeter, data from 3 independent experiments, 15 cells, 27 fibrils, and 1292 nanodomains were used.

|  | Buffer | Average PSF (nm) | Average # Localizations per nanodomain | Median Localization Precision, nm | # of mEGFP molecules per nanodomain in Fn1-GFP fibrils | Diameter of Fn1 nanodomain, nm | Nanodomain periodicity, nm |
| --- | --- | --- | --- | --- | --- | --- | --- |
| Fn1<sup>mEGFP/mEGFP</sup> MEFs; αGFP 1<sup>o</sup> antibody; AF647-conjugated 2<sup>o</sup> antibody | GLOX/BME | 143.8 +/- 2.35 | 159 +/- 77.92 | 181 | 16.85 +/- 5.1 | 28.26+/-9.7 | 102±29 |

### Localization of Fn1 epitopes within periodical nanodomain arrays

Fn1 is a large, multi-domain, ∼250 kDa glycoprotein secreted as a dimer, wherein two Fn1 subunits are linked in an anti-parallel orientation by two di-sulfide bonds at their C-termini (**Fig. 4A**), (Schwarzbauer and DeSimone, 2011; Skorstengaard et al., 1986; Wagner and Hynes, 1979). To investigate the relationship between the protein domains of Fn1 and the nanodomain architecture of Fn1 fibrils, we first used antibodies to different regions of Fn1 protein (**Figs. 4B** and **Table M2**). For these experiments, Fn1^mEGFP/mEGFP^ MEFs were plated on uncoated glass coverslips. Cells were then fixed and stained with antibodies recognizing different Fn1 epitopes: 1) R457 rabbit polyclonal antibodies raised to the N-terminal 70 kDa domain of Fn1 (Aguirre et al., 1994; Sechler et al., 2001), 2) R184 rabbit polyclonal antibodies raised to recognize the domain immediately following the 70 kDa N-terminal domain and containing the first six type III repeats of Fn1 (Fn1 III_1-6_) (Raitman et al., 2018), 3) a rabbit monoclonal antibody from Abcam recognizing an epitope within the central region of Fn1 (**Table M2** for further description of this antibody), and 4) an anti-GFP antibody recognizing the C-terminus of Fn1-mEGFP protein. In our analyses, we focused on long fibrillar adhesions that were over 1 μm in length, characteristic of mature assembled Fn1 fibrils (Lu et al., 2020).

Independent of the antibodies used, Fn1 fibrils appeared as arrays of periodically-spaced nanodomains (**Figs. 5-6**). To ensure that the beaded appearance of fibrils was not due to under-sampling, we tested 2 – 4 dilutions of each antibody, followed by SMLM imaging and autocorrelation analysis, as in (Fruh et al., 2015). Our analyses showed that nanodomain periodicity remained constant at all antibody dilutions tested (**Fig. 5, 6A”-E’, summarized in 6F**). The periodicity of nanodomains detected with either R457, R184, or Abcam monoclonal antibodies was similar to that measured with the anti-GFP antibody, suggesting that each of these antibodies was used at the ELE comparable with that of GFP-labeling reagents. The finding that nanodomain periodicity was independent of antibody specificity supports the conclusion that the beaded appearance of Fn1 fibrils is not a result of a particular Fn1 protein conformation, since staining using antibodies to three different regions of Fn1 in addition to antibodies to GFP resulted in the same pattern. In all cases, nanodomain periodicity was independent of the antibody concentrations, arguing against under-sampling (Fruh et al., 2015).

Analysis using DBSCAN (Caetano et al., 2015) discovered clusters corresponding with nanodomains, when the radius of the neighborhood (*ε*) was set to 14 nm, the average apparent radius of an Fn1 nanodomain (**Table 2**, compare (**Figs. 7A and B**). Unbiased DBSCAN clustering using the neighborhood radius that was automatically estimated by the SMAP software discovered larger clusters of Fn1 nanodomains (**Fig. 7C**). This suggests that Fn1 nanodomains are organized in a higher-order structure within long fibrils, consistent with the observation of “beads” in lower-resolution diffraction-limited images of Fn1 (**Figs. 1, 2** and **Sup. Figs. 3, 5, 6**).

**Figure 7.**
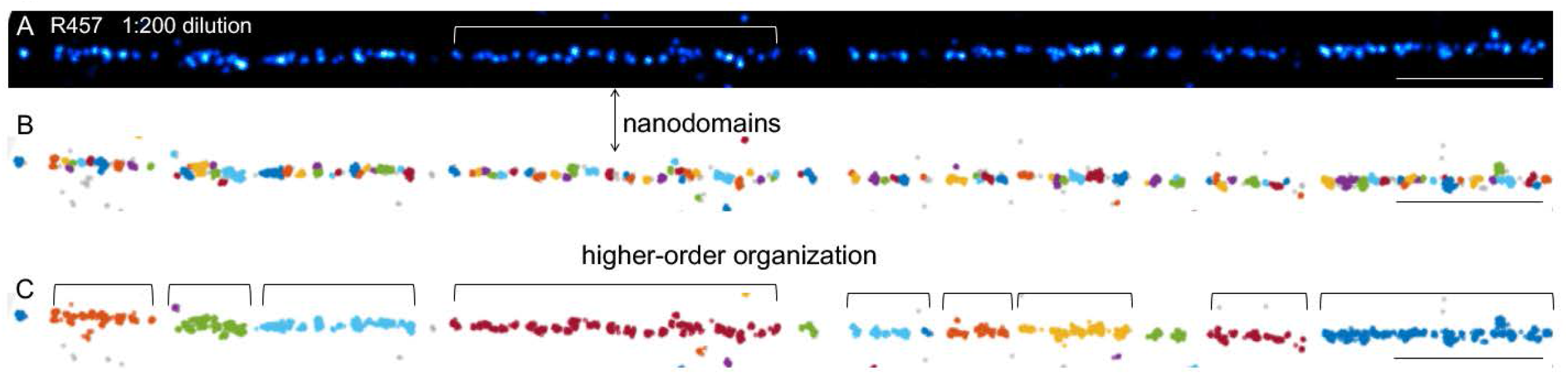
Clustering using DBSCAN suggests a higher-order organization of Fn1 nanodomains in fibrils. Fn1^mEGFP/mEGFP^ MEFs were plated on glass cover slips without coating, cultured over night, and stained with the R457 antibody. The fibril in **A** was analyzed by DBSCAN clustering in **B** and **C.** The minimum number of points per neighborhood (k) was set to 4, as recommended for all 2-dimentional datasets. **B.** The size of the neighborhood, ε, was set to 14 nm, which is an apparent average radius of Fn1 nanodomain (*Table 2*). Adjacent clusters are labeled with different colors. **C.** ε was estimated automatically by the DBSCAN algorithm. Localizations in the same cluster are given the same color; Adjacent clusters are labeled with different colors. DBSCAN clustering in **B** using ε=14 nm discovers Fn1 nanodomains (compare **B** with SMLM image in **A).** Clustering in **C** using an unbiased estimate of ε suggests a higher-order organization of Fn1 nanodomains. Scale bar is 1 micron.

### Fn1 nanodomains contain multiple full-length Fn1 molecules

It was previously-thought that Fn1 nanodomains detected by immunoelectron microscopy or by immunofluorescence SMLM corresponded with particular regions in Fn1 protein sequence (Dzamba and Peters, 1991; Fruh et al., 2015), illustrated in **Sup. Fig. 8A-B**. Surprisingly, staining using 297.1 polyclonal anti-Fn1 antibody raised against full-length plasma Fn1 and recognizing multiple epitopes along Fn1 protein resulted in the same periodicity of nanodomains as staining with antibodies to distinct regions of Fn1 protein (**Figs. 6E, 6F, Figs. 8A, 8D**). Since 297.1 polyclonal antibody recognizes multiple epitopes along Fn1, including an epitope in the 70 kDA N-terminal assembly region of Fn1 (**Sup. Fig.4**), these experiments suggested that each Fn1 nanodomains contained at least one entire Fn1 dimer. To determine the number of Fn1 molecules per nanodomain, we used the methodology developed by (Thevathasan et al., 2019), and identical labeling reagents and imaging conditions for the detection of mEGFP in NUP96-mEGFP and Fn1-mEGFP cells (Thevathasan et al., 2019). This analysis indicates that each Fn1 nanodomain contains 16.85 +/- 5.1 mEGFP proteins on average (**Table 2).** Since Fn1 assembled in ECM fibrils is an obligate dimer of two disulfide-bonded Fn1 molecules (Schwarzbauer, 1991), these findings show that Fn1 nanodomains contain 6 – 11 Fn1-mEGFP dimers, on average.

**Figure 8.**
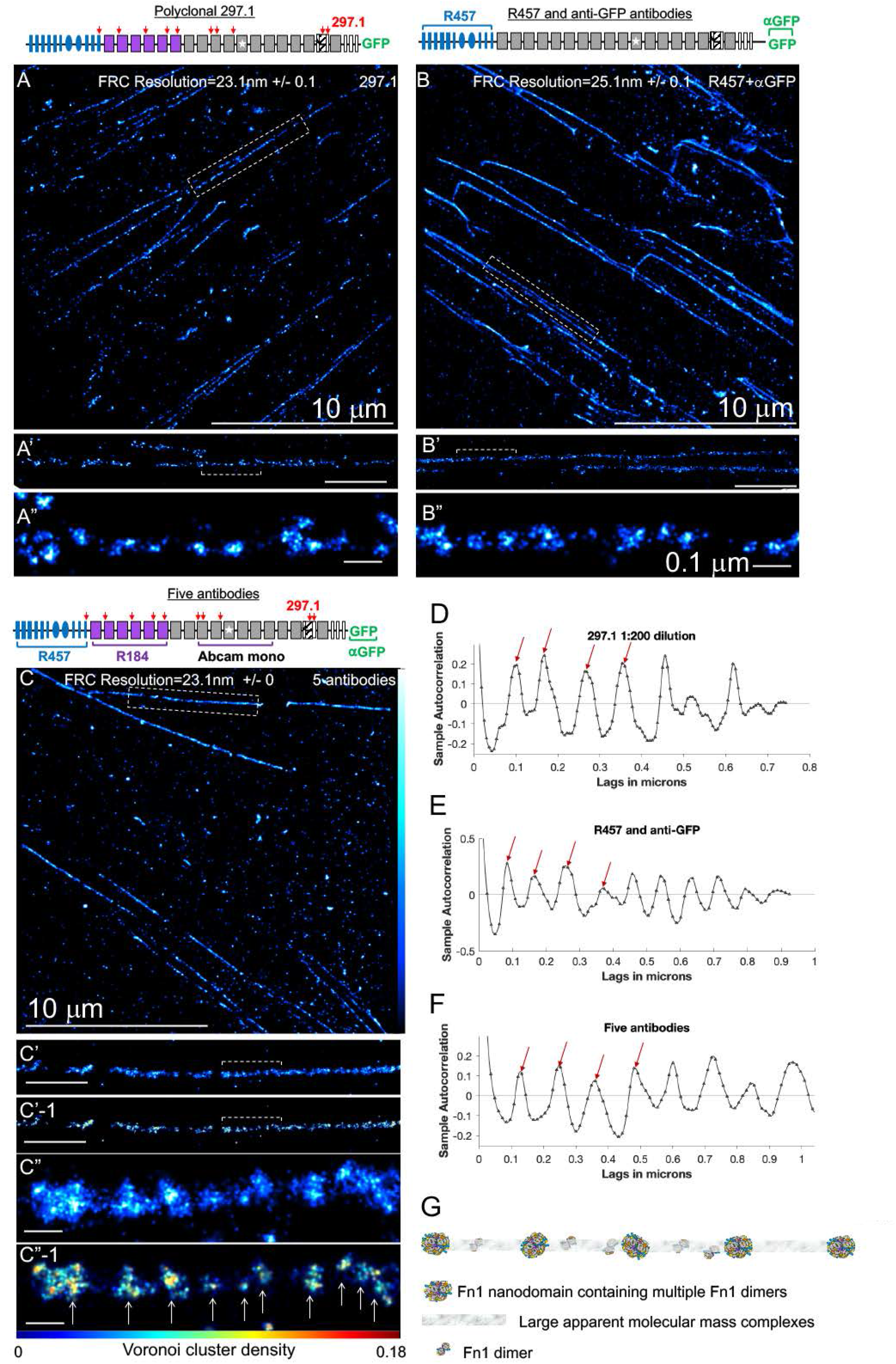
Polyclonal antibody to full-length Fn1 and combinations of antibodies recognizing epitopes along the length of Fn1 reveal periodical nanodomain architecture of Fn1 fibrils. Fn1^mEGFP/mEGFP^ MEFs were plated on glass cover slips without coating, cultured over night, and stained with the indicated antibodies. **A.** 297.1 rabbit polyclonal antibody raised to full-length, rat plasma Fn1 recognizing multiple epitopes along Fn1 molecule (red arrows), 1:200 dilution. **B.** A combination of anti-GFP antibody (1:100) and R457 antibody (1:100). **C.** Staining using a combination of 5 antibodies depicted in a schematic above the panel. For dilutions of 5 antibodies see legend to Fig. 6F. Boxed fibrils in **A, B, C** are expanded in **A’, B’, C’** and **C’-1** (scale bars in **A’,B’,C’,** and **C’-1** are 1 μm). Bracketed regions in **A’, B’, C’** are expanded in **A”, B”, C”** and **C’-1** (scale bars in **A”, B”, C”** and **C”-1** are 0.1 μm). **C’-1** and **C”-1.** Vertical bar in **C** shows color-coding according to localization density for all panels in this figure, except **C’-1** and **C”-1. C’-1** and **C”-1** are color-coded according to Voronoi cluster density (horizontal bar below **C”-1**); Clustering analysis based on Voronoi tessellation segments clusters corresponding with nanodomains (arrows in **C”-1**). **D–F.** Autocorrelation analysis and Ljung-Box Q-test for autocorrelation at the peak positions marked by the red arrows resulted in *h*=1, p=0 indicating strong evidence for autocorrelation at least for the 4 peaks; *h* (the null hypothesis for this test) is that the first 4 autocorrelations are jointly zero. *h*=7 rejects this hypothesis. **G.** A model of Fn1 fibril. In this model Fn1 fibrils are composed of periodically-spaced nanodomains containing multiple Fn1 dimers. Nanodomains may be linked via large molecular mass complexes (LAMMs, Zhang and Mosher, 1996) with a few Fn1 dimers between the nanodomains. For additional/alternative models see *Discussion* and *Sup.* Fig. 8*F1-F2*.

These results are not consistent with the model that thinnest Fn1 fibrils are made of extended single Fn1 molecules that are periodically aligned in a regular, end-to-end fashion with regions containing N-termini alternating with regions containing C-termini (Chen et al., 1997; Dzamba and Peters, 1991; Fruh et al., 2015) and illustrated in **Sup. Fig. 8A-B**. Such periodic alignment of Fn1 dimers necessitates that staining using antibodies recognizing multiple epitopes along the length of Fn1 molecule would result in a uniform labeling of Fn1 fibrils, (**Sup. Fig. 8C**). However, this is not what we observed: staining with a polyclonal 297.1 antibody showed that Fn1 fibrils were composed of periodically-spaced nanodomains (**Fig. 8A-A”, 8D**). Autocorrelation analysis showed that the spacing was periodical at multiple concentrations of 297.1 antibody (**Figs. 6E-E”, 6F,** and **8D**). Importantly, the periodicity of nanodomains detected using 297.1 antibody was statistically indistinguishable from periodicities seen with any regions-specific antibody tested, including that of anti-GFP antibodies (**Fig. 6F**). The latter suggested that 297.1 polylconal antibodies were used at an ELE that was at least as high as that for GFP labeling reagents. Finally, the periodical distribution of 297.1 epitopes (average periodicity 121±11) and the presence of multiple Fn1 dimers per nanodomain, does not fit with the model proposed by (Dzamba and Peters, 1991), illustrated in **Sup. Fig. 8A**, and suggest a different model of Fn1 fibrillogenesis (e.g., **Fig. 8G**).

To further evaluate the hypothesis that Fn1 nanodomains contain full-lengths Fn1 dimers, we used combinations of antibodies and the imaging parameters resulting in the high ELE (**Fig. 6F** and Methods). We first stained Fn1^mEGFP/mEGFP^ MEFs using a combination of R457 and anti-GFP antibodies, detecting the N-and C-termini of Fn1-mEGFP, respectively. Consistent with the staining using 297.1 antibody, the periodicity of nanodomains detected with a mixture of R457 and anti-GFP antibodies was 101±45 nm, which is statistically indistinguishable from the periodicities when either of these antibodies were used alone (**Figs. 6F, 8B-B”, 8E**).

Next, we stained Fn1^mEGFP/mEGFP^ MEFs with a cocktail of five antibodies recognizing epitopes distributed along the length of the Fn1 molecule: 1) the N-terminal 70 kDa assembly domain of Fn1 (R457), 2) the sequence immediately following the 70 kDA domain (R184), 3) an epitope in the middle of Fn1 (Abcam monoclonal), 4) multiple epitopes along the Fn1 protein (297.1), and 5) the anti-GFP antibody marking the C-terminus of Fn1-mEGFP (*α*GFP) **(Fig. 8C-C”, 8F**). Each antibody was used at a dilution resulting in a periodical staining (**Fig. 6F** and legend). The binding of all primary antibodies in the cocktail was detected by a cocktail of Alexa 647-conjugated secondary antibodies. If Fn1 fibrils were indeed composed of continuous, linear arrays of extended and periodically aligned Fn1 molecules, this cocktail of antibodies recognizing epitopes from the N- to the C-termini of Fn1 would uniformly label Fn1 fibrils. Even in the scenario in which R457 epitopes did not extend to the N-terminal-most sequence of Fn1, the staining using this mixture of five antibodies would not be expected to produce periodical staining. However, we observed that the cocktail of 5 antibodies labeled nanodomains with the average periodicity of 94±47 nm at the spatial resolution of 23 nm in thin Fn1 fibrils (**Fig. 8C-8C”, 8F**). This periodicity was statistically indistinguishable from the periodicities of individual antibodies at all the dilutions tested (**Fig. 6F**).

Unbiased clustering using the Voronoi tessellation method (Andronov et al., 2016b) implemented in SMAP showed that Fn1 fibrils stained with the 5-antibody cocktail can be segmented into clusters (**Fig. 8C”, 8C”-1**). This analysis shows that the nanodomain structure of Fn1 fibrils can be discovered using an unbiased computational approach. The size of the nanodomains detected with the 5-antibody cocktail was larger than when nanodomains were detected by the use of any single antibody (e.g., compare **Fig. 8A”** with **Fig. 8C”**). This type of behavior is expected in SMLM when molecules are clustered (Baumgart et al., 2016).

In agreement with the results described above, periodically-spaced nanodomains were detected in Fn1 fibrils using two other cell lines, heterozygous Fn1^mEGFP/+^ MEFs and wild-type MEFs (**Sup. Fig. 9**). The presence of the periodical nanodomains in fibrils produced by wild-type cells indicates that the periodical structure of Fn1 fibrils is not an artifact of GFP labeling. Nanodomain spacing in Fn1 fibrils treated with 2% DOC, which removes cells and cellular components, was similar to that of untreated fibrils (**Sup. Fig. 9**, columns **4** and **8,** and **Sup. Fig. 9F**). Together, these data indicate that the nanodomain architecture of Fn1 fibrils is independent of antibody(ies) used for staining, is seen in cells expressing wild-type Fn1 and in the absence of cell contact, and that periodically-spaced nanodomains within Fn1 fibrils contain multiple Fn1 dimers (e.g., model in **Fig. 8G**).

To further test the hypothesis that Fn1 nanodomains contain full-length Fn1 molecules, we performed double-color dSTORM experiments, wherein the N-terminus of Fn1 was labeled with the R457 primary antibody and anti-rabbit CF680-conjugated secondary immunoglobulins, and the C-terminus of Fn1-mEGFP was labeled with the anti-GFP primary antibody and Alexa-647-conjugated anti-chicken secondary antibodies. As a control, we stained Fn1^mEGFP/mEGFP^ MEFs using two different anti-GFP antibodies, one made in rabbit (detected using CF680-conjugated secondary antibodies) and the other, in chicken (detected with Alexa-647-conjugated anti-chicken secondary antibodies). The majority of nanodomains detected using the two anti-GFP antibodies contained overlapping CF680 and AF647 localizations, suggesting a high ELE for both reagents (**Fig. 9A-C,C’**; Gray lines in **Fig. 9A’, 9B’** mark overlapping staining in nanodomains). Nanodomains in which the two labels did not overlap were marked by purple lines (**Fig. 9A’, 9B’**). To determine the extent of co-localization between the two labels, we performed coordinate-based colocalization (CBC) analysis, a widely-accepted method for detecting co-localization in SMLM data (Malkusch et al., 2012). The CBC coefficient was calculated by counting the number of CF680 localizations within a radius *r* of each of the AF647 localizations and normalizing by the number of localizations within the area *π*R^2^_max_ (Malkusch et al., 2012) (**Fig. 9D,F**). Conversely, we also determined the CBC coefficient by counting the number of AF647 localizations within a radius *r* of each of the CF680 localizations and normalizing by the number of localizations within the area *π*R^2^_max_ (**Fig. 9E,F**). The CBC coefficients were calculated using algorithms implemented either by the Abbelight software or by the publicly-available ThunderSTORM software (Ovesny et al., 2014). The coefficient “+1” indicates a high probability of co-localization, while CBC ≤ 0 indicates low probability of co-localization. CBC analysis using the Abbelight software and color-coding according to the CBC coefficient, showed that the parameters chosen for these analyses, *r*=30nm and R_max_=300nm, reflect the overlap (e.g., yellow color in **Fig. 9D-E** and gray lines in **Fig. 9A’-B’**) and the lack of overlap (purple color and arrows in **Fig. 9D-E** and purple lines and arrows in **Fig. 9A’-B’**) seen in SMLM images. These figures also show that we can detect non-overlapping signals in nanodomains located next to one another (**Fig. 9D-F**). CBC analyses performed on twenty-two regions containing long (≥1μm) Fn1 fibrils from Fn1^mEGFP/mEGFP^ MEFs showed extensive overlap among localizations arising from rabbit and chicken anti-GFP antibodies (**Fig. 9F**).

**Figure 9.**
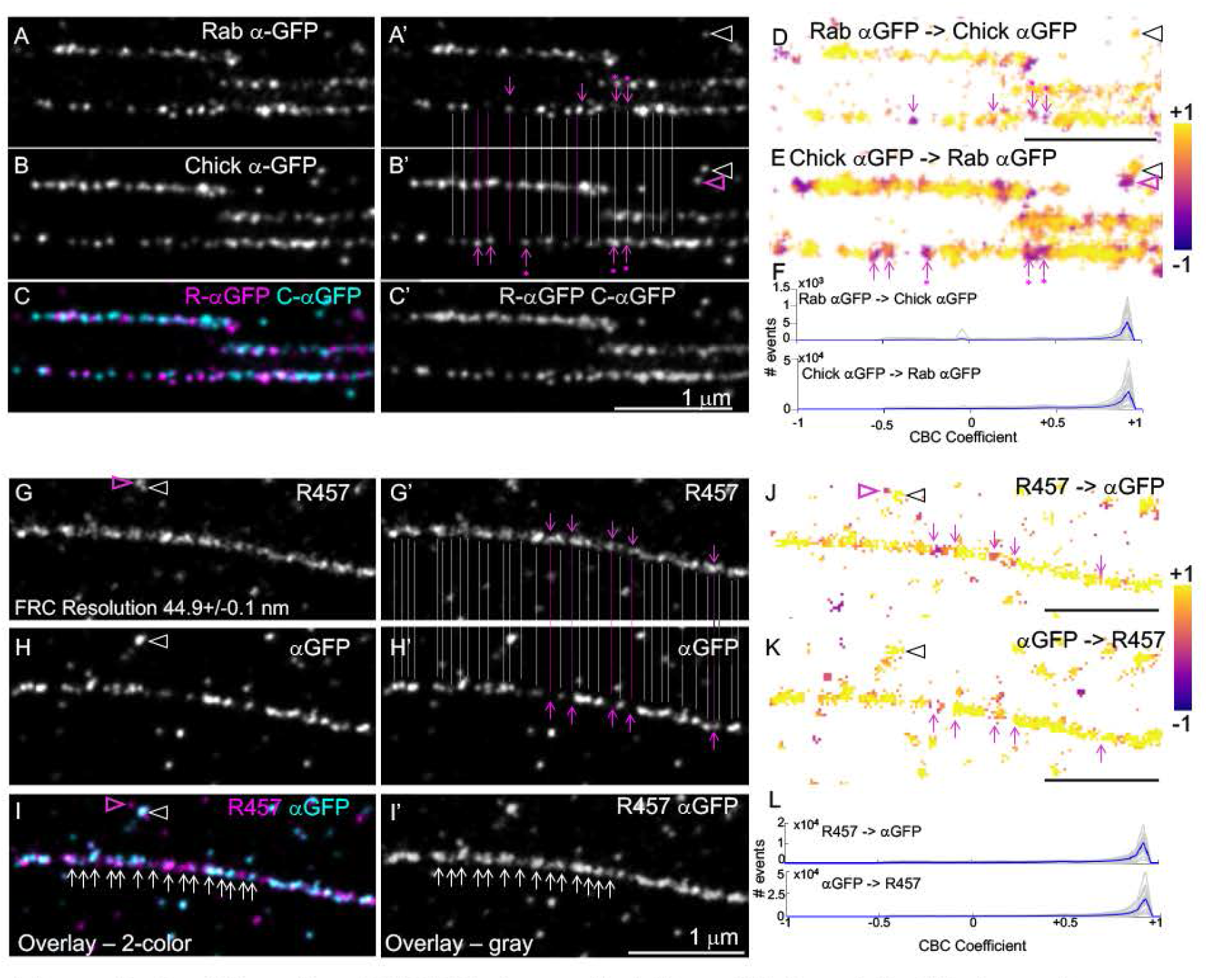
Double-color dSTORM shows that N- and C-termini of Fn1 overlap within Fn1 nanodomains. Fn1^mEGFP/mEGFP^ cells were plated on glass without coating and stained. **A–F.** Control staining. Cells were stained using polyclonal rabbit and chicken anti-GFP antibodies (anti-rabbit 2° antibodies were conjugated to CF680 and anti-chicken 2° antibodies were conjugated to AF640). Panels **A-B** were also displayed as **A’-B’** with vertical lines marking overlapping (gray lines) and non-overlapping (pink lines and arrows) nanodomains; asterisks mark nanodomains in which the signal in one channel is brighter than in the other; **C.** Overlap of two channels in color and **C’** in gray-scale. Less than100% overlap is expected since ELE < 100%. **D-E.** Localizations in **A-B** are color-coded according to the coordinate-based colocalization (CBC) analysis (R_max_=300, r=30). +1: overlap; ≤ 0: no overlap. Clear arrowheads in **A’, B’, D,** and **E** point to an overlapping staining, and pink arrowheads point to an adjacent non-overlapping staining. Pink arrows and asterisks in **A’-B’** are carried over to **D** and **E,** respectively to point non-overlapping staining in fibrils. Asterisks under arrows marks those nanodomains in which the signal in one channel is higher than in the other, but otherwise they co-localize. **F.** CBC coefficients for 22 regions containing long fibrils (gray traces) and their average (blue trace) are plotted. **G-L.** Fn1^mEGFP/mEGFP^ cells were stained using rabbit polyclonal R457 antibody (N-terminus of Fn1) and chicken anti-GFP antibody (C-terminus of Fn1-mEGFP), 2° antibodies were as in **A-F. G-H** were duplicated as **G’-H’** with gray lines indicating co-localizing staining and pink lines pointing to non-colocalizing staining. **I-I’** show merged signals for the two antibodies in color (**I**) and gray scale (**I’**). Arrows in **I-I’** point to nanodomains. White and pink arrowheads in **G-I** also show overlapping and non-overlapping staining, respectively. **J-K.** Images in **G-H** are color-coded according to the (CBC) coefficient, (R_max_=300nm, r=30 nm). Pink arrows in **G’-H’** correspond with pink arrows in **J-K,** respectively. Asterisk under the arrow marks the nanodomains in which the signal in **H’** is higher than in **G’,** but otherwise they co-localize. Note that the CBC analyses distinguish overlapping and non-overlapping localizations in adjacent nanodomains. **L.** CBC coefficients for 20 regions containing long fibrils (gray traces) and their average (blue trace) are plotted, showing a high probability of a near complete overlap between the N- and C- termini of Fn1. Magnifications are the same in all panels. All scale bars are 1 μm. For additional CBC analyses (R_max_=50nm, r=5nm) see Sup. Fig. 10.

CBC analyses of twenty regions containing long fibrils from Fn1^mEGFP/mEGFP^ MEFs stained with R457 and anti-GFP antibodies showed extensive overlap between R457 and anti-GFP localizations, detecting the N- and the C-termini of Fn1-mEGFP, respectively (**Fig. 9G-L**). Using ThunderSTORM software, we varied the parameters of CBC analysis, e.g., setting the radius *r* from 2 or 5 nm and R_max_ from 20 or 50nm, respectively. These analyses had the same outcome as above, i.e., CBC coefficients were close to +1, indicating a high probability of overlap among localizations resulting from R457 and anti-GFP antibody binding (**Sup. Fig. 10**). Together, these studies show that the N- and the C-termini of Fn1 are contained within each Fn1 nanodomain and support the model wherein Fn1 nanodomains in Fn1 fibrils contain multiple full-length Fn1 molecules (**Fig. 8G**).

### Nanoarchitecture of Fn1 fibrils formed by ectopically-added Fn1

It is known that ectopic Fn1 added to cells becomes assembled into fibrils (McKeown-Longo and Mosher, 1983). To determine whether fibrils incorporating ectopically-added Fn1 contain nanodomains, we first used live imaging to film the assembly of ectopic Fn1. In these experiments, MEFs producing Fn1-mEGFP were cultured on the uncoated glass inside an insert while a confluent monolayer of MEFs producing Fn1-tdTomato was grown in the space surrounding the insert. Prior to imaging, the insert was lifted, and Fn1-mEGFP+ MEFs were imaged at ∼17 min intervals for 16 hours. These movies show that Fn1-tdTomato fibrils become clearly visible on the surface of Fn1-mEGFP cells by about 3 hours of co-culture without first accumulating inside cells, suggesting that ectopic Fn1 is assembled into fibrils at the cell surface (**Movie 4**). To determine the nanoarchitecture of Fn1 fibrils formed by ectopically-added Fn1, we cultured Fn1^mEGFP/mEGFP^ MEFs in the presence of 10 μg of Fn1-tdTomato for 24 hours. Cells were fixed and stained without permeabilization using rabbit anti-Cherry primary antibodies and anti-rabbit Alexa647-conjugated secondary antibodies. ECM in regions between cells was imaged using TIRF to detect Fn1-mEGFP and Fn1-tdTomato fluorescence, and dSTORM to detect Fn1-tdTomato due to the Alexa 647 label. Fn1-tdTomato extensively co-assembled with Fn1-mEGFP (**Sup. Fig. 11A-B**), and high-resolution SMLM reconstructions of thin and thick fibrils contained nanodomain arrays composed of Fn1-tdTomato proteins (**Sup. Fig. 11C-E, 11F)**. Together, these experiments demonstrate that 1) Fn1 fibrils deposited into the intercellular ECM space are composed of nanodomain arrays and 2) ectopic Fn1 assembles into fibrils comprised of nanodomain arrays.

### Fibrillogenesis inhibitor FUD disrupts the organization of Fn1 nanodomains into periodical arrays

To understand the relationship between the nanodomain architecture of Fn1 fibrils and the process of fibrillogenesis, we adopted a live imaging approach using Fn1^mEGFP/+^ MEFs and inhibitors of fibrillogenesis. One such inhibitor is a 49-amino acid peptide derived from *Streptococcus pyogenes* adhesin F1, termed the functional upstream domain (FUD), a highly potent inhibitor of Fn1 fibrillogenesis (Tomasini-Johansson et al., 2001). Fn1 fibrillogenesis critically depends on the interactions mediated by the N-terminal 70 kDa assembly domain of Fn1 (**Fig. 4A**), and FUD is one of the inhibitors that interferes with these interactions (Filla et al., 2017; Morla et al., 1994; Schwarzbauer, 1991; Sechler et al., 2001; Sechler et al., 1996; Tomasini-Johansson et al., 2001).

To further investigate the mechanism of Fn1 fibrillogenesis and the role of the N-terminal domain of Fn1 in this process, Fn1^mEGFP/+^ MEFs were plated on uncoated glass for 4 hours, and then imaged for 15 – 18 hours either in the imaging medium alone, or in the medium containing 225 nM FUD. We also imaged cells incubated with 274 nM 11-IIIC peptide, a 68 amino-acid control peptide that does not interfere with Fn1 fibrillogenesis (Morla et al., 1994; Sottile and Chandler, 2005). Untreated cells or cells treated with the control peptide developed and accumulated long Fn1 fibrils (**Movie 5**). In contrast, treatment with FUD led to a) the dismantling of the pre-existing Fn1 fibrils and b) inhibiting the formation of new Fn1 fibrils (**Movie 6)**. Instead of fibrils, cells cultured in the presence of FUD mainly contained centripetally-moving Fn1-mEGFP fluorescent “beads” that only rarely assembled into fibrils (**Movie 6**). These experiments suggested that FUD inhibits fibrillogenesis by interfering with the process by which Fn1 “beads” become arranged or connected into linear arrays. To test this hypothesis, Fn1^mEGFP/+^ MEFs were plated for 16 hours in the continuous presence of either 225 nM FUD or 274 nM III-11C control peptides or were left untreated. Cells were then washed with PBS, fixed, and stained without permeabilization using monoclonal anti-Fn1 antibodies and Alexa Fluor 647-conjugated secondary antibodies, and imaged at the critical angle of incidence by STORM (Jimenez et al., 2020). This approach maximizes the detection of cell-surface Fn1 due to: a) the absence of a detergent during fixation, staining and washing and b) imaging at the critical angle of incidence to detect fluorescence in close proximity to the plasma membrane. These experiments demonstrated that the organization of Fn1 nanodomains into linear arrays was lost upon incubation with FUD (compare **Fig. 10A, B, A1, B1** with **Fig. 10C, C1**). Non-fibrillar (NF) Fn1 nanodomains in cells treated with FUD had a similar number of Fn1 localizations per nanodomain, and were of similar sizes compared with fibrillar or non-fibrillar Fn1 nanodomains in untreated cells, or cells incubated with the control peptide (**Fig. 10A2, 10B2, 10C2**, **10D**). Taken together, these data indicate that FUD does not interfere with the formation of Fn1 nanodomains but inhibits the organization of Fn1 nanodomains into linear arrays. Since Fn1 proteins lacking the N-terminal assembly domain do not form fibrils (Schwarzbauer, 1991), our experiments suggest that interactions mediated by the N-terminal assembly domain of Fn1 with a yet unidentified factor(s), are critical for the organization of Fn1 nanodomains into fibrillar arrays (**Fig. 10E** and *vide infra*).

**Figure 10.**
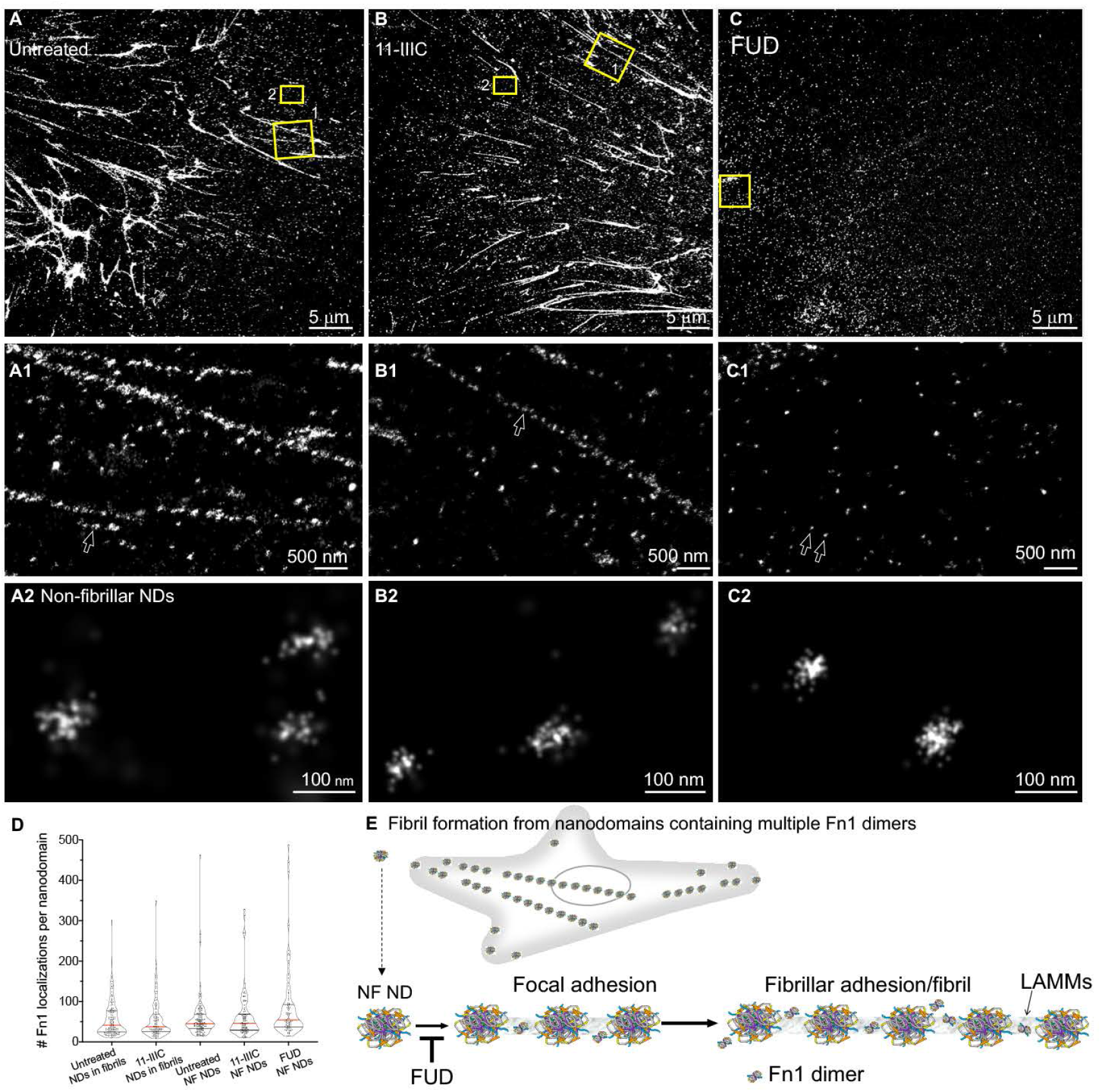
The N-terminal Fn1 assembly domain regulates the organization of Fn1 nanodomains into linear fibrillar arrays. Fn1^mEGFP/+^ MEFs were plated on glass and were either left untreated (**A-A2**), Incubated with the control 11-IIIC peptide (**B-B2**), or the FUD peptide (**C-C2**) for 16 hrs. Cells were then washed with PBS, fixed and stained with the Abeam monoclonal antibody to Fn1 followed by Alexa 647-conjugated secondary antibodies. Cells were Imaged at the critical angle of Incidence by dSTORM. **A-A2.** Untreated, non-permeabilized cells. B - B2. Cells Incubated with control 11-IIIC peptide, non-permeabilized. **C–C2.** FUD-treated, non-permeabilized cells. Boxes marked 1 in **A-B** are expanded in A1-B1. Boxes marked 2 in A-B, are expanded in A2 - B2. The box in C Is expanded in C1 and C2. Arrows in A1-B1 point to Fn1 nanodomains (NDs) in fibrils. Arrows in C1 point to non-fibrillar (NF) nanodomains expanded in **C2**; **D.** Quantification of the number of grouped Fn1 localizations in nanodomains in fibrils and in non-fibrillar nanodomains after various conditions. Red lines mark medians. Differences are not statistically significant, Kruskal-Wallis test with Dunn’s correction for multiple testing. Cells from all experimental conditions were Imaged using Identical conditions and high laser power (see SMLM Imaging protocol II in Methods) explaining lower average number of grouped localizations per nanodomain. **E.** Model of fibril formation in a cell: 6-11 Fn1 dimers assemble Into nanodomains containing Integrin α5α1 at cell periphery, move rearward with actin flow, and become organized Into linear arrays of nanodomains. Nanodomain arrays are bone fide fibrils. Joining of the additional Fn1 nanodomains to these arrays leads to the generation of longer fibrils as assemblies move toward the cell center. FUD does not Interfere with the formation of Fn1 nanodomains; Instead, It blocks the organization of Fn1 nanodomains into linear arrays. LAMMs – large apparent molecular mass complexes, defined in Zhang et al., 1996.

## Discussion

In this manuscript, we describe data supporting a novel mechanism underlying the process of Fn1 fibrillogenesis. In this model, 6-11 Fn1 dimers assemble into nanodomains containing integrin *α*5*β*1 at the cell periphery, move rearward with actin flow, and become organized into linear arrays of periodically-spaced nanodomains. Joining of the additional Fn1 nanodomains to these arrays leads to the generation of longer and longer fibrils as nanodomain assemblies move toward the cell center. We show that these periodical nanodomain assemblies are bona fide Fn1 fibrils.

The N-terminus of Fn1 is essential for Fn1 fibrillogenesis (Schwarzbauer, 1991). FUD binds tightly to the N-terminal domain of Fn1 (K_D_<2.6 nM), with a fast *k_on_* and a slow *k_off_*, and acts as a competitive inhibitor of Fn1-Fn1 interactions (Ma et al., 2015; Maurer et al., 2010; Tomasini-Johansson et al., 2001). Although in the absence of FUD, individual fibrils in an established matrix are stable and can be tracked for over 16 hours (S.A., unpublished observations), when FUD is added to cells, it localizes with Fn1 fibrils and dismantles the mature Fn1 ECM (Filla et al., 2017; Tomasini-Johansson et al., 2001). These findings suggest that the fibrillar Fn1^+^ ECM is maintained through dynamic interactions mediated by the 70 kDa Fn1 N-terminal domain. Our live imaging experiments demonstrated that in addition to dismantling pre-existing fibrils, FUD effectively blocks their de-novo formation. SMLM showed that FUD does not affect the formation of Fn1 nanodomains, instead, it blocks the organization of Fn1 nanodomains into linear arrays. Together, these data suggest that FUD blocks the dynamic interactions between the N-terminal Fn1 assembly domain and factor(s) linking Fn1 nanodomains into fibrils. One of the factors spanning the nanodomains could be an extended Fn1 dimer because we have observed occasional localizations of Fn1 molecules between nanodomains (e.g., **Figs. 6D** or **8A”**). Rare Fn1 localizations seen between Fn1 nanodomains are likely derived from cellular Fn1 as opposed to plasma Fn1 found in the fetal bovine serum in the complete medium (Table M1 in Methods). This is because bovine Fn1 in the complete medium, albeit recognizable by the 297.1 polyclonal antibodies (**Sup. Fig. 12B**), is not incorporated into Fn1 fibrils at a detectable level (**Sup. Fig. 12A**).

Previously, the periodical distribution of domain-specific antibody epitopes in EM and SMLM images of Fn1 fibrils was explained by a model stipulating that Fn1 fibrils are made of aligned and extended Fn1 molecules (Dzamba and Peters, 1991) (**Sup. Fig. 8A-B**). In EM micrographs of Fn1 molecules sprayed onto mica surface, one can see extended Fn1 dimers that are on average 120-140 nm in length (Engel et al., 1981; Erickson et al., 1981). Therefore, it was hypothesized that spacing of ∼84 nm seen in EM with domain-specific antibodies (Dzamba and Peters, 1991) or spacing of ∼100 nm in SMLM images (Fruh et al., 2015) was due to the overlap between the extended Fn1 molecules at their N-termini (Dzamba and Peters, 1991), e.g., (**Sup. Fig. 8B,** overlap marked by a bubble). The findings that the N-terminal 70 kDa domain is required for Fn1 fibrillogenesis, that this domain can mediate intermolecular Fn1-Fn1 interactions, and that inhibitors of Fn1 fibrillogenesis bind to this domain were interpreted in favor of this model. However, these biochemical data are also consistent with a very different model of Fn1 fibrillogenesis (**Sup. Fig. 8**), one that was proposed by (Tomasini-Johansson et al., 2006) and discussed below.

A canonical model poses that Fn1 fibrillogenesis entails intermolecular Fn1-Fn1 interactions mediated by the 70 kDa N-terminal domain of Fn1 (Hocking et al., 1994; Zhong et al., 1998), summarized in (Singh et al., 2010). These interactions were hypothesized to be important for linking extended and periodically-aligned Fn1 dimers into fibrils (e.g., **Sup. Fig. 8A**). It was stipulated that FUD blocks Fn1 fibrillogenesis because of its ability to compete with Fn1-Fn1 intermolecular interactions and disassemble fibrils by inhibiting intermolecular interactions between Fn1 dimers (**Sup. Fig. 8D**).

However, these biochemical and cell biological experiments are also consistent with a different model, wherein Fn1 fibrillogenesis is mediated by the binding of the 70 kDa N-terminal domain to as of yet an unidentified factor(s). Examples of such factors could be tissue transglutaminase-2 (Akimov et al., 2000; Yuan et al., 2007) or large apparent molecular mass cell-surface complexes (LAMMs) (Tomasini-Johansson et al., 2006; Zhang and Mosher, 1996) (**Sup. Fig. 8E**). Thus, as an example, FUD can disrupt fibrillogenesis by disrupting the interactions between the 70 kDa Fn1 assembly domain and LAMMs (**Sup. Fig. 8E**). Biochemical evidence supporting this model is multifold: 1) isolated 70 kDa N-terminal domain can bind to Fn1-null cells in the absence of Fn1, forming short linear arrays; this binding is time- and dose-dependent, and is detectable with as little as 5 nM of 70 kDa N-terminal segment of Fn1 (Tomasini-Johansson et al., 2006); 2) Incubation of the 70 kDa domain of Fn1 with cells precipitates LAMMs in pull-down assays (Zhang and Mosher, 1996); 3) LAMMs contain trypsin-sensitive protein(s) (Zhang and Mosher, 1996); and 4) FUD inhibits the binding of 70 kDa N-terminal fragment to Fn1-null cells (Tomasini-Johansson et al., 2006). These findings suggest that Fn1 nanodomains in fibrils can be held together by the interactions between the 70 kDa N-terminal assembly domain of Fn1 and LAMMs (**Sup. Fig. 8F1**). However, at this time, we cannot rule out the possibility that Fn1 nanodomains are held by an extended Fn1 molecule via N-N-terminal interactions (**Sup. Fig. 8F2**), or that a combination of the latter two models may be true.

Our model in which Fn1 fibrils are composed of periodically-spaced nanodomains containing multiple Fn1 dimers is also supported by electron microscopy and atomic force microscopy studies done in the past: 1) periodical staining of Fn1 fibrils was observed in EM with polyclonal antibodies raised to entire human plasma Fn1 (Furcht et al., 1980a); 2) grapes- on-a-vine appearance of immuno-gold complexes detecting plasma Fn1 in fibrils in electron micrographs (Peters et al., 1990); 3) bulbous appearance of Fn1 fibrils in cryo-scanning transmission electron tomography images (Lansky et al., 2019); and 4) beaded appearance of Fn1 fibrils detected by atomic force microscopy (Gudzenko and Franz, 2015). The latter two methods did not rely on antibody staining to detect Fn1 nanodomains, supporting the notion that the beaded architecture of Fn1 fibrils is not an artifact of antibody labeling.

Together with our data, these experiments suggest a new model of Fn1 fibrillogenesis (**Fig. 10E**). In this model, Fn1 nanodomains containing multiple Fn1 dimers form at cell periphery. These nanodomains may be similar to focal complexes or adhesion-related particles seen by cryo-electron tomography by (Patla et al., 2010). The centripetal translocation of Fn1 nanodomains is coordinated with their organization into linear arrays, giving rise to focal adhesions composed of ∼3-5 Fn1 nanodomains and long Fn1 fibrillar adhesions composed of dozens of nanodomains. Fn1 fibrils generated in this process are then incorporated into Fn1 ECM (Mautner and Hynes, 1977; Pankov et al., 2019; Sivakumar et al., 2006). Two pieces of evidence suggest that Fn1 nanodomains in fibrils are linked: 1) the preservation of the linear organization and the nanoarchitecture of Fn1 fibrils after the treatment of cells with DOC which dissolves cell membranes, and 2) the presence of fibrous material between immunogold densities in electron micrographs (Chen et al., 1997; Dzamba and Peters, 1991; Furcht et al., 1980a; Furcht et al., 1980b; Furcht et al., 1980c; Lansky et al., 2019; Peters et al., 1990). The sparsity of Fn1 localizations between Fn1 nanodomains in fibrils suggests that molecules other than or in addition to Fn1 participate in the potential linking of Fn1 nanodomains into periodical nanodomain arrays.

The beaded architecture of Fn1 ECM has important implications for the mechanisms of ECM formation, remodeling, and signal transduction. The tensile strength of knotted strings is significantly lower than that of strings with uniformly-aligned fibers (Arai et al., 1999; Saitta et al., 1999), thus the beaded architecture of Fn1 fibrils may facilitate their rupture under strain (Ohashi et al., 1999). The non-uniform, nanodomain architecture of Fn1 may facilitate the accessibility of Fn1 fibrils to matrix metalloproteases. In this model, degradation of Fn1 fibrils by metalloproteases may be accomplished by the cleavage of LAMMs between Fn1 nanodomains facilitating dynamic ECM remodeling. Finally, Fn1 is known to bind growth factors (Martino and Hubbell, 2010; Saunders and Schwarzbauer, 2019; Wijelath et al., 2002), and cell adhesion to ECM is known to orchestrate growth factor signaling (Hynes, 2009). Thus, Fn1 nanodomains could serve as platforms for the binding and presentation of concentrated packets of growth factors to cells, the organization of Fn1 nanodomains into closely-spaced arrays could further facilitate clustering and signaling by growth factor receptors.

## Acknowledgments

We thank Richard Hynes and Nathan Astrof for insightful discussions and careful reading of the manuscript, Sydney Astrof for encouragement and help with data entry, Patrick Murphy for endothelial cells, Richard Hynes for the gift of 297.1 antibody, Tung Chan for help with setting up Western Blotting using ProteinSimple, and Jonas Ries for advice that helped us improve SMLM resolution of our imaging, and for his help with adopting the SMAP software. Epitope mapping of 297.1 antibody was performed by PEPperPRINT GmbH, Heidelberg, Germany.

## Materials and Methods

### Generation Fn1-fluorescent protein targeting constructs

Sequences of monomeric (m) green fluorescent protein (GFP), mNeonGreen, mScarlet-I, and tdTomato were obtained from FPbase (https://www.fpbase.org). The sequence encoding one of the above fluorescent proteins (FPs) was knocked into the Fn1 locus following the last coding exon of mouse Fn1, and separated from the last coding amino acid by a flexible, proline-rich linker, PPPELLGGP (Snitkovsky and Young, 1998). Targeting was achieved by CRISPR/Cas9 (Ran et al., 2013). The sequence of the guide RNA was chosen and off-target sites were identified using GuideScan and Off-Spotter software (Perez et al., 2017; Pliatsika and Rigoutsos, 2015). The guide RNA (gRNA) sequence 5’-AGC GGC ATG AAG CAC TCA AT-3’ targeting the last coding exon of *Fn1* was subcloned downstream the U6 promoter into the PX459 vector (Addgene, cat # 62988) encoding the Cas9-2A-Puromycin cassette (Ran et al., 2013). The homology-directed repair (HDR) template was constructed using pBS-KS vector (Sup. Fig. 1a). The sequence of the last coding exon of *Fn1* 5’- AACGTAAATTGCCCCATTGAGTGCTTCATGCCGCTAGATGTGCAAGCTGACAGAGACGATTCTCGAGAG-3’ was modified to 5’- AACGTAAATTGCCCCAT<u>c</u>GA<u>a</u>TGCTTCATGCCGCTAGATGTGCAAGCTGACAGAGACGATTCTCGAGAG-3’ in the HDR template by introducing silent mutations (underlined) to prevent targeting of the template by the gRNA. Homology arm 1 contained 677 bp encoding exon #45, intron, and a portion of the last exon (#46), of the transcript *ENSMUST00000055226.12*. Homology arm 2 encoded 1739 bp immediately downstream of the *Fn1* termination codon and included the unmodified 3’UTR of Fn1. Knockin Fn1^mEGFP/+^ mice were generated by Biocytogen using the same HDR construct and a longer gRNA, 5’-<u>T</u>AG CGG CAT GAA GCA CTC AAT <u>G</u>G-3’, targeting the same sequence in the last coding exon (differences between the two gRNAs are underlined). Targeting was confirmed by sequencing and Southern Blotting (Sub. Fig. 1b). 500 bp around each of the top ten predicted off-target sites were sequenced and no mutations were found in the founder mice. Mice containing correctly-targeted *Fn1* locus were used to establish living colonies of Fn1^mEGFP/mEGFP^ animals. Wild-type, Fn1^mEGFP/+^, and Fn1^mEGFP/mEGFP^ mice were genotyped using the following primers Fn1-WT-Fwd 5’- TCCCCGAAACACACACACTTTTGGT-3’, Fn1-WT-Rev 5’ GTCACCCTGTTCTGCTTCAGGGTTT-3’, and Fn1GFP-Rev 5’- GACCCGCGCCGAGGTGAAG-3’. Wild type band is Fn1-WT-Fwd and Fn1-WT-Rev give rise to 372 bp for the wild-type allele; Fn1-WT-Fwd and Fn1GFP-Rev primer located in the GFP sequence give rise to 512 bp if the targeted allele is present. Mice were housed in an AAALAC-approved barrier facility. All experimental procedures were approved by the Institutional Animal Care and Use Committee of Rutgers University and conducted in accordance with the Federal guidelines for the humane care of animals.

### Cell Culture

**Table M1.**
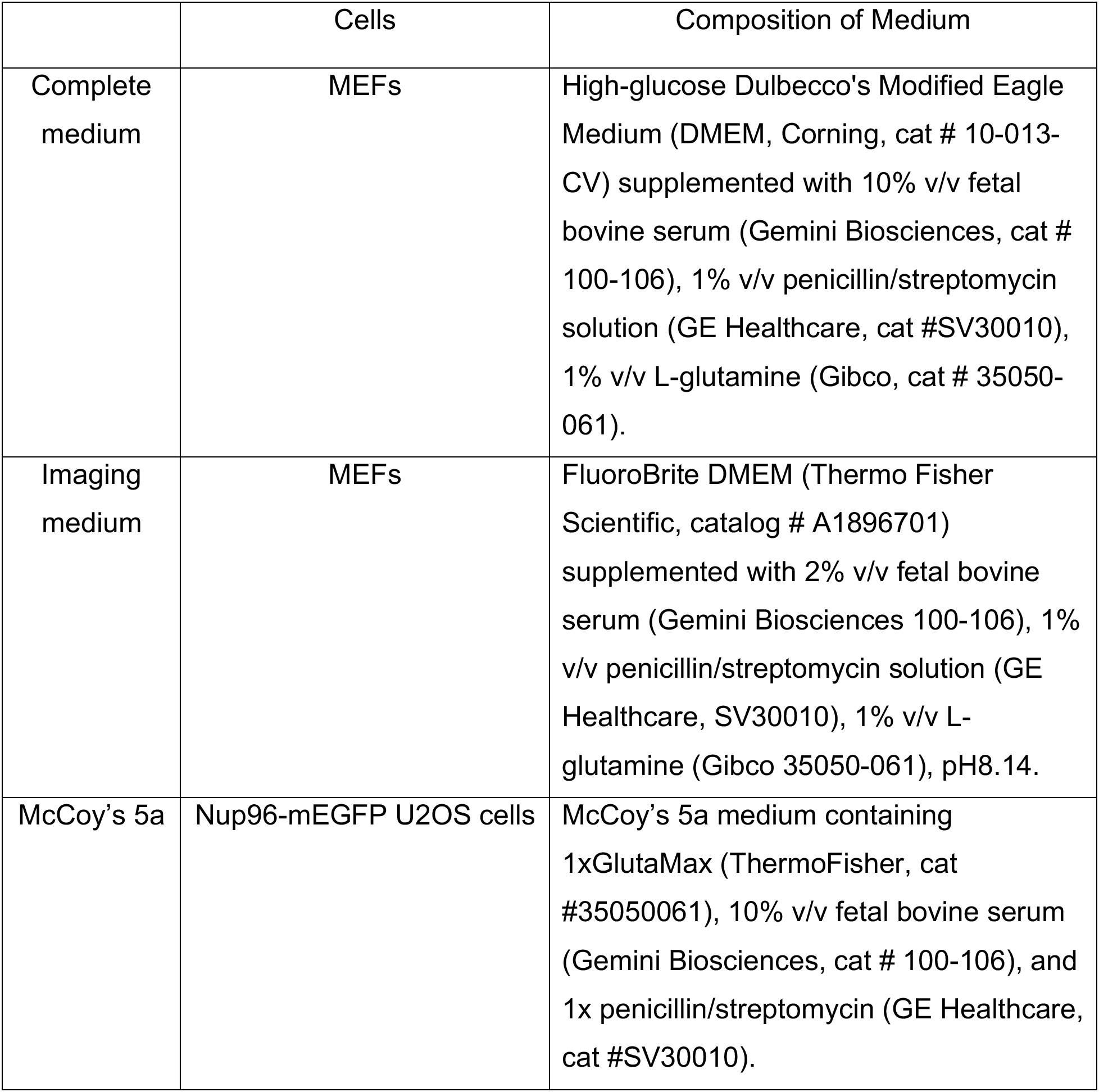
Media.

#### Cells

Wild-type mouse embryonic fibroblasts (MEFs) were isolated from embryonic day (E) 13.5 embryos derived from the C57BL/6J strain (Jackson Labs, stock # 664) according to established protocols (Behringer et al., 2014) and cultured in complete medium. For propagation, MEFs were plated in flasks pre-coated with 0.1% gelatin solution and grown in complete medium at 37°C, 5% CO_2_.

Fn1^flox/+^;Rosa^mTmG/+^ MEFs were isolated from E13.5 embryos, as above, and express membrane-bound tdTomato and wild-type Fn1 in the absence of Cre recombinase (MuzumdarIRED-GFP virus according to the manufacturer’s recommendations (Vector Biolabs, #1710). In this experiment, Cre-recombinase is expressed transiently and mediates site-specific recombination between a pair of loxP sites flanking the 1^st^ exon of Fn1 (Sakai et al., 2001) and another pair of loxP sites flanking the STOP cassette in the mTmG reporter (Muzumdar et al., 2007). Three days following infection with Ad-Cre-IRED-GFP, GFP+ cells were sorted resulting in a pure population of Fn1-null MEFs (confirmed by Western Blotting, data not shown, and immunofluorescence, **Sup. Fig. 12**). Fn-null MEFs were culture on gelatin-coated dishes in complete medium.

Fn1-FP cells (except Fn1-mEGFP cells) were generated by CRISPR/Cas9 mutagenesis of wild-type MEFs. CRISPR/Cas9 targeting was performed by transfecting wild-type MEFs using the PX459 plasmid encoding Fn1 gRNA and the HDR template using lipofectamine 3000, as described (Ran et al., 2013). MEFs expressing Fn1-mEGFP proteins used for live imaging and STORM were generated from homozygous E13.5 Fn1^mEGFP/mEGFP^ or, when noted from Fn1^mEGFP/+^ embryos. Fn1^mEGFP/mEGFP^ MEFs were used to quantify Fn1 molecule number in nanodomains, measure nanodomain spacing, and nanodomain diameter, as well as in experiments to label both the N- and the C-termini of Fn1. For live imaging, Fn1^mEGFP/+^ were used. Fibronectins from Fn1^mEGFP/mEGFP^ and Fn1^mEGFP/+^ cells behave equivalently in Fn1 matrix assembly assays (data not shown).

Nup96-mEGFP cells ((Thevathasan et al., 2019), clone #195, Cell Line Services, CLS, clsgmbh.de, catalogue no. 300174)) were cultured according to the vendor’s specifications in McCoy’s 5a medium containing 1xGlutaMax (ThermoFisher, cat #35050061), 10% v/v fetal bovine serum (Gemini Biosciences, cat # 100-106), and 1x penicillin/streptomycin (GE Healthcare, cat #SV30010).

### Reagents and buffers

FUD and III-11C peptides were generated as described (Sottile and Chandler, 2005; Tomasini-Johansson et al., 2001) and stored in PBS at −80° C. 4% DOC solution was prepared by dissolving 0.4 g deoxycholate salt (Sigma, catalog # D6750) in 10 ml of imaging medium; the solution was then vortexed and filter sterilized. The pH of the final solution was 8.01. 16% paraformaldehyde (PFA) (Electron microscopy Sciences; catalog # 50-980-487) was diluted in 1x PBS to prepare 4% PFA. The 4% PFA solution was aliquoted into 1 ml microfuge tubes, stored at −80° C, and thawed at 37° C immediately before use.

1X phosphate buffered saline pH 7.5 (PBS) was prepared from 10X PBS (VWR, catalog # 76180-740). PBST was prepared using Triton X-100 (Sigma-Aldrich, catalog # T-8787) and contained 0.1% Triton for all stainings except those involving Nup96-mGEFP cells, as detailed below. Blocking buffer was prepared by adding 10% Donkey serum (Sigma-Aldrich, catalog # D9663) to 1X PBST; 5 mg/ml stock of DAPI (Fisher Scientific, cat #D3571) was prepared in water and used at 1:300 dilution. Stain Buffer (cat # 554656 BD Pharmingen) was used for antibody dilutions and washing of cells that were stained without permeabilization. Hoechst 33342 Trihydrochloride (Thermo Fisher, catalog # H1399, 10mg/ml) was used for labelling live MEFs at 1:300 dilution. In live MEFs, F-actin was labelled using SiR actin (cat# CY-SC001 used at 1 μM final concentration). mCardinal-Lifeact-7 was a gift from Michael Davidson (Addgene plasmid # 54663 ; http://n2t.net/addgene:54663 ; RRID:Addgene_54663). Vectashield antifade mounting medium (Vectorlabs, catalog # H-1000) was used for cover slipping.

For STORM imaging, we used 25 mm high precision glass cover slips #1.5H (Marienfeld ref# 0117650, obtained from Azer Scientific, PA, USA, cat # ES0117650) without coating. Prior to their use, glass cover slips were cleaned using concentrated nitric acid, washed in water, air dried, and autoclaved, as described in (Kaech and Banker, 2006). Clean coverslips were stored in sealed 6-well plates for no longer than a week prior to their use.

GLOX/BME STORM buffer contained 50 mM Tris-HCl (Fisher Scientific, catalog # T-395-1), pH 8.0, 10 mM NaCl (Sigma-Aldrich, catalog # S-7653), 10% glucose (Sigma-Aldrich, catalog # G8270), 0.5 mg/ml glucose oxidase (Sigma-Aldrich, catalog # G2133), 40 μg/ml catalase (Sigma-Aldrich, catalog # C40), and 143 mM β-mercaptoethanol (*β*ME, Sigma-Aldrich, cat# 444203) (Thevathasan et al., 2019). Stocks of enzyme solutions were prepared and stored at - 20°C as described in (Jimenez et al., 2020). GLOX/BME buffer was used for STORM imaging of cells plated on coverslips.

GLOX/MEA STORM buffer was used for STORM imaging of cells plated in Ibidi glass bottom 8-well chambers (catalog # 80827). This buffer was prepared as above, but instead of *β*ME, it contained 50 mM mercaptoethylamine (MEA, Sigma-Aldrich, catalog # 30070) (Jimenez et al., 2020). For double-color STORM imaging, we used the SMART Kit buffer (Abbelight).

### Antibodies

All primary antibodies were checked for specificity on cells that were genetically-null for the antigen (e.g., **Sup. Fig. 12**) and tissues: Fn1-null tissue sections obtained from Fn1-null embryos were used to assay the specificity of each of the anti-Fn1 antibodies; Tissues isolated from GFP-null, Itga5-null, and mCherry-null embryos were used to authenticate the specificity of anti-GFP, anti-Itga5, and anti-mCherry antibodies. For each of the antibodies, staining of control tissues resulted in no more fluorescent signal than the background fluorescence produced by the use of secondary antibodies only. Rabbit polyclonal antibody R457 was raised against the 70 kDa N-terminal domain of Fn1 (Aguirre et al., 1994; Sechler et al., 2001) and rabbit polyclonal R184 was raised against the first six type III repeats of Fn1 (Raitman et al., 2018). The specificity of these antibodies was verified by ELISA and Western blotting reporter in the references in Table M2.

**Table M2.**
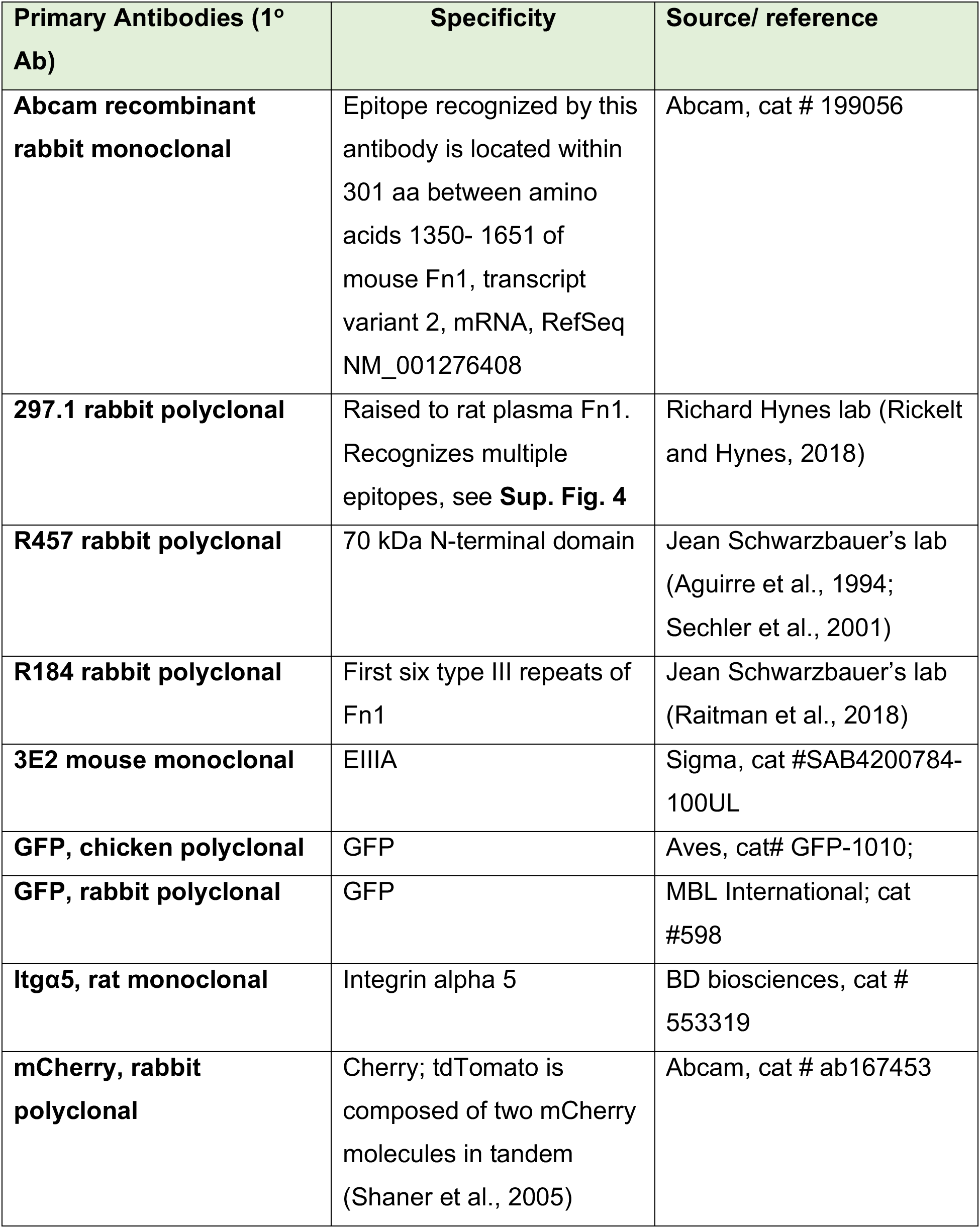
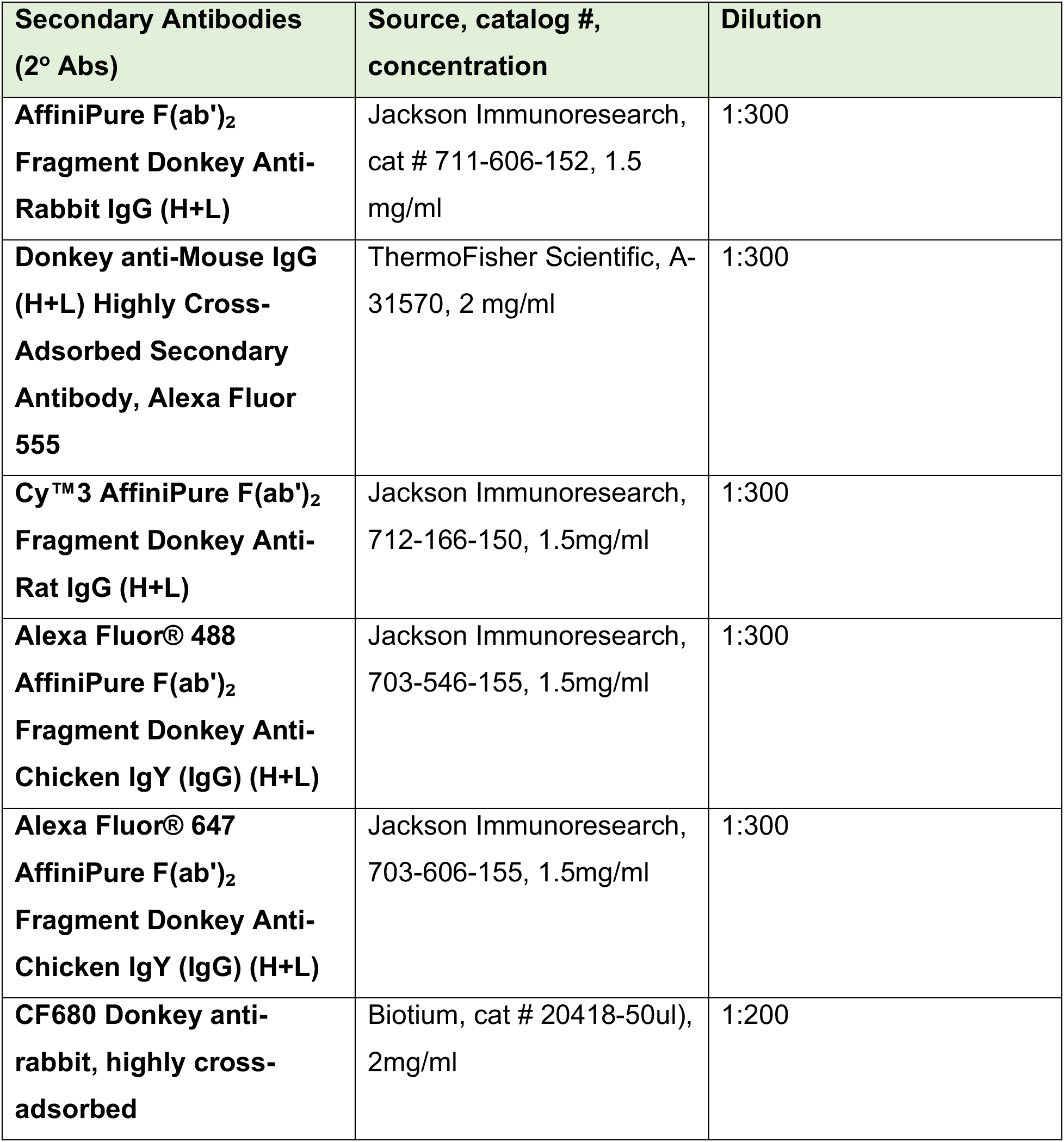
Antibodies.

#### Epitope mapping of rabbit 297.1 polyclonal antibody

Epitopes recognized by the polyclonal 297.1 antibody were mapped by generating custom overlapping peptide arrays (PEPperPRINT GmbH, Heidelberg). Fn1 protein sequence including alternatively-spliced exons was encoded by 15-amino acid peptides with a peptide-peptide overlap of 13 amino acids. The resulting Fn1 peptide microarrays contained 1,239 different peptides printed in duplicate (2,478 peptide spots), and were framed by additional HA-tag (YPYDVPDYAG, 106 spots) control peptides. HA-tag peptide was used to monitor array quality and served as a positive control for clean antibody binding detected with anti-HA-tag antibody (mouse monoclonal anti-HA DyLight800 used at 0.5 µg/ml concentration). Prior to staining, arrays were incubated with blocking buffer (Rockland, cat # MB-070) for 30 min at RT. To measure background antibody binding, arrays were first incubated with secondary goat anti-rabbit IgG (Fc) DyLight680 at 0.2 µg/ml concentration diluted in PBS, pH 7.4, containing 0.05% Tween 20 and 10% blocking buffer, and imaged. To determine specific 297.1 antibody binding, arrays were incubated for 16 hours with shaking at 4°C with two dilutions of 297.1 antibody (1:300 and 1:1000). Antibody dilutions were made in PBS, pH 7.4 containing 0.05% Tween 20 and 10% blocking buffer (Rockland, cat # MB-070). Arrays were then washed with PBS, pH 7.4 containing 0.05% Tween 20 and incubated with goat anti-rabbit IgG (Fc) DyLight680 (0.2 µg/ml) for 45 min at RT. Arrays were then washed and imaged using LI-COR Odyssey Imaging System; scanning offset 0.65 mm, resolution 21 µm, scanning intensities of 7/7 (red = 700 nm/green = 800 nm). Quantification of spot intensities and peptide annotation were done with PepSlide® Analyzer.

#### Embryo Staining

E9.5 embryos were isolated either from matings of wild-type C57BL6/J mice or from mating Fn1^mEGFP/mEGFP^ mice with wild-type mice to obtain Fn1^mEGFP/+^ embryos. Embryos were fixed using cold 4% PFA overnight at 4°C, washed 3 x 5 min in 1xPBS, and blocked overnight at 4°C in blocking buffer containing PBS, 0.1% Triton-X, and 10% donkey serum. Embryos were stained either with Abcam monoclonal anti-Fn1 antibodies at 1:300 dilution in blocking buffer or with both Abcam monoclonal anti-Fn1 antibodies and anti-GFP antibodies at 1:300 dilution. The staining procedure was executed exactly as described in (Ramirez and Astrof, 2020).

#### Analysis of Fn1 matrix assembly and Western Blotting using MEFs

Matrix Assembly was performed according to the established protocols (Wierzbicka-Patynowski et al., 2004). MEFs were plated in 6-well dishes (9 cm^2^ growth area) at a density of 2 × 10^5^ cells per well for 48 h, in complete medium and incubated under sterile conditions at 37°C, 5% CO2. Cells were washed twice with ice cold PBS (supplemented with Mg^2+^ and Ca^2+^), scraped with a cell scraper and lysed with either 500 μl RIPA lysis buffer pH 8.0 (50 mM Tris-Cl, 150 mM NaCl, 2 mM EDTA, 1% v/v NP-40, 0.5% w/v sodium deoxycholate, 0.1% w/v SDS, 1X protease inhibitor cocktail (Cell Signaling Technology, cat # 5871), or DOC lysis buffer, pH 8.8 (20 mM Tris-Cl, 2 mM EDTA, 2% w/v sodium deoxycholate, 1X protease inhibitor cocktail (Cell Signaling Technology, 5871). Extracts were carefully transferred to Eppendorf tubes containing 1 μl (250 units) Benzonase® Nuclease (Sigma-Aldrich, E1014), mixed by inverting a few times and incubated at 37 °C for 15 mins. The samples were then centrifuged at 16,000 × g for 15 min at 4 °C. For cells lysed with DOC lysis buffer, the supernatant containing DOC-soluble material was carefully removed, and the pellet containing the DOC-insoluble material was resuspended in 100 μl SDS solubilization buffer, pH8.8 (20 mM Tris-Cl, 2 mM EDTA, 1% w/v SDS, 1X protease inhibitor cocktail (Cell Signaling Technology, 5871). The DOC-insoluble pellet was thoroughly dissolved by heating the sample to 95 °C and vortexing. All samples were aliquoted and stored at −80 °C until further use. Prior to quantification of Fn1 in the samples, the total protein concentration of the RIPA and DOC lysates was determined using the BCA protein assay (Pierce™ BCA Protein Assay Kit, 23225). Lysates containing Fn1 and Fn1-FP fusion proteins were reduced an resolved using 66-440 kDa Wes separation module (ProteinSimple, SM-W007). Primary antibodies were used at the following dilutions: anti-total Fn1 – 1:1000 (Abcam, ab199056), anti-GFP – 1:1000 (Roche, 11814460001), anti-mCherry – 1:1000 (Abcam, 167453). Primary antibodies were detected using horseradish peroxidase-conjugated secondary antibodies (anti-Rabbit Detection Module ProteinSimple, DM-001), and chemiluminescence was quantified using the Compass for SW software (v3.1.8). Prior to running experimental samples, care was taken to optimize the dilutions of lysates to be within the linear range of the detection.

#### Analysis of Fn1 matrix assembly and Western Blotting using E9.5 embryos

To analyze Fn1-mEGFP matrix assembly *in vivo*, we mated Fn1^mEGFP/+^ mice to obtain wild-type, Fn1 ^mEGFP/+^ and Fn1^mEGFP/mEGFP^ littermate embryos. At the time of dissection, each embryo was frozen on dry ice in individual Eppendorf tubes and stored at −80°C until the genotyping was complete. Yolk sacs were used for genotyping. 300 μl of DOC ice-cold lysis buffer pH 8.8 containing protease inhibitors (see recipe above) were added to each embryo, and the embryos were dissociated by passing through a 27-gauge syringe needle 5 times, keeping the tubes on ice. Lysates were centrifuged at 16,000 × g for 15 min at 4°C. The supernatant containing DOC-soluble material was transferred to another tube and was supplemented with 100 μl of 4x SDS loading buffer and *β*-mercaptoethanol (*β*ME), at a final concentration of 350 mM of *β*ME. The DOC-insoluble pellet was washed twice with 300 μl of DOC lysis buffer on ice, and the pellet was then resuspended with 200 μl SDS solubilization buffer with protease inhibitors (see recipe above). The pellet was dissolved by heating at 95°C and vortexing. And supplemented with 66.6 μl of 4x SDS loading buffer and *β*-mercaptoethanol (*β*ME), at a final concentration of 350 mM of *β*ME. Doc soluble and insoluble samples were heated at 95°C for 5 min and 40 μl were loaded on 4-12% acrylamide gel (Invitrogen cat # XP04120BOX). Fn1 was detected using rabbit anti-Fn1 1° antibody (Abcam, cat # 199056) and IRDye 680RD Donkey anti-Rabbit 2° antibody (Licor, cat # #926-68073).

Membranes were imaged using Li-Cor Odyssey 9120 Gel Imaging System (#ODY-2425) and quantified using Fiji software.

#### Testing the reactivity of 297.1 polyclonal antibody to bovine Fn1 present in the fetal bovine serum

To test whether 297.1 antibody can bind bovine Fn1, we used a complete medium (CM, see Table M1) and, as a control, a conditioned completed medium (CCM) which we prepared by incubating 2×10^5^ cells plated in a well of a 6 well plate for 48 hrs with 2 mL of CM. CCM was centrifuged at 1000 rpm for 3 minutes to discard dead cells. 100 μL of trichloroacetic acid (TCA) were added to 900 μL of CM or CCM to precipitate the protein and incubated in ice for 30 min. Samples were centrifuged at 14000 rpm at 4C for 15 min. Pellets were washed with 700 μL of 100% acetone and then resuspended in a gel loading buffer containing 100 μl of 0.1N NaOH, 75 μl of 4X SDS-PAGE loading buffer, 0.35 M *β*ME and 125 μl of H20. Samples were heated at 95°C for 5 min and 10 μl of each sample were resolved using Novex™ WedgeWell™ 4 to 12%, Tris-Glycine, 1.0 mm, Mini Protein Gel (Invitrogen cat # XP04120BOX) and run using Tris-Gly SDS Running buffer (Invitrogen, cat # LC2675). Following the transfer to nitrocellulose membrane (Bio-Rad, cat # 1620122), Fn1 was detected by immunoblotting using 1:1000 dilution of 297.1 polyclonal antibodies and IRDye 680RD Donkey anti-Rabbit secondary antibody (Licor, cat # #926-68073). Protein standards were from Invitrogen, cat # LC5925. Membranes were imaged using Li-Cor Odyssey 9120 Gel Imaging System (#ODY-2425)

#### Coating of coverslips with different ECM proteins

#1.5 round glass coverslips (Electron Microscopy Sciences. Catalog # 72230-01) were coated with the following ECM proteins when noted: gelatin (Sigma Aldrich, catalog # G2500) (0.1% (w/v) of gelatin was prepared in Milli-Q water and autoclaved to dissolve), vitronectin (Sigma Aldrich, catalog # SRP3186; stock solution was prepared as 200µg/ml in 0.1% BSA and water) and laminin (R&D systems, catalog # 3400-010-02, stock 1 mg/ml was pipetted into 10ul aliquots and stored at −80°C). To coat with gelatin, glass surfaces were incubated with the 0.1% gelatin solution for 5 min at room temperature (RT). To coat with vitronectin or laminin, glass surfaces were incubated at 37° C for 1 hr in 20 μg/ml of either vitronectin or laminin, excess liquid was removed, cover slips were rinsed once with 1X PBS, and blocked with 10 μg/ml heat denatured BSA for 30 min before plating cells (Lu et al., 2020).

#### MEFs plated on different substrata

MEFs were grown either on #1.5 round glass coverslips in 24-well dishes or in 8-well glass Ibidi dishes depending on the experiment for the times indicated in figure legends. MEFs were then rinsed with 1X PBS (warmed to 37^0^ C) for 5 min, fixed with freshly thawed 4% PFA pre-warmed to 37^0^ C for 20 min, and washed three times with 1X PBS (warmed to 37^0^ C) with mild shaking. All subsequent washing steps were done with shaking. For permeabilization, cells were washed once in 1X PBS containing 0.1% Triton-X 100 (PBST). Blocking was done for 30 min in 10% Donkey serum prepared in PBST (blocking solution). After blocking, cells were incubated in primary antibodies diluted in blocking solution overnight at 4^0^ C, as specified in **Table M2**. This was followed by 3 washes in PBST for 10 min each. Cells were then incubated with secondary antibodies diluted in PBST for 60 min at RT. Finally, cells were washed three times with PBST for 10 min each. DAPI (1:300) was added to the second wash. Cells were mounted using Vectashield.

### Hydrogels

#### Methacrylated Alginate Synthesis

Methacrylated alginate (MeAlg) was synthesized according to a previously established protocol (Khetan et al., 2013). In brief, alginic acid sodium salt from brown algae (Sigma-Aldrich, USA) (3% w/v) was fully dissolved in Dulbecco’s phosphate buffered saline (dPBS, Sigma-Aldrich, USA). Then, methacrylic anhydride (Sigma-Aldrich, USA) (8% v/v) was added drop-wise to the alginate solution and stirred for 12 h at 4°C, using 2M NaOH (Sigma-Aldrich, USA) to ensure that the pH remained between 8 and 9 for the duration of the reaction. The resulting solution was passed through filter paper (GE Whatman) and poured into Spectra/Por dialysis membrane with a 6–8 kDa molecular weight cutoff (Fischer Scientific) and kept in DIW under stirring for 7 days to eliminate the unreacted MA and salts. Dialyzed solution was then freeze-dried for 4 days to obtain MeAlg foam.

#### Fabrication of the Hydrogel Substrates

MeAlg substrates were fabricated using a previously established protocol (Guvendiren and Burdick, 2012). Briefly, petri dishes with glass bottoms were treated with UV/ozone (UVO) for 30 minutes, immediately followed by a coating of 3- (trimethoxysilyl)propyl methacrylate (TMS) (Sigma-Aldrich, USA) to methacrylate the glass surfaces (Guvendiren et al., 2009). The dishes were left in a desiccator overnight. The hydrogels were fabricated using Michael-type addition polymerization. First, 2-hydroxy-4’-(2- hydroxyethoxy)-2-methylpropiophenone (I2959) (Sigma-Aldrich, USA), a photoinitiatior (0.5% w/v) was completely dissolved in Dulbecco’s PBS (dPBS), followed by the lyophilized MeAlg (3% w/v) synthesized previously. This was kept at room temperature until a clear solution was achieved. Crosslinking occurs with the introduction of DL-Dithiothreitol (DTT) (Sigma-Aldrich, USA) to the solution, along with 0.2M triethanolamine (Sigma-Aldrich, USA) at pH 10. To form 3kPa and 12 kPa gels, 20% and 30% (w/v) DTT are used, respectively. To promote cell adhesion, GRGDSPC peptide (1% w/v) (Genscript) was added to the solution. After all contents were thoroughly mixed, 5μL of MeAlg solution was pipetted onto the surface of the dish before being covered with a glass coverslip in order to create gels less than 30μm thick. These were left at room temperature for an hour to crosslink before being submerged in dPBS to remove the coverslip.

#### Atomic Force Microscopy

For stiffness measurements, hydrogel samples were submerged in PBS and placed in a Dimension Icon AFM with ScanAsyst (Bruker). Using the PeakForce-QNM mode, hydrogel samples were indented using an MLCT-Bio probe tip with pyramidal geometry (Bruker, CA) and a nominal spring constant of 0.03 N/m, checked by thermal calibration.

#### Treatment of cells with Deoxycholate (DOC)

10^4^ Fn1^mEGFP/+^ MEFs were plated for 48 hrs in 8-well glass bottom Ibidi dishes in complete medium and incubated at 37° C and 5% CO_2_. Two hours before imaging SiR-actin was added at 1 μM final concentration. SiR-actin contains a far-red dye, silicon rhodamine, conjugated to jasplakinolide that labels F-actin in live and fixed cells (Lukinavicius et al., 2014). Just before imaging, complete medium was replaced by 150 μl imaging medium containing 33 μg/ml of Hoechst 33342. Positions were marked in each well and live imaging was initiated at 37°C and 5 % CO_2_ humidified chamber. After 15 min, 150 μl 4% DOC solution prepared in imaging medium containing 33 μg/ml Hoechst was added to the experimental well (final pH 8.01) and 150 μl imaging medium containing 33 μg/ml Hoechst but without DOC was added to the control well. Cells were imaged at 50 sec intervals until F-actin and DNA disappeared (see Movie 4). The medium was then removed, cells were rinsed for 1 min with 1X PBS pre-warmed to 37°C, fixed with 4% PFA pre-warmed to 37°C. For staining, cells were permeabilized and blocked as above, and then incubated with Abcam monoclonal anti-Fn1 antibody diluted 1:300 dilution in the blocking solution at 4°C overnight. Primary antibodies were detected with anti-rabbit secondary antibodies conjugated to AlexaFluor647. Enhanced-resolution imaging was used to image the DOC-treated fibrils, as described below.

#### Confocal settings for enhanced resolution imaging

Confocal images of fixed samples were recorded using Nikon A1-HD25 inverted confocal microscope equipped with CFI Apochromat TIRF 100xC Oil objective with the pinhole set to 0.8 Airy units, and imaged through 2 – 4 microns with step size of 0.125 μm - 0.15 μm at a sampling of 40 nm per pixel and 180 nm optical resolution. Deconvolution was done using Nikon 3D deconvolution software (v5.11.01). Airyscan imaging was performed using Zeiss LSM 880 fitted with a 32 array AiryScan GaAsP-PMT detector and the Plan Apochromat 63X Oil (NA 1.4) objective. Deconvolution and pixel reassignment were done using Zeiss LSM software.

#### Quantification of effective labeling efficiency and the number of Fn1 molecules in Fn1 nanodomains

To quantify the number of Fn1 molecules per nanodomain and to measure effective labeling efficiency, we used Nup96-mEGFP U2OS cells as a reference cell line and SMAP software (Ries, 2020; Thevathasan et al., 2019). In Nup96-mEGFP cells, two copies of the nucleoporin Nup96 gene are tagged with the monomeric enhanced green fluorescent protein (mEGFP) at the C-terminus of the Nup96 gene, generating Nup96-mEGFP fusion protein (Thevathasan et al., 2019). The stoichiometry of Nup96-mEGFP in nucleopores is well characterized, and careful measurements, imaging methodology, and software have been developed (Diekmann et al., 2020; Ries, 2020; Thevathasan et al., 2019). Together, these tools allow the use of Nup96-mEGFP cell line as a reference to assess the quality of SMLM imaging protocol, measure effective labeling efficiency, and determine the number of Fn1- mEGFP molecules in Fn1 nanodomains.

Nup96-mEGFP cells (clone #195, Cell Line Services, CLS, clsgmbh.de, catalogue no. 300174) were cultured according to the vendor’s specifications in McCoy’s 5a medium. Fn1-mEGFP MEFs were cultured in complete DMEM as described above. 3×10^5^ Nup96-mEGFP cells and 5×10^4^ Fn1-mEGFP cells were plated in their specified culture medium on 25 mm high precision glass cover slips #1.5H (Marienfeld ref# 0117650, obtained from Azer Scientific, PA, USA, cat # ES0117650) positioned in 6-well plates (Corning, cat # 353046). Prior to their use, glass cover slips were cleaned using concentrated nitric acid, washed in water, and autoclaved, as described in (Kaech and Banker, 2006). Coverslips were used without coating. 24 hours after plating, cells were fixed and stained as described in (Thevathasan et al., 2019) with minor modifications. Cover slips with Nup96-mEGFP cells and Fn1-mEGFP cells were handled contemporaneously in pairs at each step. For fixation, permeabilization and washing, coverslips were kept in 6-well plates. Cells were fixed in PBS containing 2.4% PFA for 20 min at RT. PFA was aspirated and cells were incubated with 100mM NH_4_Cl in PBS for 5 min at RT, and then washed 3 x 5 min in PBS with agitation. Cells were permeabilized for 20 min at RT using PBS containing either 0.2% Triton-X for Nup96-mEGFP cells or 0.1% Triton-X for Fn1- mEGFP. Cells were then washed in PBS 3 x 5 min at RT, and washed once more for 5 min in PBS containing 0.1% Triton-X, and then either used immediately for staining, or stored at 4°C in PBS containing 0.1% Triton-X (PBST) until further use. For each experiment, sufficient amounts of each solution were prepared such that Nup96-mEGFP and Fn1-mEGFP cells were incubated with the same mixtures. Solutions containing blocking reagents and antibodies were spun for 5 min at top speed using tabletop centrifuges to get rid of particulates. Humidified chambers were prepared from empty pipet tip boxes with water placed in the lower chamber and parafilm partially covering the surface of the tip rack. Drops of solutions were placed on the parafilm, and cells were incubated with various solutions by inverting coverslips on top of the droplets. To block non-specific antibody binding, cells were first incubated with a blocking solution (10% donkey serum in PBST) for 30 min at RT. Cover slips were then gently lifted, drained, and incubated with anti-GFP antibody (Aves lab, cat# GFP-1010) diluted 1:100 in the blocking solution overnight at 4°C. Coverslips were then placed into 6-well plates and washed 3 x 5 min in PBST with agitation. Cells were then incubated with Alexa-647-conjugated anti-chicken F(ab)_2_ (Jackson ImmunoResearch 703-606-155) diluted 1:300 in the blocking solution for 4 hours at RT, and then washed 3 x 5 min in PBST with agitation. Stained coverslips were stored at 4°C until imaging.

#### Preparation of cover slips for SMLM imaging

Coverslips with plated cells were rinsed with GLOX/BME buffer and mounted onto single cavity glass slides (VWR, cat #10118-600), pre-filled with 80 μl GLOX/BME buffer immediately prior to the placement of coverslip. Excess buffer was fully adsorbed from the sides and the top of the coverslip with Whatman paper, taking care to keep the coverslip centered on top of the cavity. Coverslips were sealed onto the cavity slide by pipetting Acid-Free Elmer’s No-Wrinkle Rubber Cement (Amazon.com, https://www.amazon.com/Elmers-No-Wrinkle-Rubber-Cement-Acid-Free/dp/B014JUDMBA/ref=sr_1_2?keywords=Elmer%27s+No-Wrinkle+Rubber+Cement%2C+Acid-Free&qid=1637633240&sr=8-2) around the edge, and allowing the rubber cement to cure for ∼15 min. After imaging, rubber cement was gently peeled off, and the slides were soaked in PBST for 5 min at RT to remove cover slips which were then stored in 6-well plates filed with PBST containing 0.02% NaN_3_ at 4°C until further use. In this set up, the GLOX/BME buffer remained at pH 8 for at least 12 hours.

#### SMLM imaging protocol I: Quantification of the effective labeling efficiency and the number of mEGFP molecules in Fn1-mEGFP nanodomains

For the following experiments, 2D imaging was used to maximize the resolution in the plane of imaging. To minimize fluorophore bleaching and maximize the number of collected photons we followed the protocol protocol developed by (Diekmann et al., 2020). Stained Nup96-mEGFP and Fn1-mEGFP cells were imaged in pairs using the same preparation of the GLOX/BME buffer and the same imaging conditions (described below). For each independent experiment (n=4) and for each round of measurements, images of Nup96-mEGFP and Fn1-mEGFP cells taken on the same day were analyzed. SMLM was performed using Nikon A1-HD25 Ti2E microscope equipped with motorized TIRF illumination, 125mW 640 nm solid-state laser, Perfect Focus, and CFI Apochromat TIRF 100xC Oil objective with numerical aperture 1.49 (cat # MRD01905). All images were acquired at the critical angle of incidence (57°) and recorded using a 512 x 512 EMCCD camera (Princeton Instruments), using 128 x 128 central region on the camera. Gain was set to 3, multiplication gain amplifier was set to 20 MHz. Prior to acquisition, the centered ROI (128 x 128) was bleached at 57° angle, and 10% laser power for 500 frames followed by additional 500 frames at 20% laser power, with the exposure of 60 ms per frame. Images were then acquired at 57° angle, 20% laser power, for 60,000 frames at 60 ms per frame. Images were processed, rendered, and quantified using SMAP and Matlab version 2020a (Ries, 2020), as described (Diekmann et al., 2020) and the documentation found on https://github.com/jries/SMAP. Settings for peak finding in the SMAP software were set according to our camera’s manufacture’s specifications and were the following: EM was set to “on”, camera pixel size was set to 0.162 μm, EM gain was set to 300, e-/ADU conversion factor was set to 2.35. Data was fitted using a Gaussian PSF model, the cutoff parameter for peak finding was set to 2, and ROI size was set to 7 pixels. Localizations were then grouped and rendered, using default parameters in SMAP. Drift correction was performed on rendered localizations in timepoint blocks of 10 or 20, as recommended (Thevathasan et al., 2019) and https://github.com/jries/SMAP. Following satisfactory drift correction judged by the overlapping cross-correlations, localizations were filtered according to the recommended settings for Alexa-647 fluorophore (Thevathasan et al., 2019); In brief, localizations with poor precision were filtered out by limiting localization precision to 0 – 15 nm, out-of-focus localizations were excluded by setting the PSF range to 0 – 150 nm in the x-y plane, poorly fitted localizations were further filtered out by setting the LLrel parameter (relative log-likelihood) in SMAP to a negative cut-off value leaving the majority of the peak intact, and the first 1000 frames were excluded from the analyses. All the remaining grouped localization were rendered according to 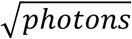 and images were color-coded using LUTs set according to localization density.

Image resolution was measured using the Fourier Ring Correlation (FRC) method described by (Nieuwenhuizen et al., 2013) and implemented in SMAP. Voronoi cluster discovery was performed according to (Andronov et al., 2016a) and their algorithm implemented in SMAP. DBSCAN cluster analysis was performed according to (Caetano et al., 2015; Ester et al., 1996) implemented in SMAP; For DBSCAN, the minimum number of points in the neighborhood (k) was set to 4, as recommended for all 2-dimentional data (Ester et al., 1996). The neighborhood radius *ε* was set either to 14 nm (the average apparent radius of Fn1 nanodomains determined as described below) or automatically estimated by the DBSCAN algorithm in SMAP (Ries, 2020).

NPC radius, effective labeling efficiency and the number of gourped localizations per fluorophore were determined using the algorithm in SMAP following the published automated workflow to segment and analyze nucleopore complexes (NPCs), using the parameters recommended by (Thevathasan et al., 2019) without modifications. In brief, NPCs in focus (mean fitted PFS size of each NPC was less then 145 nm) were automatically segmented using circular ROI of 220 nm in diameter. The number of grouped localizations per NPC, NPC radius, and effective labeling efficiency (ELE) were determined using published algorithms implemented in SMAP without modifications, as described in (Diekmann et al., 2020; Thevathasan et al., 2019) and the SMAP user manual “Analysis of NPCs using SMAP”. Since we used identical staining, imaging, and processing parameters for Nup96-mEGFP and Fn1-mEGFP cells, the number of Fn1-mEGFP molecules per nanodomain was calibrated to the number of grouped localizations per NUP96-mEGFP protein, as described (Thevathasan et al., 2019). In brief, we first determined the number of merged localizations per NPC in the ROI (L_ref_). Each NPC contains 32 NUP96-mEGFP proteins, therefore, the number of merged localizations per Nup96-mEGFP (N_ref_) is N_ref_= L_ref_/32. To determine the number of Fn1-mEGFP proteins per Fn1 nanodomain, we segmented Fn1 nanodomains manually using a circular ROI of 60 nm in diameter and the number of merged localizations per ROI (L_t_) was determined by using the *countingStatistics plugin* in SMAP, as outlined in the SMAP manual “Analysis of NPCs using SMAP”. The number of Fn1-mEGFP proteins per nanodomains (N_t_) is L_t_ / N_ref_. Since Fn1 is only secreted as a dimer, the number of Fn1 dimers per nanodomain is N_t_ /2.

Nanodomain periodicity in Fn1 fibrils stained by a variety of different antibodies was assayed and quantified according to (Fruh et al., 2015). In brief, MEFs were stained using a variety of antibodies at different dilutions, or using combinations of antibodies, as noted in **Figs. 5, 6, 8**, according to the staining protocol described above and imaged using GLOX/BME buffer and the exact imaging settings as described for imaging Nup96-mEGFP and Fn1-mEGFP cells stained with anti-GFP antibodies (see above). Images were processed and rendered in SMAP according to the parameters described above. Gaussian rendering of the localizations (min *σ*Gaussian was set to 3nm) were saved as uncompressed tiff files and opened in Fiji version 2.1.0/1.53c, intensity line profiles along each fibril were generated and imported into Matlab 2021a. To assay periodicity, we used the autocorrelation function implemented in the Matlab’s 2021a Econometrics toolbox and the criteria outlined in (Fruh et al., 2015). In brief, the presence of at least four regularly-spaced peaks in the autocorrelation profile was considered to reflect the periodical nature of Fn1 nanodomains, and the position of the first autocorrelation maximum was taken as a quantitative measure of nanodomain periodicity, as extensively discussed and computationally modelled in the Supplemental Data section of (Fruh et al., 2015). Ljung-Box Q-test for residual autocorrelation was performed using Matlab’s 2021a Econometrics toolbox to assess statistical significance of autocorrelation.

To determine the apparent diameter of Fn1 nanodomains, we imported fibril intensity line profiles obtained from SMLM images of Fn1^mEGFP/mEGFP^ MEFs stained for GFP into Matlab 2021a. To automate the analyses, we used Signal Processing toolbox in Matlab 2021a to fit each intensity peak in the line profile with a Gaussian curve and calculate full width at half height (FWHT) for each peak. Altogether, 1292 nanodomains in 27 long (> 1 μm in length) fibrils from six cells and 3 independent experiments were assessed. We have also performed this analysis manually on a subset of Fn1 nanodomains (n=248) in long fibrils by fitting a Gaussian to an intensity profile of each individual nanodomain and calculating FWHT. The results were the same.

#### Assembly of exogenously-added Fn1: Part I, Live Imaging

3-well culture insert (Ibidi, cat # 80366) was placed in the middle of the 35 mm glass-bottom dish (Ibidi, cat # 81158) and then 0.8 x 10^6^ Fn1-tdTomato-expressing MEFs were plated surrounding the inserts and cultured for 24 hours to reach confluency and establish Fn1-tdTomato matrix. At a 24-hour time point, Fn1-mEGFP-expressing MEFs were plated inside the inserts on glass without coating for 5 hours. Prior to imaging, the culture medium was removed and replaced with the imaging medium. Live imaging was performed using Plan Fluor 40x Oil (numerical aperture 1.3). Positions containing Fn1-mEGFP-expressing MEFs were imaged at ∼17-18 min intervals for ∼16 hours in humidified Tokai Hit stage-top incubator at 5% CO_2_. Each position was imaged 40-43 confocal slices at 0.5 μm thickness, the pinhole was set to 1 Airy unit.

#### Assembly of exogenously-added Fn1: Part II, SMLM

The medium containing secreted Fn1-tdTomato fusion proteins was collected after a 72-hour culture of confluent Fn1-tdTomato MEFs generated by CRISPR/Cas9 mutagenesis, as described above. The medium was spun at 300 x *g* for 5 min to pellet debris. The concentration of Fn1-tdTomato in the supernatant was quantified using ELISA kit using antibodies specific to mouse Fn1, which does not cross react with bovine Fn1 (Abcam, cat # ab21097). Fn1-tdTomato-containing supernatant was diluted 4-fold with fresh MEF culture medium to the final concentration of 5 μg/ml of Fn1-tdTomato, and 2 ml of this supernatant were added to Fn1-mEGFP MEFs plated on 25 mm 1.5H glass coverslips the day before at 10^5^ cells per well in a 6-well plate. Following 24-hours of incubation at 37°C and 5% CO_2_, cells were washed with PBS and fixed in 4% PFA in PBS for 20 min at RT. PFA was quenched with 100mM NH_4_Cl in PBS, cell were washed 3×5 min in PBS, and permeabilized with PBS containing 0.1% Triton-X (PBST). Cells were then incubated with a blocking solution containing 10% donkey serum in PBST for 30 min at RT. To detect Fn1-tdTomato, cells were incubated at 4°C overnight with rabbit anti-mCherry antibody (Abcam, cat #ab167453) diluted 1:100 in the blocking solution. Primary antibodies were detected with Alexa-647-conjugated donkey anti-rabbit F(ab)_2_ secondary antibodies (Jackson laboratories, cat # 711-606-152) diluted 1:300 in blocking solution and incubated for 4 hrs at RT. Cells were then washed 3×5min in PBST and stored at 4°C until imaging. dSTORM imaging was performed using GLOX/BME buffer. Cell were imaged at 57° angle using the SMLM imaging protocol, as described above. Images were processed and reconstructed using SMAP using the same parameters, as described above.

#### Live imaging using TIRF

Fn1^mEGFP/+^ MEFs were plated on 35-mm round glass bottom Mattek dishes (catalog # P35G-1.5-14-C), complete medium was switched to imaging medium prior to filming. Live TIRF microscopy was performed using Nikon A1-HD25 inverted confocal microscope equipped with 4 laser lines of 100mW per line at 405, 488, and 561nm and 125mW at 640nm, and motorized TIRF illumination. CFI Apochromat TIRF 100xC Oil objective and EMCCD camera were used. Before imaging lasers were aligned and the critical angle of incidence for imaging was determined by the software. The exposure time was 20 ms and readout speed was set at 10 MHz.

#### Live imaging using confocal point-scanning microscopy

Live cell imaging was performed using Ibidi glass bottom 8-well chambers (catalog # 80827). 0.6*10^4^ wild-type or Fn1^mEGFP/+^ MEFs were plated on glass in each well of Ibidi glass bottom 8-well chambers and allowed to grow overnight prior to staining and imaging by direct Stochastic Optical Reconstruction Microscopy (dSTORM), see SMLM imaging protocol II below. For FUD and III-11C treatment, Fn1^mEGFP/+^ MEFs were plated in 8-well glass Ibidi dishes (1 cm^2^ growth area) without coating at a density of 0.6×10^4^ cells/well in complete medium. After 5 hours, DMEM was removed and cells were rinsed once with 1X PBS. Subsequently, the medium was changed to imaging medium. For FUD experiments, imaging medium was supplemented either with 225 nM FUD or 274 nM of control III-11C peptide. Untreated wells contained cells incubated with imaging medium. Following the addition of the imaging medium (with or without the peptides), the chamber was immediately set up for imaging in the humidified Tokai Hit stage-top incubator at 37°C, 5% CO_2_. Live imaging was performed using Nikon A1-HD25 inverted confocal microscope with the DUG 4-Channel Detector and 2 GaAsP, 2 high-sensitivity PMTs, and a motorized XYZ stage with Nikon’s Perfect Focus 4 system, and Plan Fluor 40x Oil (numerical aperture 1.3, cat # MRH01401). mEGFP was excited using 488 nm laser at 1% power and pinhole set to 1 Airy unit. An optical zoom of 2 and Z step size of 0.5 μm were used, and stack size was set to 10-15 microns allowing to image the entire cell. For overnight movies, each position was filmed every 1.5 min – 4 min, as noted in Movie legends. Movies in the mp4 format were generated using Imaris 9.5.1 (Bitplane), titles and arrows were added using Adobe Premiere Elements Editor 2020.

#### SMLM imaging protocol II: Imaging cells plated on Ibidi glass-bottom dishes

This protocol was used to image cells following the live DOC assay or overnight live imaging (**Fig. 10** and **Sup. Fig. 9**). Following fixation with 4% PFA, cells were washed with PBS, incubated with blocking buffer containing PBS, 0.1% Triton-X, and 10% donkey serum for 30 min at RT. Cells were then incubated with Abcam monoclonal anti-Fn1 antibody diluted at 1:300 in blocking buffer overnight at 4°C. Following three washes at 5 min each in PBST (PBS with 0.1% TritonX), cells were incubated with Alexa-647-conjugated anti-rabbit secondary antibodies diluted 1:300 in blocking buffer for 4 hours at RT, and washed three washes at 5 min each in PBST. Prior to imaging, freshly prepared GLOX/MEA buffer was added and the chamber was immediately sealed using parafilm. STORM was performed using Nikon A1-HD25 Ti2E microscope equipped with motorized TIRF illumination, 125mW 640 nm solid-state laser, Perfect Focus, and a 100x/1.49NA objective. Images were acquired at the critical angle of incidence and recorded using a 512 x 512 EMCCD camera (Princeton Instruments). Calibration was obtained by imaging of 100 nm Tetraspeck beads (Life technologies, catalog # T-7279) using the same glass surface and buffer conditions. To drive Alexa-647 into the dark state, samples were pre-bleached by the illumination at 640 nm for 10 seconds at 100 % laser power. 40,000 frames were acquired at 8.4 ms exposure. Blinking events were fitted using the Nikon N-STORM localization software. Images were analyzed using Nikon software (Nikon Elements AR Software v5.11.01). Localization events with fewer than 800 or more than 50000 photons were filtered out to remove blinking evens that were either too faint or too bright (Jimenez et al., 2020). In addition, blinking events were filtered out if they occurred in more than 5 consecutive frames or where outside the z-range determined by the calibration using 100 nm Tetraspeck beads. Images in which z-rejection was below 50% were used for further analyses.

#### Analysis of localization numbers in fibrillar and non-fibrillar nanodomains

In order to enrich for non-fibrillar nanodomains, Fn1^mEGFP/+^ MEFs were plated in 8 well glass Ibidi dishes (1 cm^2^ growth area) without coating. Cell were plated at the density of 0.6×10^4^ cells/well in imaging medium with or without FUD (225 nM) or III-11C (274 nM), and incubated in at 37°C, 5% CO_2_ for 1 hr. Subsequently, MEFs were rinsed once in warm 1X PBS and fixed using pre-warmed 4% PFA for 20 min. After fixation, wells were rinsed three times, 5 min each with Stain Buffer (cat # 554656 BD Pharmingen), blocked for 30 min at room using 5% Donkey serum prepared in Stain Buffer, and incubated with the monoclonal anti-Fn1 (Abcam, cat # 199056) overnight at 4° C. Cell were then washed with Stain Buffer three times, 10 min each, and incubated with anti-rabbit antibodies conjugated with Alexa-647 for 1 hour at rt. Cell were then rinsed again with Stain Buffer three times, 10 min each, and stored at 4° C in 1X PBS. STORM imaging was performed, as described in SMLM imaging protocol II. To quantify the number of grouped localizations per nanodomain, we used the free-hand ROI tool in the STORM window (Nikon Elements AR Software v5.11.01) to determine the number of localizations within non-fibrillar and fibrillar Fn1 nanodomains. Fn1 nanodomains were analyzed in 5 random regions from 3 independently acquired images (a total of 15 fields) for each sample. To determine the number of localizations in Fn1 nanodomains within fibrils, we analyzed more than 20 from 3 or more independently acquired images. All the counts were plotted in Prism 8.2.1 (GraphPad Software, USA), and compared using either one-way ANOVA test with Tukey’s correction or Kruskal-Wallis test with Dunn’s correction for multiple testing.

#### Double-color dSTORM acquisition

Samples were mounted with the SMART Kit buffer (Abbelight). 2D or 3D STORM images were acquired using a SAFe360 module (Abbelight) coupled to an inverted bright-field Olympus IX83 microscope, equipped with a 100X oil-immersion objective (1.49 NA). This dual-cam system (sCMOS cameras, Orcaflashv4, Hamamatsu) allows to perform multicolor STORM by spectral demixing strategy coupled to the use of far-red dyes (BioOptics World, 2021, Caorsi and Karlsson).

Briefly, a dichroic at 700nm is used to separate the fluorescence emission from AF647 and CF680 dyes. The Point Spread Function (PSF) of each detection can be retrieved on both cameras and the measured photon numbers is related to the spectral separation of the fluorophore (**Fig. M1**).

**Figure M1.**
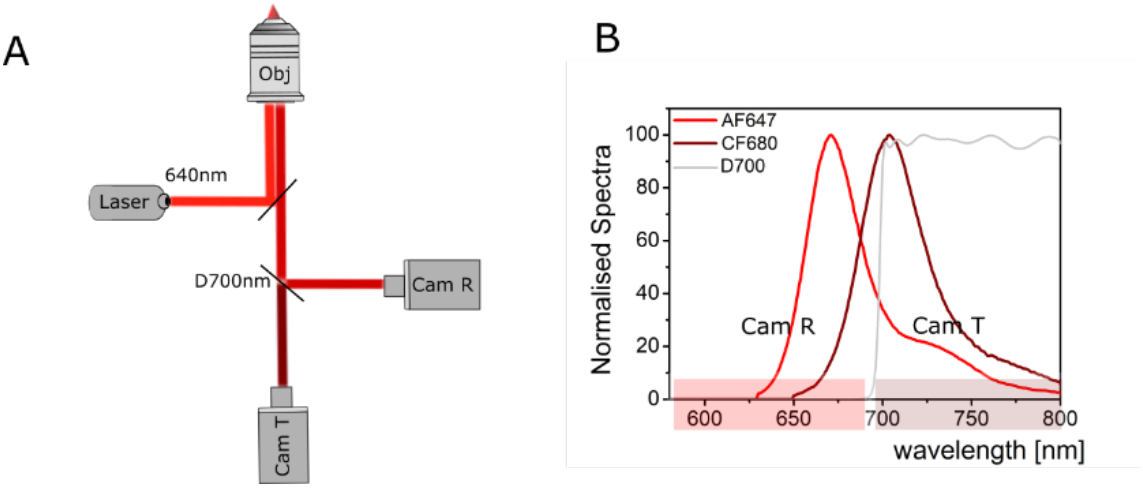
Spectral de-mixing optical scheme. **A.** A 640 nm laser is sent through the objective lens to excite AF647 and CF680 fluorophores. The emitted light is separated by a 700 nm long pass dichroic filter that reflects light below 700 nm into the Reflected camera (Cam R) and transmits light above 700 nm into the Transmitted camera (Cam T). **B.** Emission spectra of AF647 and CF680 dyes together with a 700nm dichroic filter showing spectral separation into the two cameras Cam R and Cam T.

We acquired 60000 frames at 20ms exposure time on a camera sensor size of 1024×1024 pixel to collect single-molecule detections. The irradiation at the sample was tuned according ASTER technology (Mau et al., 2021) implemented on the SAFe360 Abbelight module. Resulting coordinate tables and images were processed and analyzed using NEO software (Abbelight).

As the PSF is captured on both cameras, transmitted and reflected, a ratiometric analysis is applied: a ratio for each detection is calculated and the final ratio distribution is used for lambda assignment:

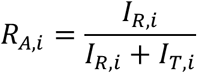

where the suffix A is the fluorophore, i the localisation and I_R_ and I_T_ the intensity measured on camera R and camera T (i.e. the number of photons emitted per molecule).

Average ratio distributions obtained from measurements are shown in **Fig.M2**:

**Figure M2.**
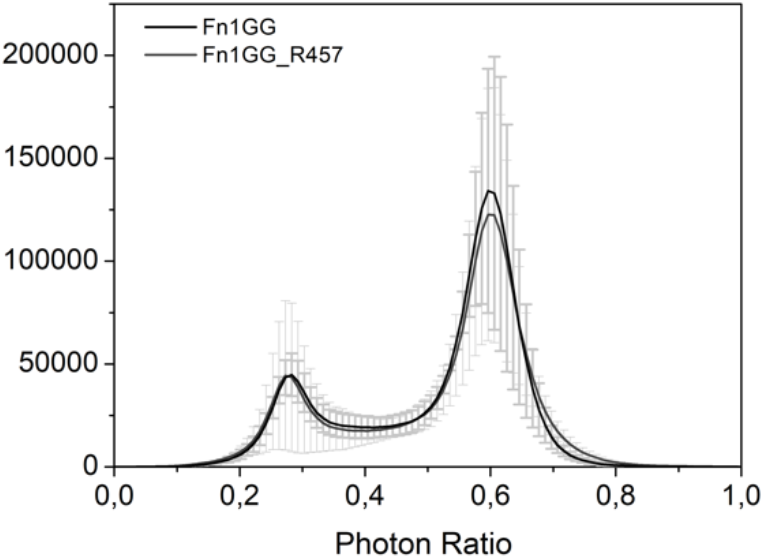
Average ratio distributions for Fn1^mEGFP/mEGFP^ cells stained using chicken anti-GFP and rabbit anti-GFP antibodies (Fn1GG, dark gray) or chicken anti-GFP and rabbit R457 antibodies (Fn1GG_R457, lighter gray). The two fluorophore populations can be clearly distinguished for lambda assignment.

Following the ratio distributions measured on the samples the following parameters have been used for separation: detections with ratios between 0-0.45 were assigned to CF680 and detections with ratios between 0.5-1 were assigned to AF647. On average 3% of detections were rejected while keeping crosstalk below 1%. Following de-mixing, colocalization analysis was performed using Neo software following the CBC algorithm (Malkusch et al., 2012). Parameters set for CBC analysis were Rmax at 300 nm and the number of steps equal to 10. In addition the CBC algorithm as implemented in ThunderSTORM, with R_max_ = 50 nm, and the number of steps equal to 10 (Ovesny et al., 2014) was used to perform CBC analyses (Malkusch et al., 2012).

## Legends for Movies

**Movie 1. Rotational views through the Fn1+ ECM in the cardiac jelly.** The whole E9.5 embryo was stained using Abcam rabbit monoclonal anti-Fn1 antibody and imaged using 100x objective, N.A. 1.49, with the pinhole set at 0.8 Airy units, and sampling of 40 nm per pixel in x, y. The movie shows 3D reconstruction through 3.4 μm of tissue sampled every 0.121 μm in z. Fn1 is in white, DAPI is in blue. Arrows point to examples of beaded Fn1 fibrils.

**Movie 2. Fn1 fibrillogenesis imaged by TIRF microscopy.** Fn1^mEGFP^ MEFs were transiently transfected with mCardinal-Lifeact, plated on gelatin-coated glass coverslips, and imaged 48 hours later. Filming was done every 2 min for 30 min using TIRF and 100x objective, N.A. 1.49. The first set shows the Fn1-mEGFP channel. Yellow arrows point to centripetally-moving Fn1 nanodomains organized into an elongating linear fibril. The second set is an overlay between Fn1-mEGFP and mCardinal-Lifeact.

**Movie 3. 2% DOC dissolved cytoplasm and nucleus in under 13 min leaving cell-free Fn1 fibrils.** MEFs expressing Fn1-mNeonGreen were plated on glass-bottom slides without coating and labeled with SiRActin (magenta) to visualize F-actin, and Hoechst (blue) to visualize DNA. Time-lapse was recorded every 54 sec immediately following the addition of the DOC solution pH 8.01 to live cells. The presence of 2% DOC dissolves actin cytoskeleton and nuclei and leaves cell-free Fn1 ECM fibrils (green). Fn1 fibrils collapse following the dissolution of the actin cytoskeleton due to the loss of tension.

**Movie 4. Fibrillogenesis of ectopic Fn1.** 3-well culture insert was placed in the middle of the 35 mm glass-bottom dish and 0.8 x 10^6^ Fn1-tdTomato-expressing MEFs were plated surrounding the inserts. 24 hours later, Fn1-mEGFP-expressing MEFs were plated inside the inserts on glass without coating for 5 hours. Confocal live imaging of areas containing Fn1-mEGFP-expressing MEFs was performed using Plan Fluor 40x Oil (NA 1.3). Positions containing Fn1-mEGFP-expressing MEFs were imaged at 17-18 min intervals for ∼16 hours. This movie contains maximum intensity projections composed of forty-three confocal slices at 0.5 μm thickness, the pinhole was set to 1 Airy unit. The first still panel in this movie is a montage of the plate to show Fn1-tdTomato and Fn1-mEGFP-expressing cells prior to the start of the time-lapse.

**Movie 5. Cells incubated with 11-IIIC, show robust fibrillogenesis.** Fn1^mEGFP/+^ MEFs were plated on uncoated glass in 8-well Ibidi chambers for 4 hours. Medium containing 11-IIIC control peptide was then added and cells were filmed every 90 sec for about 15 hours, as described in Methods. The movie begins approximately 30 min after the 11-IIIC-containing medium was added, the time it takes to set up time-lapse recording. Arrows point to the cell periphery and examples of centripetally moving Fn1 fibrils.

**Movie 6. FUD interferes with linking centripetally moving Fn1+ nanodomains into fibrils.** Fn1^mEGFP/+^ MEFs were plated on glass in 8-well Ibidi chambers for 4 hours. Medium containing FUD peptide was then added and cells were filmed every 3 min for about 15 hours, as described in Methods. The movie begins approximately 30 min after the FUD-containing medium was added, the time it takes to set up time-lapse recording. Note the dismantling of pre-existing fibrils at the beginning of the movie. Yellow and red arrows point to the cell periphery. Note the presence of centripetally moving Fn1-mEGFP “beads” and the scarcity of Fn1 fibrils for the majority of the duration of the movie.

---

**Supplemental Figure 1.**
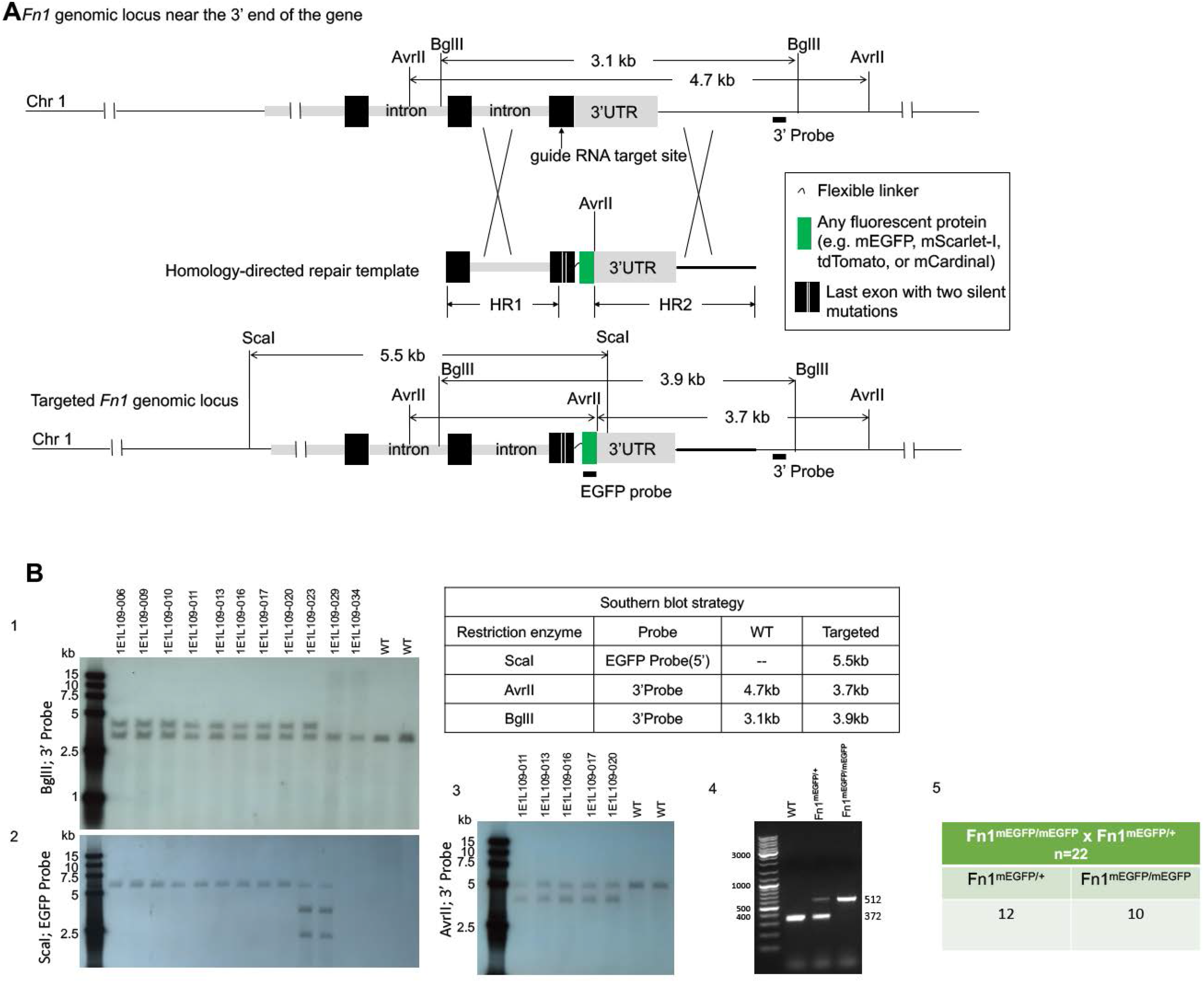
Construction of Fn1^mEGFP^ allele. **A.** Targeting construct. **B1-B3.** Southern blots. **B1.** Southern blot with 3’ probe after digestion with Bgl II. **B2.** Southern blot with GFP probe after digestion with Sea I. **B3.** Southern blot with 3’ probe after digestion with Avrll. **B4.** Diagnostic PCR detecting Fn1^+/+^; Fn1^mEGFP/+^; and Fn1^mEGFP/mEGFP^. **B5.** Fn1^mEGFP/mEGFP^ mice are obtained at a Mendelian ratio in the cross between Fn1^mEGFP/mEGFP^ and Fn1^mEGFP/+^ mice.

**Supplemental Figure 2.**
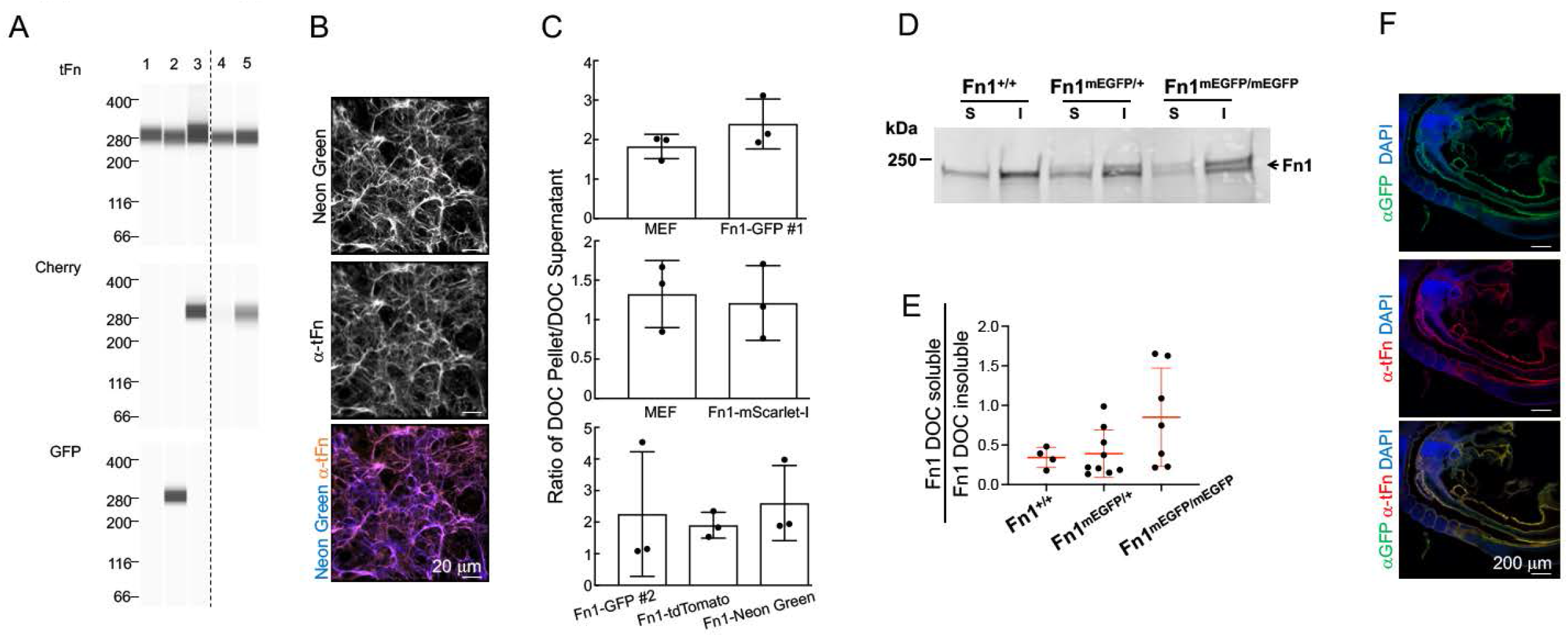
Fn1-FP fusion proteins behave like wild-type Fn1 in biochemical assays and *in vivo.* CRISPR was used to knock-in mEGFP, tdTomato, mNeonGreen, or mScarlet-l at the 3’ of Fn1 replacing the termination codon in MEFs (see Sup. Fig. 1). **A.** Western blot of cell extracts: 1) wild-type MEFs, 2) Fn1^mEGFP^ MEFs, 3) Fn1^,dTomato^ MEFs, 4) wild-type MEFs, 5) Fn1^mScarlet-l^ MEFs. **B.** Fn1^mNeonGreen^ MEFs were stained with antibody to total Fn1; Native mNeonGreen fluorescence of Fn1-mNeonGreen proteins co-localizes with Abeam monoclonal antibody to tFn1. **C.** Fn1-FP fusion proteins behave like wild-type in DOC matrix assembly assays. **D-E.** Fn1-mEGFP proteins extracted from E9.5 embryos behave like wild-type in DOC matrix assembly assay. **D.** E9.5 embryos were solubilized using DOC lysis buffer and DOC-soluble and insoluble fractions were resolved on a 4-12% SDS-PAGE gel followed by a Western blot using Abeam monoclonal anti-Fn1 antibody. E. Quantification of the fraction of Fn1 DOC-soluble Fn1 to DOC-insoluble Fn1. Each dot is an embryo. One-way ANOVA analysis showed no statistical differences between samples from different genotypes. **F.** E9.5 Fn1^mEGFP/+^ embryos were stained with Abeam monoclonal antibody to Fn1 and anti-GFP antibody to detect Fn1 and GFP proteins; Fn1-mEGFP and Fn1 co-localize and are expressed in the expected pattern.

**Supplemental Figure 3.**
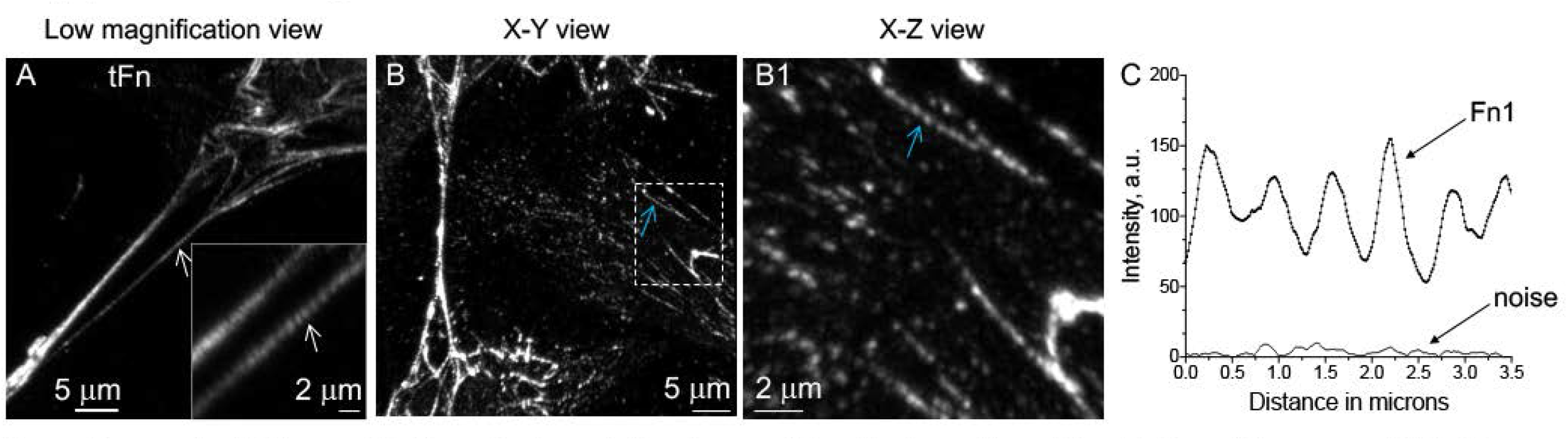
Beaded architecture of Fn1 visualized by Zeiss Airyscan. Wild-type MEFs were plated for 48 hours on gelatin-coated glass, fixed and stained to detect Fn1 using the Abeam monoclonal anti-Fn1 antibody, and imaged with the Airyscan modality on Zeiss confocal microscope. **A.** Arrow points to a Fn1 fibril between cells, magnified in the inset. Note the beaded appearance of this fibril, arrows in **A.** Box in **B** is expanded in **B1** and rotated to show the x-z axis, blue arrow points to a beaded fibrillar adhesion. **C.** Intensity profile of the Fn1 fibril marked by the blue arrows in **B** and **B1.** Panel **C** shows that Fn1 fibrils consist of regions of high and low fluorescence intensity.

**Supplemental Figure 4.**
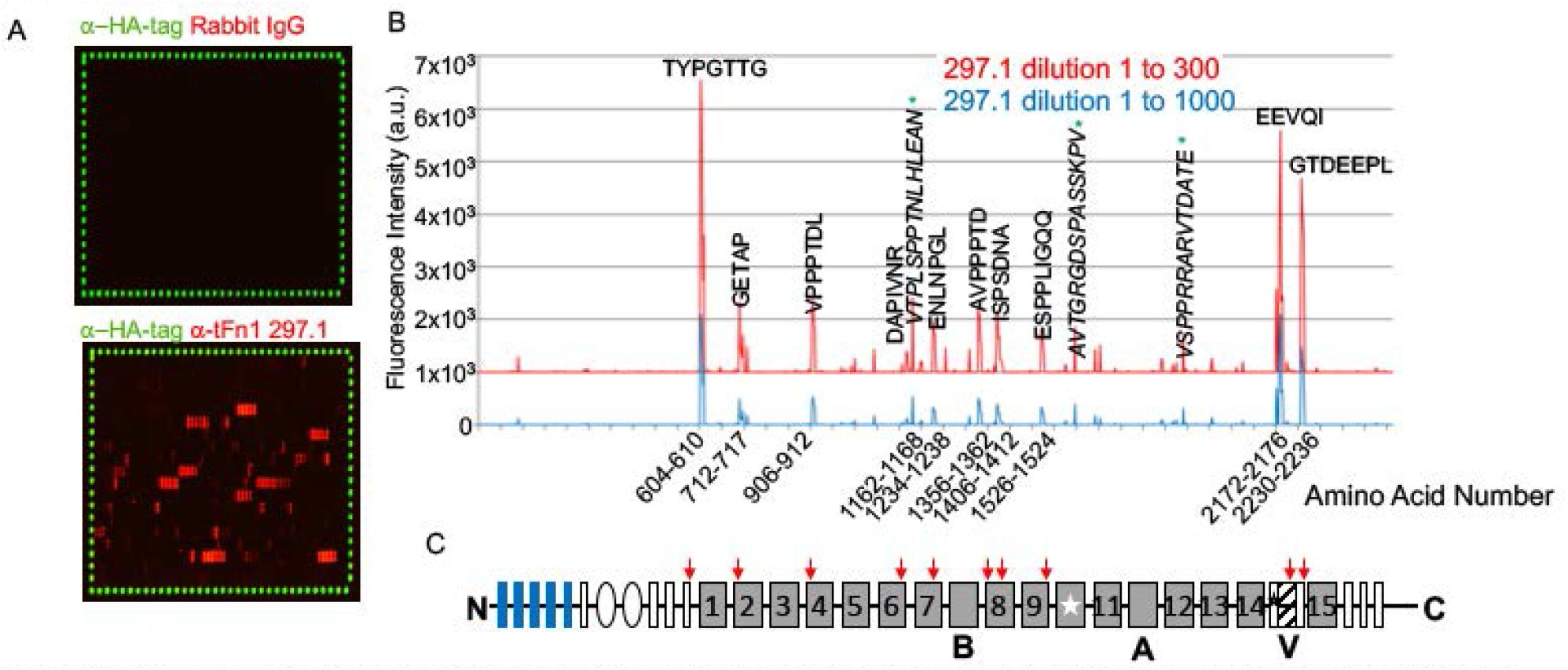
297.1 antibody recognizes multiple epitopes along the Fn1 molecule. **A.** Custom-prepared peptide arrays fabricated to contained 15-amino acid peptides with 13-amino acid peptide-peptide overlap, spanning the entire mouse Fn1 sequence. HA-tag peptides were spotted around the perimeter for control purposes. Arrays were incubated with anti-HA-tag antibodies followed by the appropriate secondary antibody (green) and with either rabbit IgG (top panel) or 297.1 antibodies (bottom panel) and the appropriate secondaries, red. **B.** Multiple strong to very strong antibody responses against epitope-like spot patterns formed by adjacent peptides with the consensus motifs annotated in the intensity plot at high signal-to-noise ratios; We also observed a few atypical interactions with peptide as highlighted in *italics and marked by asterisks.* Amino acid positions of epitopes are marked on the x-axis. **C.** Positions of 297.1 epitope-binding sites are mapped on mouse Fn1 molecule. The splicing of the V region is depicted according to the mouse mRNA tracks in GENCODE VM27.

**Supplemental Figure 5.**
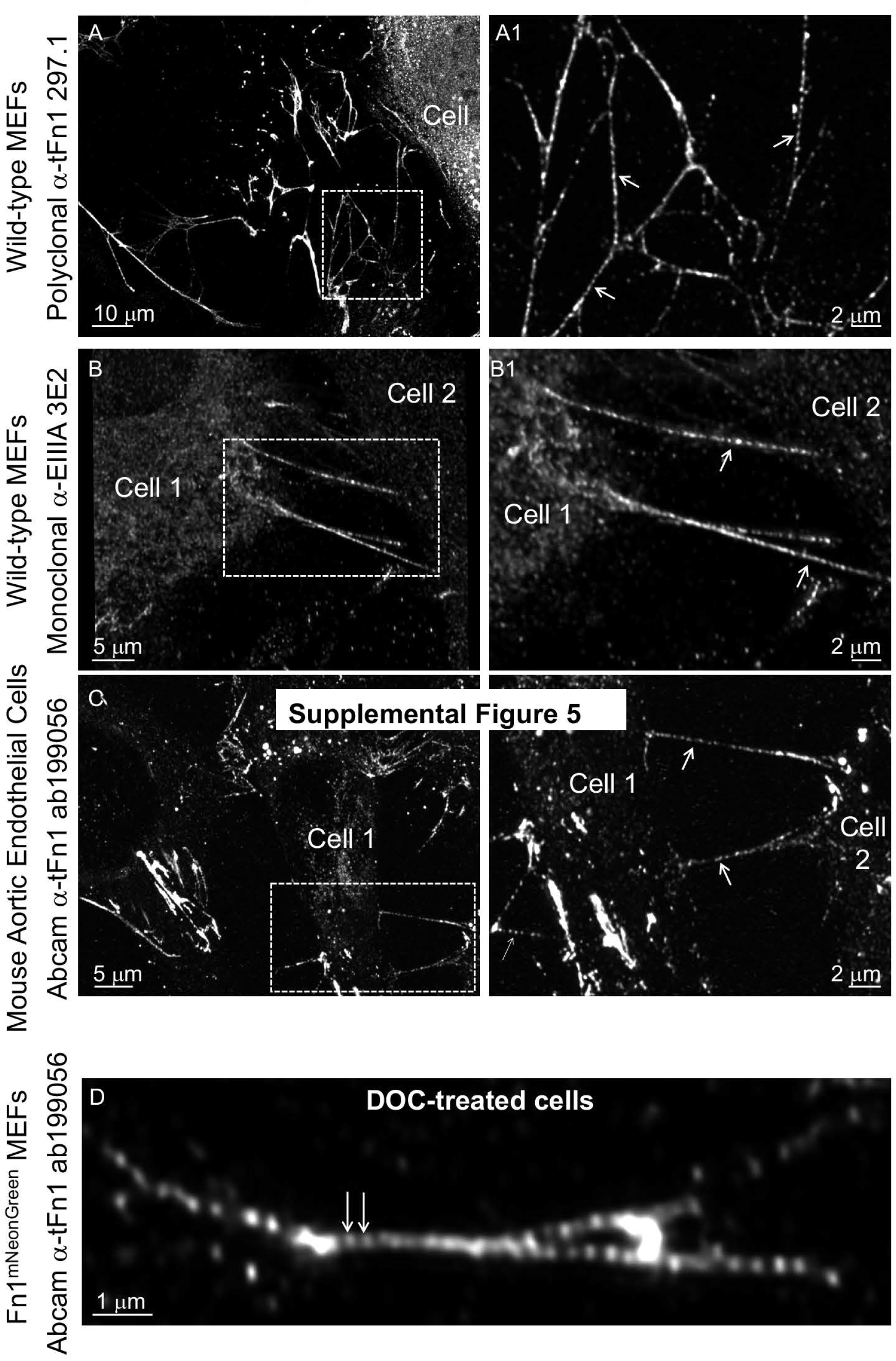
Beaded architecture of Fn1 fibrils is seen with multiple antibodies, in different cell types, and is retained in the absence of cell contact. Cells were plated on glass without coating. **A–B.** Fn1 secreted by wild-type MEFs and deposited **A)** on glass or **B)** located between two cells. **C.** Fn1 fibrils between endothelial cells. Boxes in **A–C** are magnified in **A1-C1.** Arrows point to fibrils deposited in the intercellular space (**A1-C1**). **D.** Fn1 fibril imaged following the treatment with 2% deoxycholate for 10 minutes, and staining with Abeam monoclonal antibody to Fn1. Note the beaded architecture of long Fn1 fibril (arrows). Solubilization of cell components by this treatment is shown in **Movie 3**. All images were collected using 100x oil objective, NA 1.49, pinhole size 0.8 Airy units, and sampling resolution of 40 nm/pixel in x,y.

**Supplemental Figure 6.**
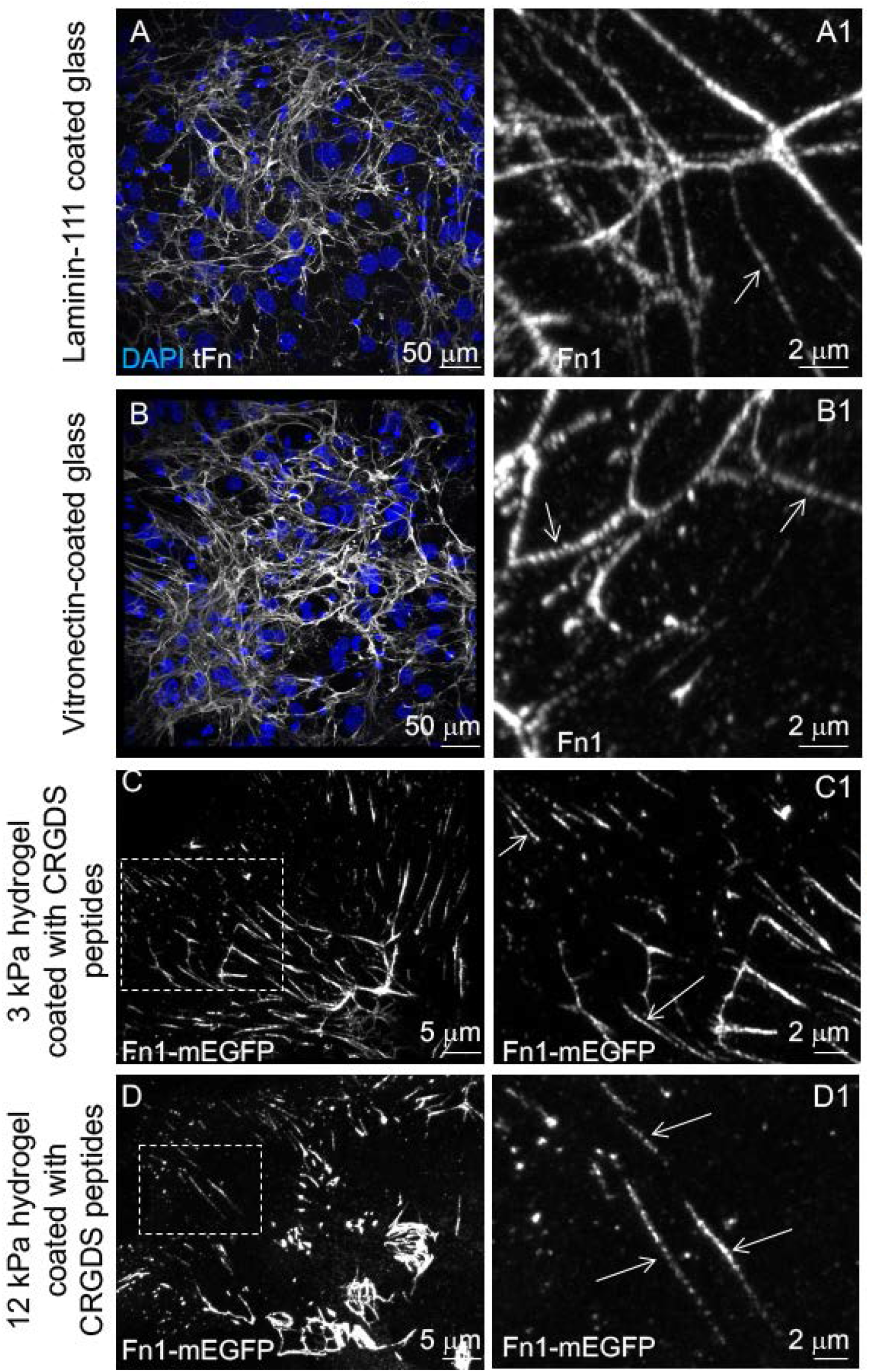
Beaded architecture of Fn1 fibrils is observed when cells are plated on different substrata. **A-A1.** Wild-type MEFs plated on laminin-111 for 48 hrs. **B-B1.** Wild-type MEFs plated on vitronectin for 48 hrs. Cells in **A-A1, B-B1** were stained using Abeam monoclonal anti-Fn1 antibody. **A-B.** Low magnification images were recorded with 40x oil objective, **A1-B1.** High magnification images were captured using 100x oil objective, NA 1.49; pinhole was set to 0.8 Airy units, and sampling was done at 40 nm/pixel in x,y (See Methods). **C-D.** Fn1^mEGFP/+^ MEFs were plated for 16 hours on hydrogels derivatized with GRGDSPC peptides (See Methods). Native mEGFP fluorescence was imaged using 100x oil objective, NA 1.49, pinhole size 0.8, and sampling at 40 nm/pixel in x,y). **C–C1.** 3 kPa hydrogel; **D–D1** 12 kPa hydrogel. Arrows point to examples of beaded fibrils.

**Supplemental Figure 7.**
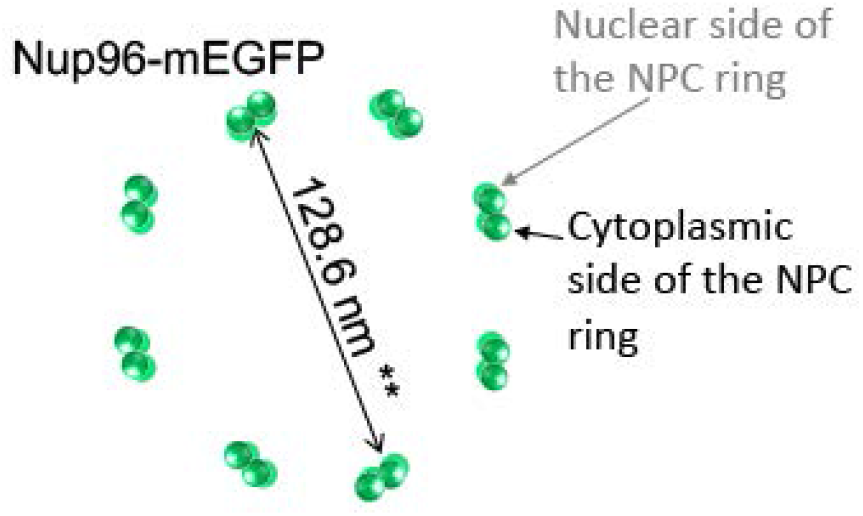
Nucleopore complex schematic and diameter measured using anti-GFP antibodies. NPC contains 32 copies of NUP96 protein arranged with an 8-fold symmetry. 16 copies of NUP96 face the cytoplasm, and 16 face the nucleoplasm. **The radius of NPCs measured using NUP96-mEGFP and anti-GFP primary and Alexa-Fluor647-labeled secondary antibodies is 64.3+/-2.6 nm (Thevasian et al., 2019)

**Sup. Figure 8.**
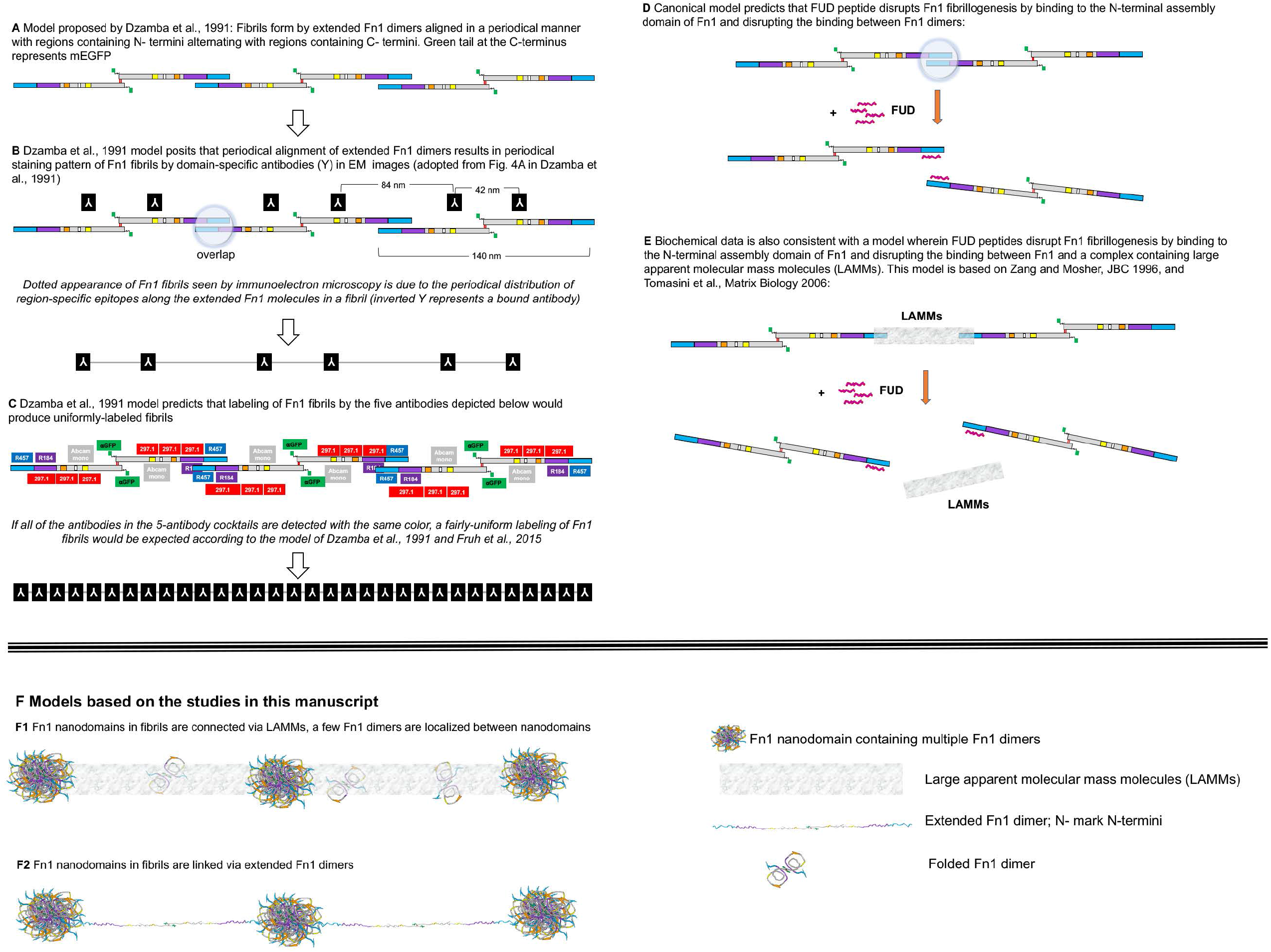

**Supplemental Figure 9.**
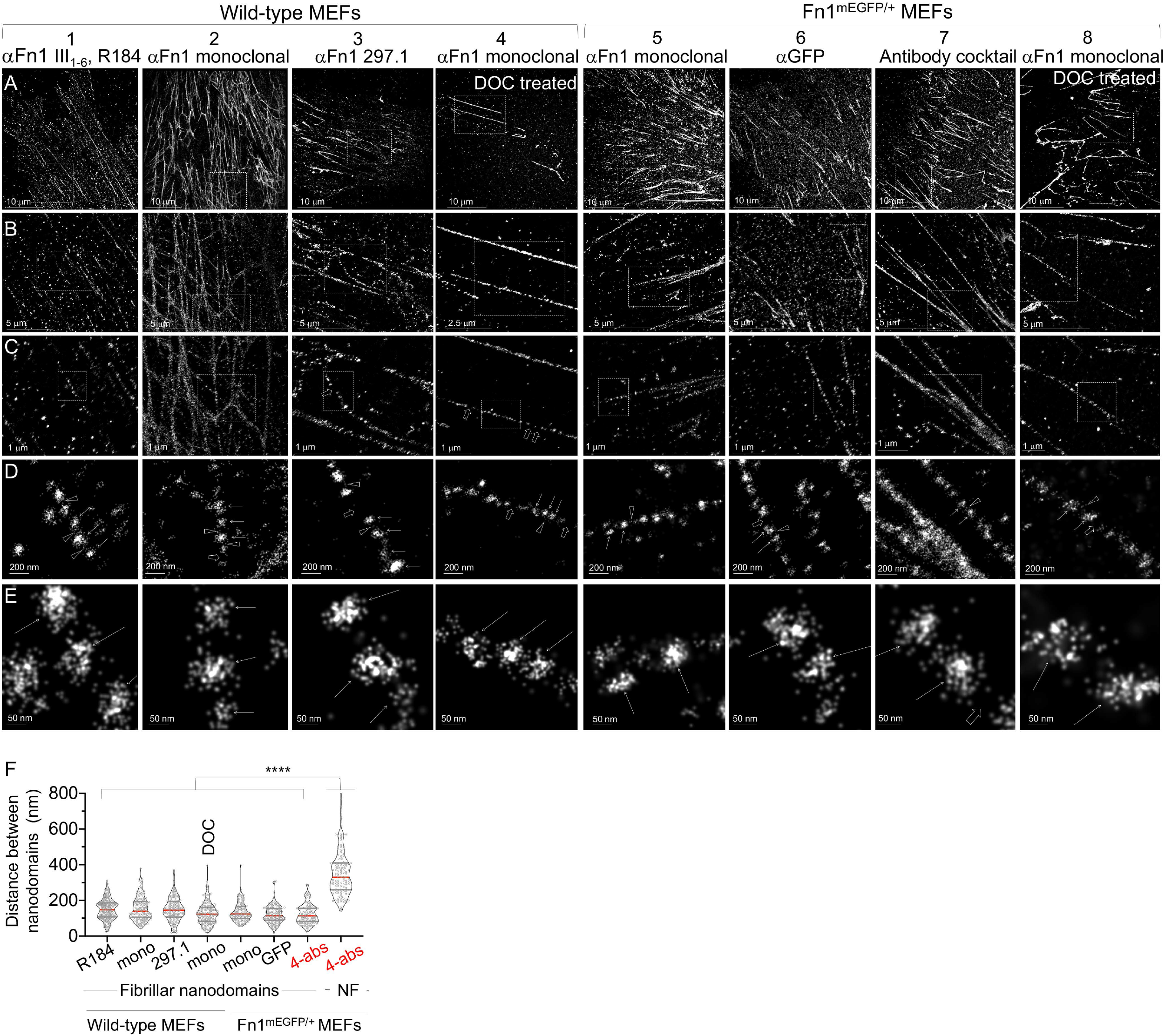
Wild-type or Fn1^mEGFP/+^ MEFs were plated on glass for 16 hrs, fixed and stained with different antibodies to Fn1 followed by Alexa 647-conjugated secondary antibodies. Columns 4 and 8 show cells treated with 2% DOC prior to fixation. Cells were imaged using SMLM imaging protocol II. **A.** zoom-out views to show the overall appearance of Fn1 fibrils. **B-E.** Successive magnifications of fibrils shown in row (**A**). Arrows in **D** point to nanodomains magnified in **E;** arrowheads in **D-E** point to Fn1 localizations between nanodomains, wide open arrows point to Fn1-free zones between nanodomains in a fibril. **F.** distances between nanodomains within fibrils or non-fibrillar (NF) nanodomains, **** p<10^4^, Kruskal-Wallis test, with Dunn’s correction for multiple testing. Please note that nanodomain sizes appear larger in these images than in Fig. 5–8 because the resolution was lower due to the use of high laser power during bleaching and imaging steps, and the 3D acquisition; FRC-measured resolution in these images ranged between 40-50 nm. Antibody cocktail contained: R184 (1:50), Abeam mono (1:300), 297.1 (1:100) and anti-GFP (1:300) antibodies.

**Supplementary Figure 10.**
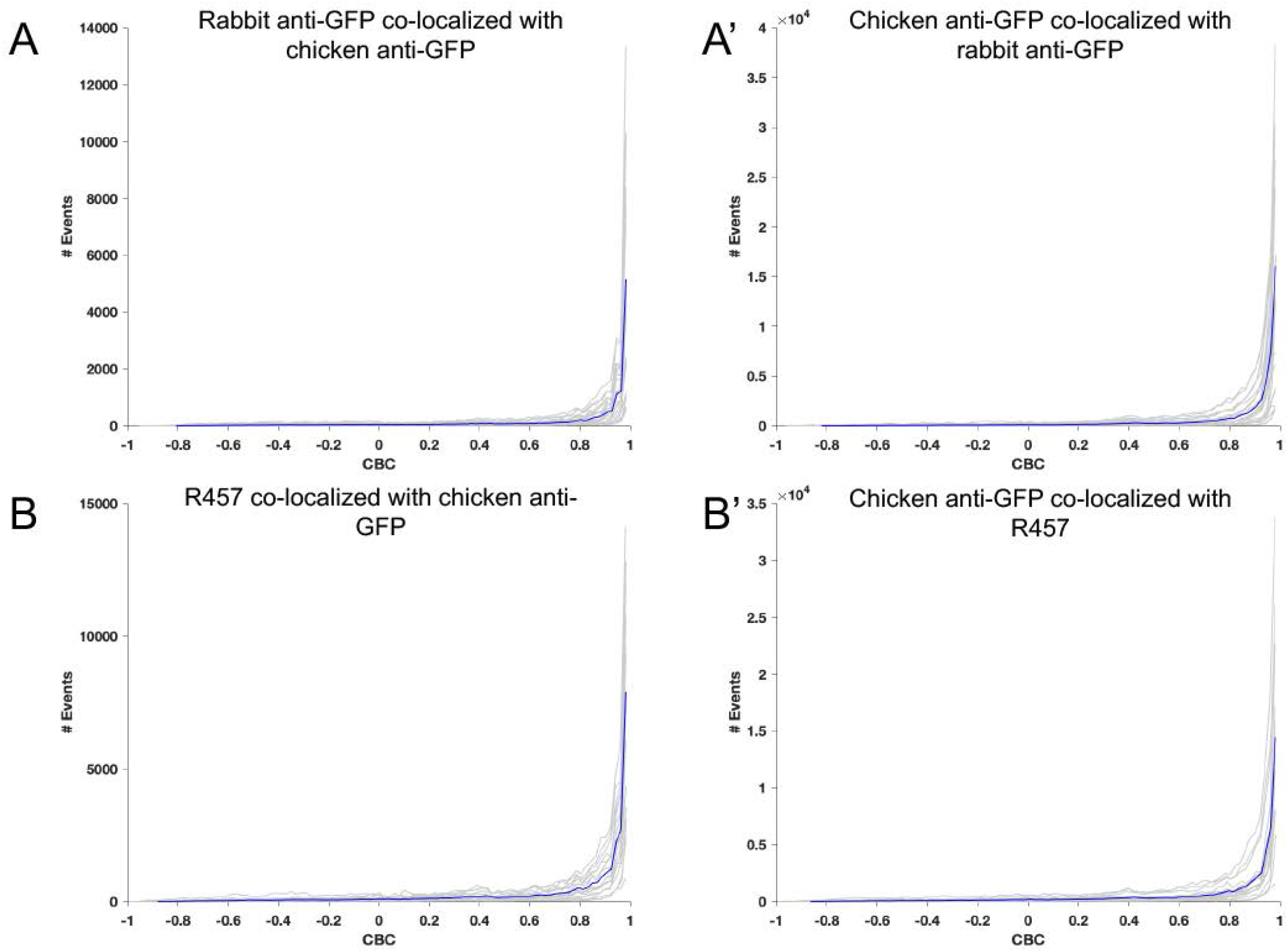
CBC analyses show a near complete colocalization of Fn1 N-terminal and C-terminal domains. Cells, plating and staining were as described in Fig. 9. CBC analyses were done using ThunderSTORM, R_max_ was set to 50 nm, number of steps was set to 10 (r=5 nm). Blue curve is the average of gray curves. For GFPGFP experiments (**A-A’**), 22 images from 4 cells were analyzed; For R457GFP experiments (**B-B’**), 17 images from 4 cells were analyzed. +1 indicates a high probability of co-localization, ≤ 0 absence of co-localization.

**Supplemental Figure 11.**
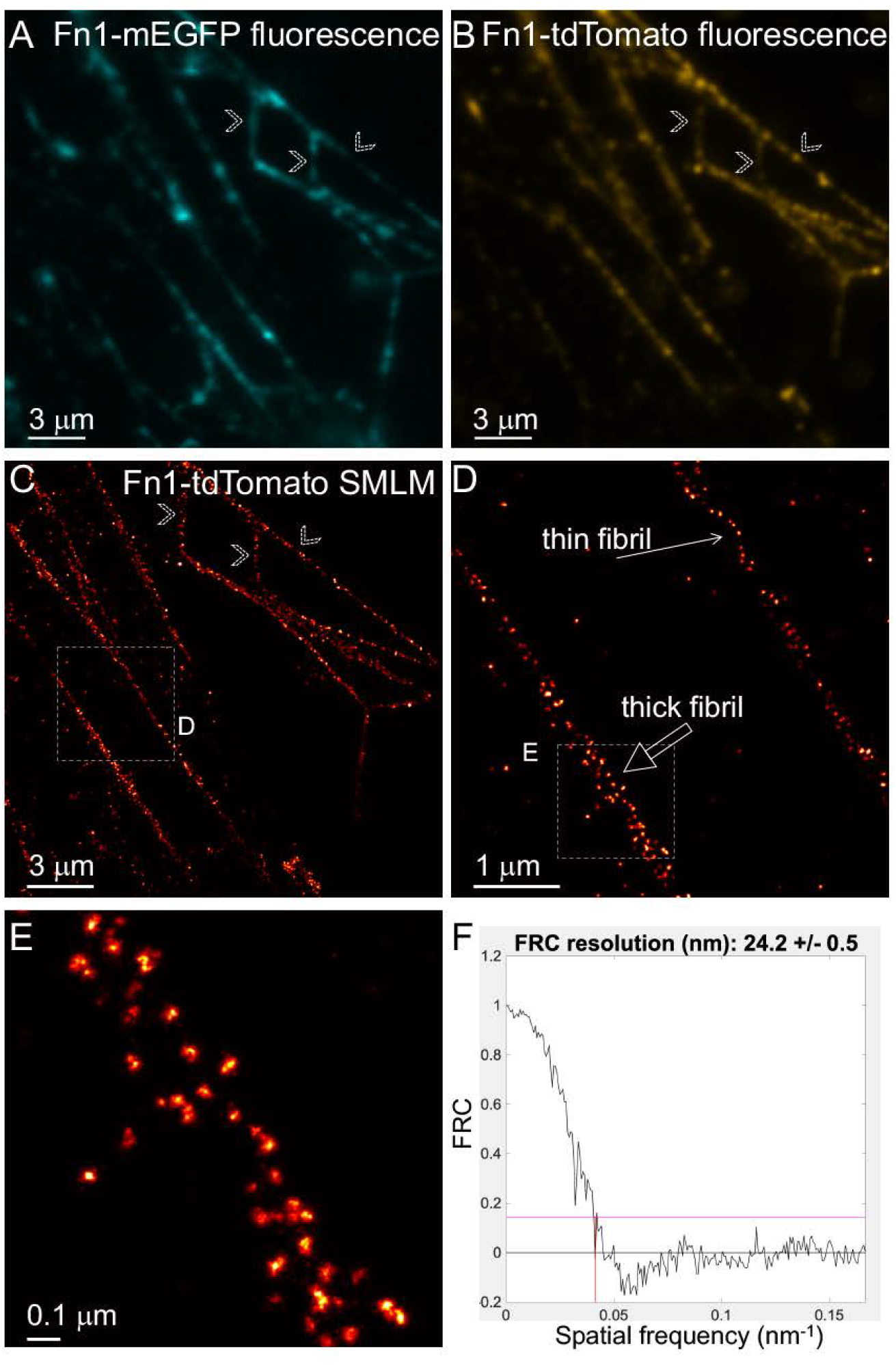
Ectopically-added Fn1-tdTomato co-assembles with endogenous Fn1-mEGFP fibrils and forms nanodomains. Fn1^mEGFP/mEGFP^ were plated on glass without coating at 50% confluency for 16 hours and then Incubated with 10 μg of Fn1-tdTomato for 24 hours. Cells were then fixed and stained with rabbit anti-Cherry antibody, which was detected with secondary antibodies conjugated with Alexa-647. Fn1 ECM deposited between cells was Imaged. **A–B.** Widefield Images taken with 100X Objective, NA 1.49. A. Fn1-mEGFP fluorescence. B. Fn1-tdTomato fluorescence. **C - E.** dSTORM using SMLM protocol I. **C.** dSTORM image of the region shown in **B.** Examples of fibrils in **C** corresponding with fibrils in the wide-field Images (**A-B**) are marked with chevrons. **D.** Magnified region containing a thick and a thin fibril seen in **C. E.** Nanodomain architecture of the thick fibril region boxed in **D.** Note nanodomain architecture of ectopically-added Fn1-tdTomato. **C-E** SMLM images were reconstructed using SMAP (see Methods). **F.** Fourier ring correlation (FRC) analysis performed in SMAP. FRC curve shows the decay of correlation with Increasing spatial frequency. Pink line marks the threshold value of 0.143 calculated for SMLM data (Nieuwenhuizen et al., 2013). Red line marks the spatial frequency for which the threshold falls below the value of 0.143. Resolution is calculated as the inverse of the spatial frequency. Resolution of the region shown in **E** is 24.2+/-0.5 nm.

**Supplemental Figure 12.**
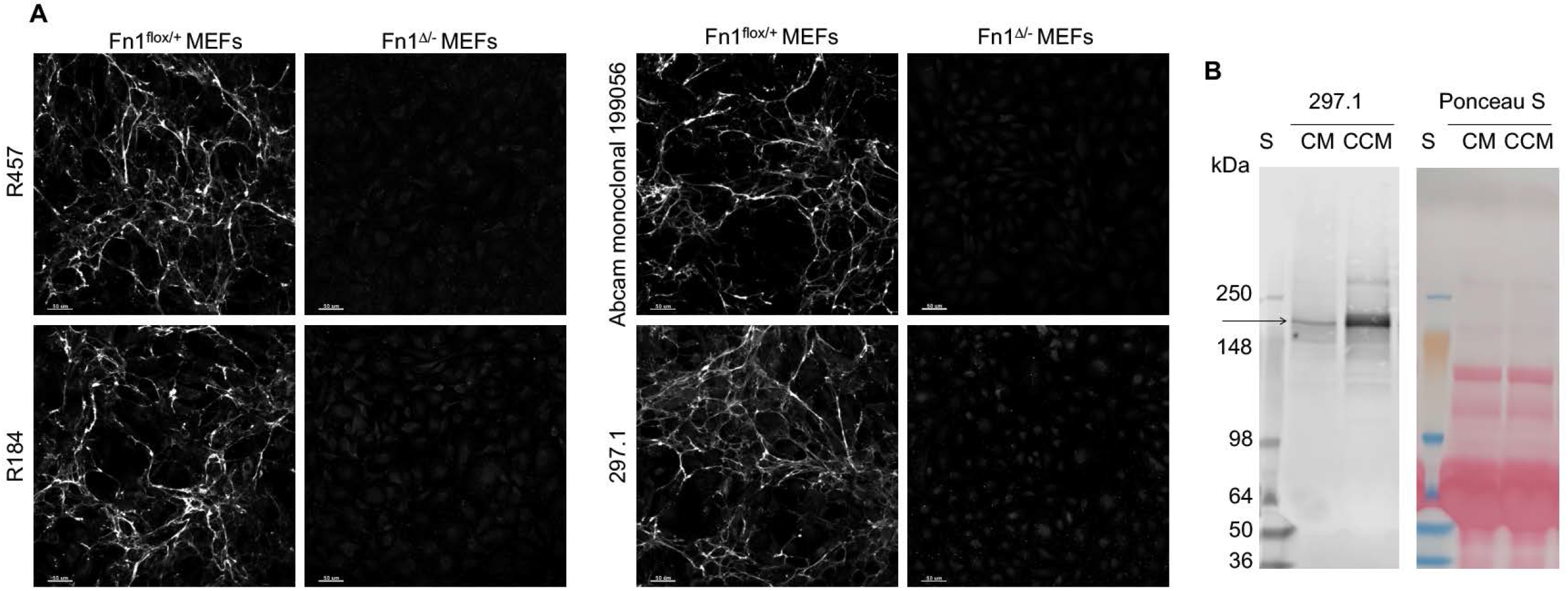
Antibodies utilized in this study are specific to fibronectin. **A.** MEFs were isolated from Fnl^flox/+^ or Fnl^flox/-^ embryos at E14.5. Fn1^flox/-^ MEFs were infected with adenoviruses encoding Cre recombinase, and sorted two days after infection, generating Fnl^Δ/-^ MEFs lacking Fn1. Deletion of Fn1 was confirmed using Western Blotting (data not shown). 3x10^4^ cells per well were plated in a 24-well plate on 1.2 mm glass coverslips without coating, and fixed with 4% PFA 72 hours later. Cells were stained with antibodies indicated to the left of the panels at 1:100 dilution. The same antibody solution was applied to Fnl^flox/+^ and Fnl^Δ/-^ MEFs for each antibody. Primary antibodies were detected with anti-rabbit secondary antibodies conjugated with AlexaFluor-647 and imaged using identical settings. Signal intensity for stained Fnl^Δ/-^ cells was increased when making these panels to show background fluorescence. **B. 297.1 antibody reacts with bovine Fn1.** CM – complete medium as in Table M1. CCM – conditioned complete medium. CCM was prepared by incubating 2x10^5^ wild-type MEFs with 2 ml of CM for 48 hours in one well of a 6 well plate. Equal amount of medium was loaded in wells of Novex™ WedgeWell™ 4 to 12% Tris-Glycine protein gel. Arrow points at Fn1 band.

